# Gut microbiota–mediated lipid accumulation as a driver of evolutionary adaptation to blue light toxicity in *Drosophila*

**DOI:** 10.1101/2024.08.22.608892

**Authors:** Yuta Takada, Toshiharu Ichinose, Naoyuki Fuse, Kokoro Saito, Wakako Ikeda-Ohtsubo, Hiromu Tanimoto, Masatoshi Hori

**Author notes:** Correspondence to Yuta Takada or Masatoshi Hori.

## Abstract

Human agriculture has always raced against insect adaptation, requiring updated pest control methods and knowledge of evolutionary processes. Excessive exposure to blue light (BL) kills a wide range of insect species and has attracted attention as an alternative to chemical pesticides. Here, to understand how insects adapt to BL toxicity, we investigated evolutionary responses to BL toxicity in *Drosophila melanogaster* over 70 generations using laboratory selection experiments. The selected line exhibited an obese phenotype accompanied by midgut elongation, with BL tolerance dependent on gut microbiota– mediated lipid accumulation. Whole-genome and transcriptome analyses consistently highlighted interactions between the microbiota and host lipid metabolism–related genes. Remarkably, manipulating genes associated with lipid accumulation conferred BL tolerance even in the absence of selection. We suggest that the acquisition of BL tolerance occurs through ’adaptive obesity’. Our study introduces a mechanism of evolutionary adaptation of insects against BL-based selective pressure by maximising the benefits from the gut microbiota via midgut elongation.

**Graphical abstract:** 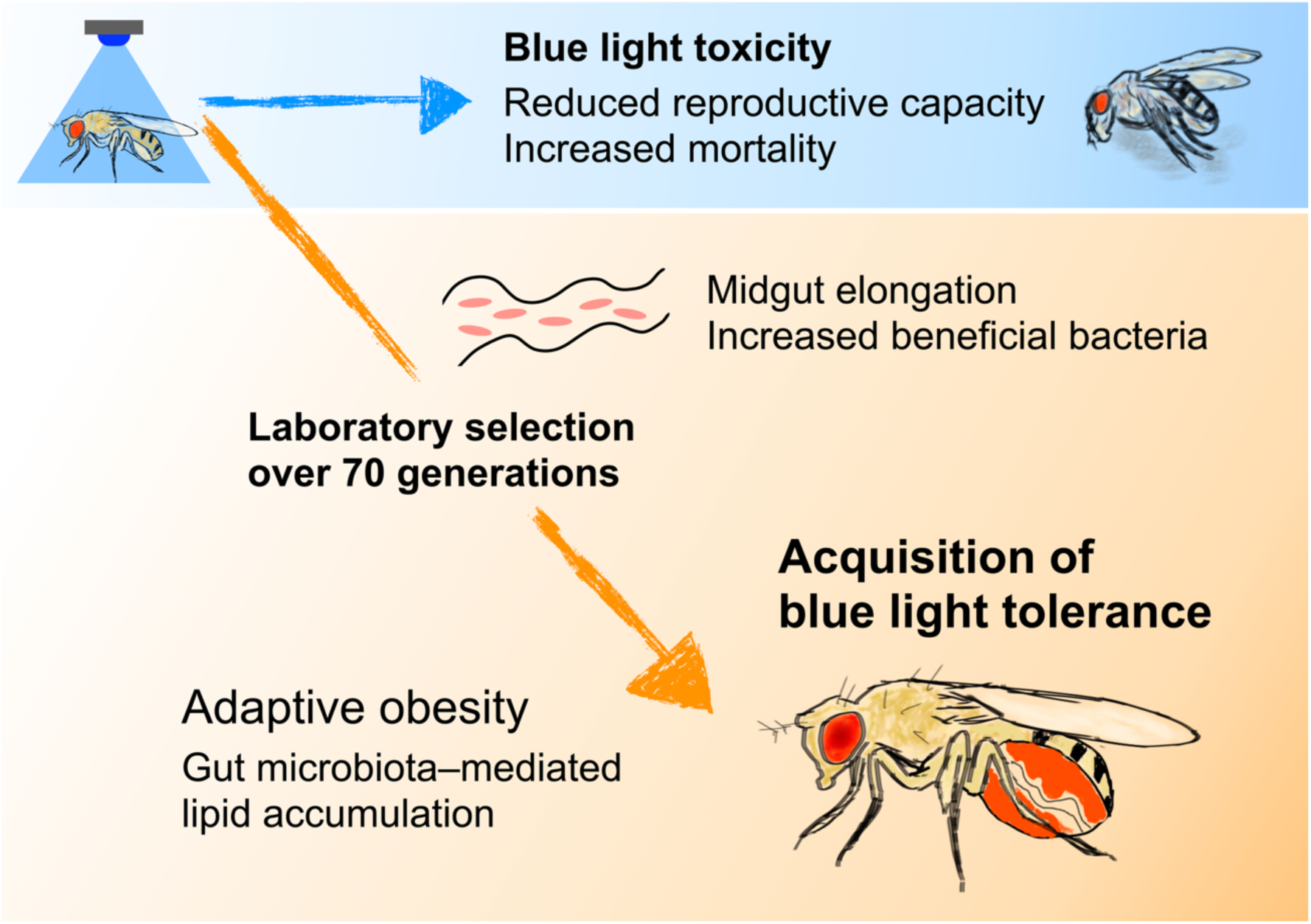

**Brief summary:** Laboratory selection for tolerance to blue light toxicity in *Drosophila* revealed that gut microbiota–mediated obesity, marked by midgut elongation and increased beneficial bacteria, is a key evolutionary adaptation.

## Introduction

One of the most challenging issues in pest control is the rapid evolution of insects. Traditional chemical pesticides have led to the development of resistance, environmental contamination and unintended effects on non-target organisms, highlighting the need for alternative approaches^1,2^.

Blue light (BL) is within the 400–500 nm range, is abundant in sunlight, and is crucial for insect physiology and ecology because it influences processes such as visual perception, circadian rhythms, and photoreactivation (Supplementary Fig. 1)^3–6^. However, excessive exposure to BL is lethal in a variety of insect species^7^. We propose the use of BL as a pest control method and have demonstrated that exposure to BL at intensities below 40% of the BL content of sunlight for several days can be lethal to fruit fly *Drosophila melanogaster* (hereinafter referred to as *Drosophila*), mosquito *Culex pipiens* form *molestus,* flour beetle *Tribolium confusum,* and leaf beetle *Galerucella grisescens*^7–9^. Notably, in *Drosophila*, *C. pipiens* f. *molestus*, and *G. grisescens*, BL around 420, 440, and 465 nm has been found to be more lethal than shorter wavelength 375–405 nm^7–9^. In *Drosophila*, BL around 465 nm is particularly toxic to both pupae and adults, with males being more sensitive than females^9^. These results suggest that the sensitivity of BL toxicity cannot be explained by photon energy alone. The BL toxicity is thought to be associated with reactive oxygen species (ROS) generated by mitochondria and other cellular components, leading to oxidative damage^9–14^. This damage can impair fitness and act as a selective pressure in insects, yet tolerance and evolutionary adaptation to this stressor remain largely unexplored. Achieving the societal implementation of chemical-free, BL-based pest control requires the gathering of evolutionary insights.

Laboratory selection has contributed to understanding the evolutionary adaptations of organisms to environmental stressors^15,16^. *Drosophila* is well-suited for laboratory selection experiments due to its short life cycle and the extensive genetic and physiological knowledge available^17^. Importantly, these experiments often reveal both evolutionary responses and trade-offs that were not anticipated at the outset^18^. Observing evolutionary processes in the laboratory and identifying the phenotypic, genomic and epigenomic factors that lead to tolerance in *Drosophila* are crucial to understanding the mechanisms of evolution in insects. In this study, by selecting flies that are tolerant to excessive BL exposure, we established *Drosophila* strain with enhanced BL tolerance. The characterisation of these selected line (SL) flies revealed that obesity is a key adaptive trait for BL tolerance. Moreover, we found that parental BL exposure and selection under BL toxicity induce midgut elongation and that increased gut bacterial load is critical for acquired BL tolerance. This study highlights that maximising the benefits of the gut microbiota has been key to evolutionary adaptation in the host.

## Results

### *Drosophila* evolved BL tolerance through laboratory selection

Reproductive capacity is one of the most important determinants of fitness. We investigated the effects on ovarian development to determine whether damage caused by BL toxicity acts as a selective pressure. Older flies are known to be more sensitive to BL toxicity^13^, but our results showed that even very young flies, such as those on the first day after eclosion with underdeveloped ovaries, were particularly susceptible (Supplementary Fig. 2a and Supplementary Data 1)^19^. BL toxicity consistently suppressed ovarian development at all stages between days 1 and 10 post eclosion (Supplementary Fig. 2a). Even after a recovery period of up to 7 days post-BL exposure, ovarian recovery remained minimal (Supplementary Fig. 2b and Supplementary Data 1). These results highlight that BL toxicity has a dramatic effect on fitness.

First, we generated a *Drosophila* strain with BL tolerance through laboratory selection. In brief, we derived two experimental strains from the same population: one maintained without selective pressure and the other reared under BL toxicity-driven selection (Fig. 1 a). This procedure was replicated in three independent lines for each strain and each line was repeated for over 70 generations (Fig. 1 a). For laboratory selection, a Canton Special (Canton-S) strain, which is a long-inbred strain of the common wild type, was used as the ancestral population (for details, see "Laboratory selection for BL tolerance" in Methods)^20^. Over the generations, 1–5 day-old flies were exposed for 3 days to BL at around 465 nm, a wavelength highly toxic to *Drosophila* adults, at a photon flux density at which approximately 50% of individuals exhibited the “low impact” phenotype, which was defined as being capable of climbing after a 2-day recovery (Fig. 2b and Supplementary Fig. 1 and Supplementary Fig. 3a)^9^. We then selected the low impact flies and let them breed to develop the SL lines (Fig. 1b). The unselected control line (UCL), maintained without BL irradiation, served as a negative control for comparative analyses with the SL. In the negative control group, 3 days of BL irradiation were replaced by darkness and UCL parents were randomly chosen (Fig. 1b and Supplementary Fig. 3a). Although not selected, a subset of UCL flies in each generation was exposed to BL under the same conditions as the SL to serve as a positive control group (Supplementary Fig. 3a). Across generations, no strain-specific differences in climbing ability were observed without BL irradiation (Supplementary Fig. 3b and Supplementary Data 1). In SL, both male and female flies showed greater climbing ability after BL irradiation following a single selection event (Supplementary Fig 3a). In contrast, fecundity in SL remained suppressed by BL toxicity even after more than 10 generations of selection (Fig. 2a). After around 20 generations, SL flies began to show improved adaptability, with fecundity maintained at similar levels to flies in the non-irradiated UCL (Fig. 2a–c, Supplementary Data 2 and Supplementary Data 3). Ovarian development was affected little by BL exposure in the SL flies but severely in the UCL flies (Fig. 2b and Supplementary Data 3). Moreover, fecundity in SL flies was slightly increased in response to BL irradiation, whereas that in UCL flies was reduced (Fig. 2c and Supplementary Data 3). Additionally, SL flies survived longer than UCL flies under continuous BL irradiation (Fig. 2d and Supplementary Data 3). After 73 generations of selection, the lifespan was further extended, with disparity from the UCL increasing as light intensity decreased (Fig. 2d, e, Supplementary Fig. 4 and Supplementary Data 3). These results collectively suggest that we have successfully established a strain of *Drosophila* with enhanced BL tolerance.

**Fig 1:**
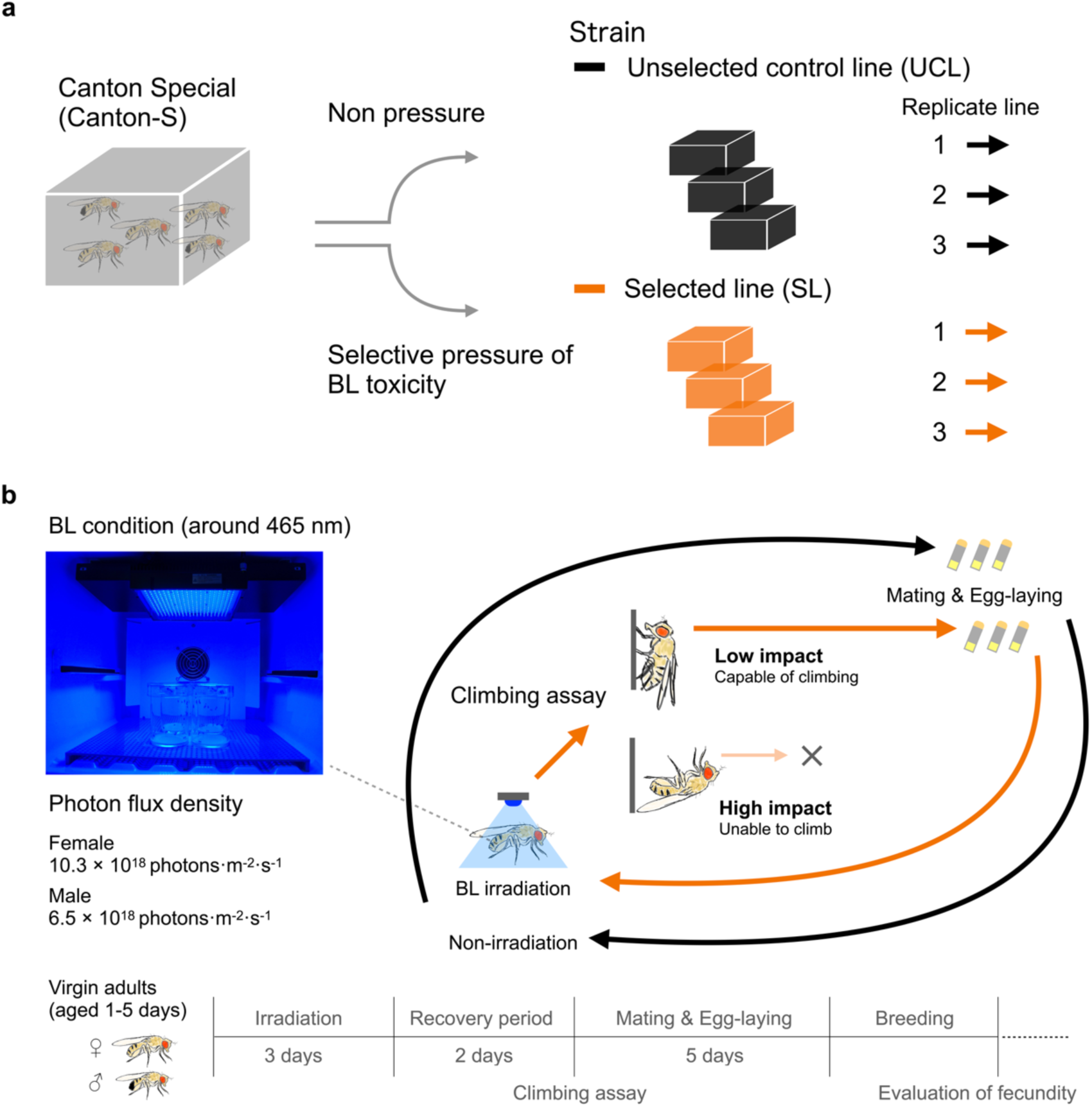
Experimental design of laboratory selection using blue light (BL) toxicity. **a**, Overview of the ancestral population and the laboratory selection experiment. We established two strains from the Canton Special (Canton-S) strain: a selected line subjected to BL toxicity pressure at the adult stage in each generation, and an unselected control line without selection pressure. Each strain was maintained with three replicate lines. **b**, Detail of the selection experiment. Flies were BL-irradiated for 3 days, recovered for 2 days, and categorized in a climbing assay into those capable of climbing ("low impact") and those unable to climb ("high impact"). In SL, parents selected from low impact flies. Flies were allowed 5 days for mating and egg-laying. UCL parents were randomly chosen from the negative control group and reared under the same conditions as the SL flies.

**Fig 2:**
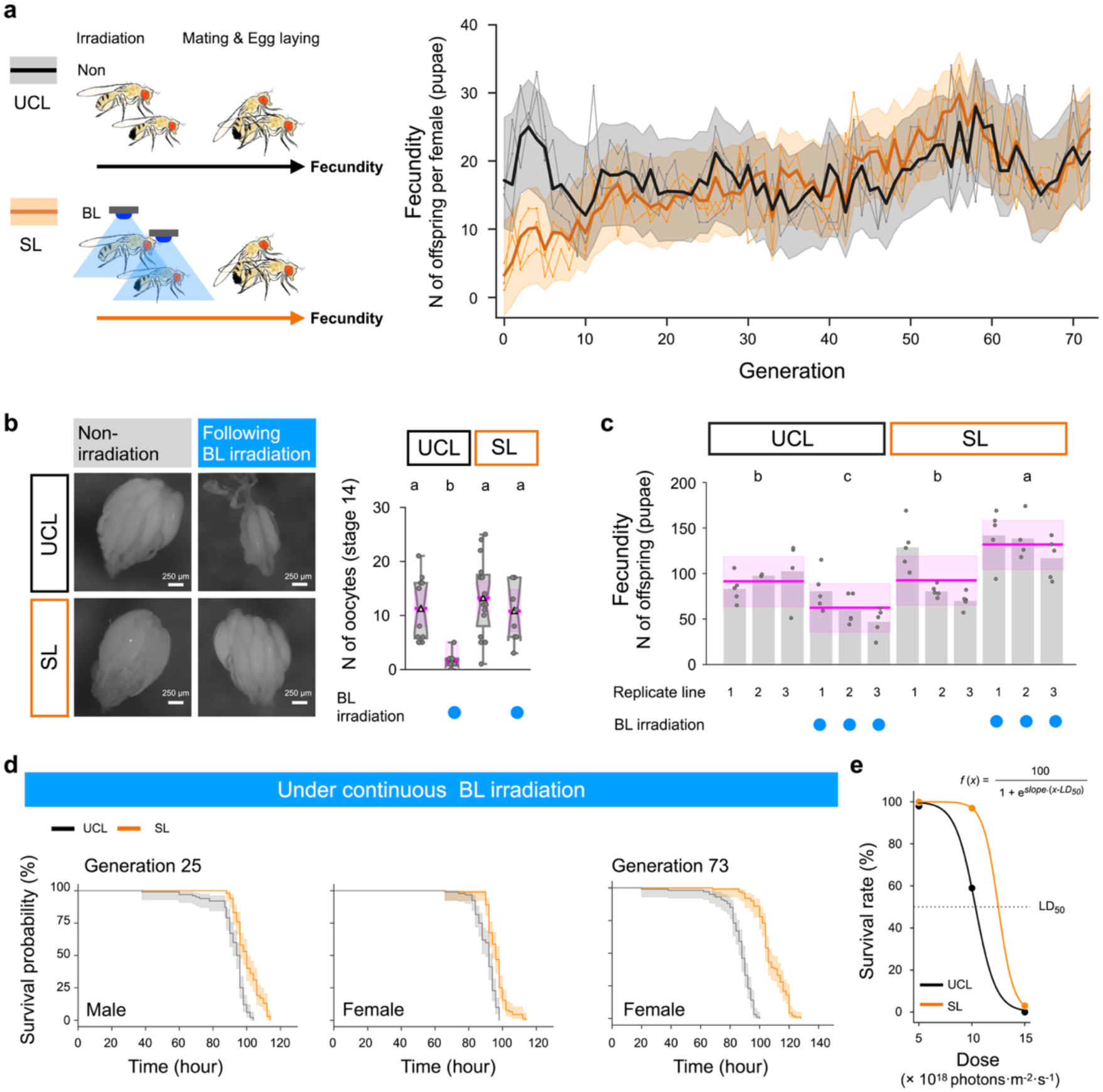
Establishment of *Drosophila* lines with BL tolerance evolved by laboratory selection. **a**, Changes in fecundity across generations. UCL represents fecundity under non-irradiated conditions, while SL represents fecundity after BL irradiation. Fecundity was measured as the number of offspring per female (pupae). *n* = 3. The bold lines and shaded areas show the posterior median and Bayesian 95% credible interval (CI) for each strain. These were estimated using a state space model with Markov chain Monte Carlo (MCMC) sampling. The grey and orange lines represent the observed values for each replicate (UCL and SL, respectively). **b**, Ovarian development differences after 3 days of BL irradiation followed by 3 days of recovery. Number of stage 14 oocytes after a 3-day recovery period. *n* = 12, 10, 15, 9. Bayes factor (BF10) = 50.74. (Replicate lines 2, generation 25). Box plots show the median (centre line), the interquartile range (box), the full range of the data (from minimum to maximum), the notch (which visualises the variability of the data around the median) and the mean (triangle). **c**, Fecundity measured as the number of offspring (pupae). *n* = 11, 14, 15, 14. BF10 > 100, generation 38. **d**, Lifespan under continuous BL irradiation. Male *n* = 100. BF10 >100. Estimates: UCL 90.62 h (89.38–91.82), SL 96.35 h (95.16–97.52), *p*MCMC < 0.001. Female *n* = 92. BF10 = 0.72. Estimates: UCL 92.09 h (90.38–93.78), SL 100.39 h (98.74–102.13), *p*MCMC < 0.001, replicate lines 1, generation 25. Female *n* = 100. BF10 > 100. Estimates: UCL 86.16 h (84.05–88.27), SL 107.18 h (105.09–109.28),*p*MCMC < 0.001, replicate lines 1, generation 73. **e**, Dose-response curve for females exposed to BL. The lethal dose (LD50) was estimated by logistic regression: 10.35 × 10¹⁸ photons-m-²-s-¹ for UCL and 12.50 × 10¹⁸ photons-m-²-s-¹ for SL. See Supplementary Fig. 4 for details of lifespan at 5 × 10¹⁸ photons-m-²-s-¹ and 15 × 10¹⁸ photons-m-²-s-¹. Statistical analyses for panels **b**–**d** used a Hierarchical Bayesian Model (HBM) with MCMC methods, posterior median, and Bayesian 95% CI. The magenta, bold lines and shaded areas show the posterior median and the Bayesian 95% CI. In **b** and **c**, different letters indicate significant differences at *p*MCMC < 0.05. See Supplementary Data 2 and Supplementary Data 3 for details of all results, including statistical models.

### SL flies exhibited obese phenotype, lipid accumulation, and oxidative stress resistance

Next, we examined the phenotypic characteristics of the SL flies in non-irradiated conditions. We noticed that they exhibited an obese phenotype characterised by abdomen bloating, increased body weight, and well-developed fat bodies with markedly increased lipid droplets (Fig. 3a, b and Fig. 4a–b^ii^ and Supplementary Fig. 5a and Supplementary Data 4). They also showed an elongated midgut and shortened wings (Fig. 4c and Supplementary Fig. 5b and Supplementary Data 4). Additionally, we observed a slightly low number of oocytes in the SL flies (Supplementary Fig. 5c and Supplementary Data 4) and a strong inter-individual correlation between oocyte number and body weight in the UCL flies (Supplementary Fig. 5d and Supplementary Data 4). SL flies maintained higher body weight than UCL flies despite similar food consumption levels (Fig. 4d and Supplementary Data 4). Taken together, these results show that the increased body weight of SL flies was not due to larger ovaries or higher food intake but may instead be associated with increased lipid reserves and elongated midguts. SL flies were rich in major lipids, including fatty acids (fatty acid methyl esters, FAMEs) and triacylglycerol (TAG), and were starvation resistant, providing evidence of lipid accumulation (Fig. 4e–g and Supplementary Data 4)^21,22^. Consistent with previous reports that lipid droplets and their constituent TAG may protect against oxidative damage^23–25^, resistance to oxidative stress was higher in SL flies than in UCL flies (Fig. 4h and Supplementary Data 4). We did not find any physical traits associated with light reflection by cellular melanin, despite previous reports suggesting these as adaptive traits to phototoxicity (Supplementary Fig. 6)^26^. These results suggest that BL tolerance in SL flies is acquired physiologically through obesity, rather than through physical traits.

**Fig 3:**
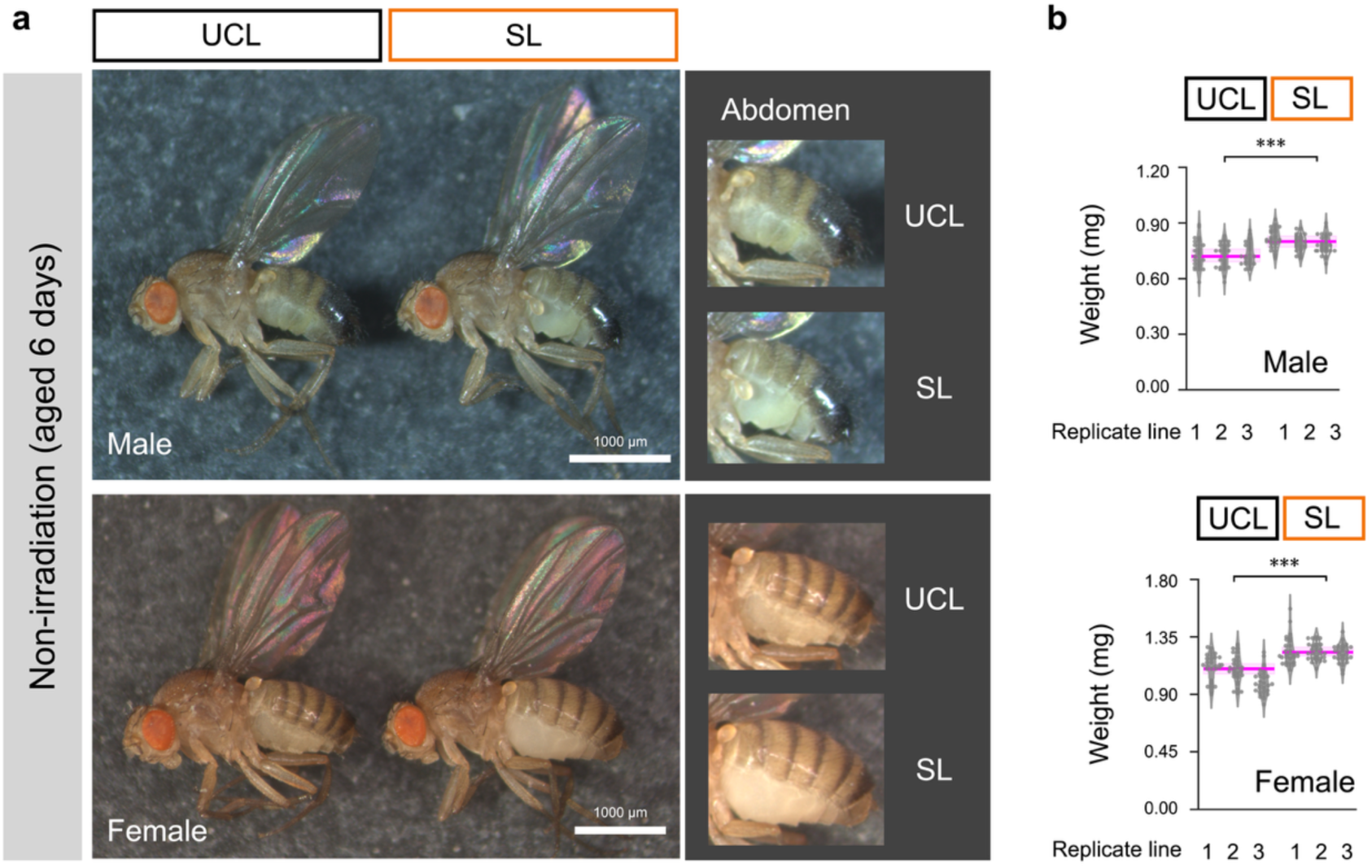
Selected line (SL) flies characterised by enlarged abdomen and heavier body weight in both sexes. **a,** Phenotype of adult virgin male and virgin female aged 6 days. The zoomed view of the abdomen is brightened with maximum exposure. **b**, Body weight, *n* = 90, BF10 >100, generation 39. Statistical analysis for panel **b** was performed using a HBM with MCMC methods, posterior median, and Bayesian 95% CI. The magenta, bold lines and shaded areas show the posterior median and the Bayesian 95% CI. Asterisks indicate significance levels: *** *p*MCMC < 0.001. See Supplementary Data 4 for details of all results, including statistical models.

**Fig 4:**
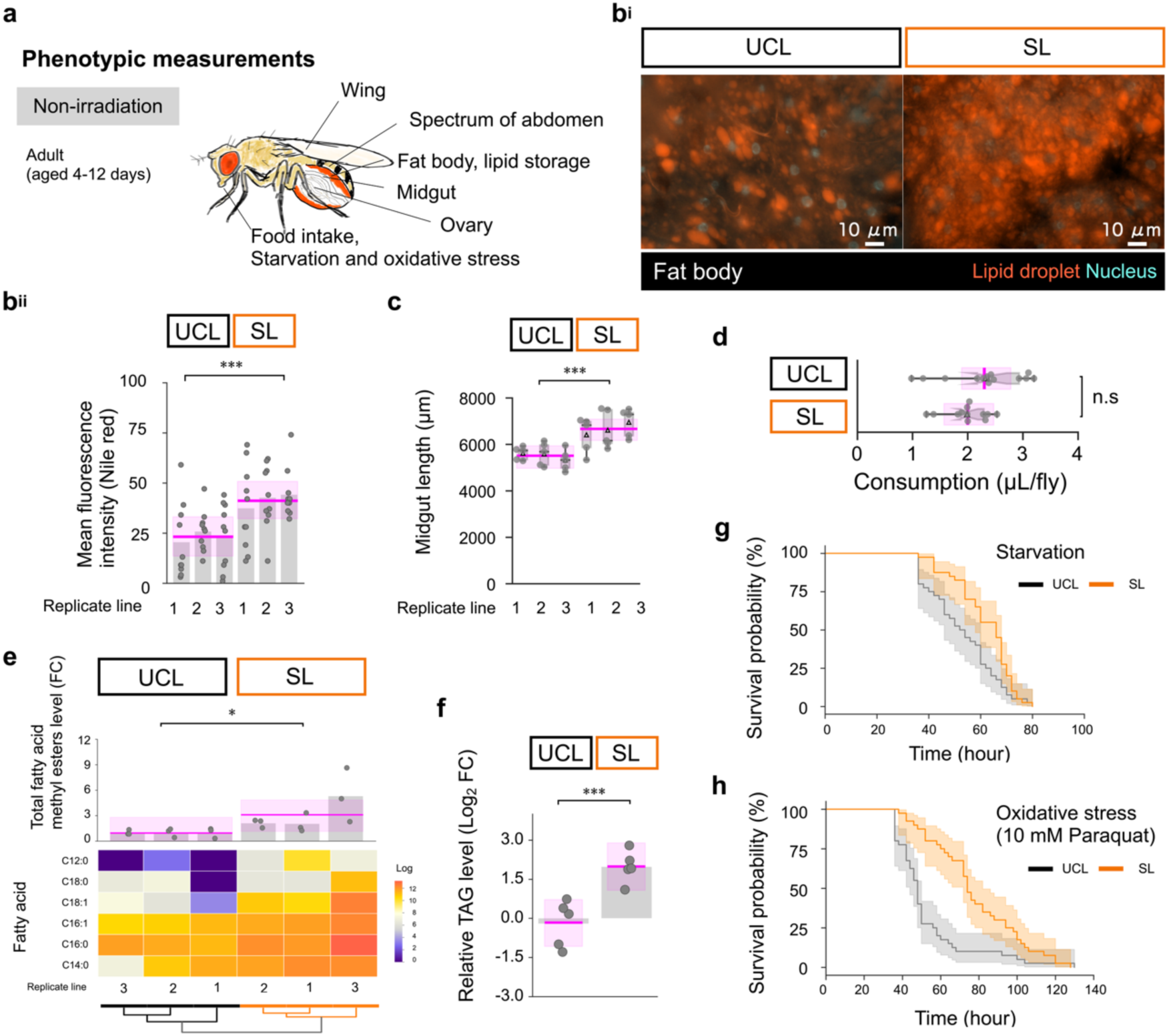
SL flies exhibited obese phenotype, lipid accumulation, and oxidative stress resistance. **a**, Phenotypic measurements of virgin adult females (aged 4-12 days) under non-irradiated conditions (**b**-**h**). Age matching was performed between strains at the time of measurement. See Supplementary Fig. 5 and Supplementary Fig. 6 for details of wing, ovary and abdomen spectrum. **b^i^**, Fluorescence imaging of the abdominal fat body and lipid droplets (lipid droplets: Nile red; nuclei: NucBlue). **b^ii^**, Mean fluorescence intensity from Nile red staining, *n* = 30, BF10 >100, generation 35. **c**, Midgut length, *n* = 15, BF10 >100, generation 45. **d**, Food consumption, *n* = 13, 12, BF10 = 1.85, replicate line 1, generation 52. **e**, Gas chromatography mass spectrometry analyses of fatty acid methyl esters (FAMEs), total FAMEs, *n* = 9, BF10 = 1.69, generation 30. **f**, Relative triacylglycerol (TAG) level, *n* = 5, BF10 =15.14, replicate line 1, generation 57. **g**, Starvation resistance, Kaplan– Meier survival curve, *n* = 40, BF10 = 4.49. Estimates: UCL 53.04 h (49.20–56.89), SL 61.91 h (58.09–65.43), *p*MCMC < 0.01, replicate line 2, generation 43. **h**, Oxidative stress resistance (10 mM paraquat), Kaplan–Meier survival curve, *n* = 40, BF10 >100. Estimates: UCL 53.07 h (45.90–59.03), SL 77.94 h (71.19–84.08), *p*MCMC < 0.001, replicate line 3, generation 43. Statistical analyses for panels **b** and **d**–**h** were performed using a HBM with MCMC methods, posterior median, and Bayesian 95% CI. The magenta, bold lines and shaded areas show the posterior median and the Bayesian 95% CI. Asterisks indicate significance levels: * *p*MCMC < 0.05, *** *p*MCMC < 0.001, n.s = not significant. FC, fold change. Box plots show the median (centre line), the interquartile range (box), the full range of the data (from minimum to maximum), the notch (which visualises the variability of the data around the median) and the mean (triangle). See Supplementary Data 4 for details of all results, including statistical models.

### Increased bacterial abundance in SL flies

Gut microbiota–derived acetate serves as a potential substrate for de novo lipogenesis in the host via acetyl-CoA^27,28^. Considering the elongated midgut and accumulated lipids in SL flies, we compared the gut microbiota profiles and amounts in SL and UCL flies. In both strains, 16S rRNA gene sequencing of whole-body DNA revealed that a single species of *Acetobacter* (*A. persici)*, a common gut bacteria in *Drosophila*, dominated (>99%) the microbiota (Supplementary Fig. 7a and Supplementary Data 5)^29–31^. *Wolbachia* was not detected in our microbiota analyses (Supplementary Data 5). *A. persici* has also been detected in wild flies, and several studies have reported that in some cases more than 90% of the microbiota consisted of this or other *Acetobacter* species^30–34^. Relatedly, several studies have reported that *Acetobacter* can dominate the gut microbiota in natural environments^35,36^. Notably, bacterial counts were higher in SL flies than in UCL flies, and a mild positive relationship between body weight and bacterial abundance was observed, with no change in microbiota profiles (Fig. 5a^i^–a^ii^, Supplementary Fig. 7bi, b^ii^, Supplementary Fig. 8, Supplementary Data 5 and Supplementary Data 6). All randomly sequenced colonies were identified as *A. persici* (Supplementary Data 6). These results consistently show that while both strains colonise the same species, SL has the higher bacterial abundance.

**Fig 5:**
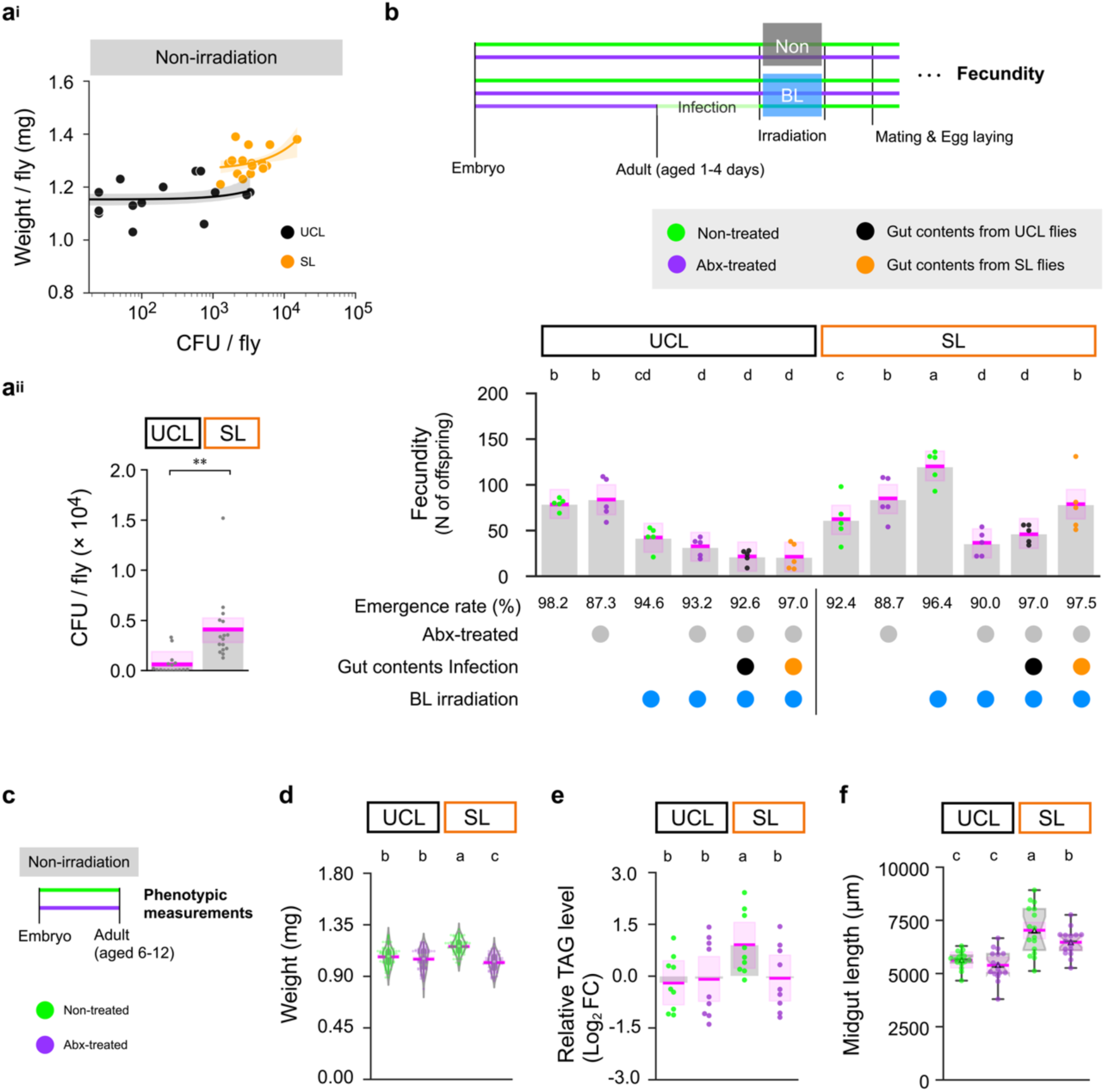
High bacterial abundance is the key to BL tolerance and lipid accumulation in SL flies. **a^i^,** Relating body weight to bacterial abundance. *n* = 16, UCL; *y* = 0.00 × *x* +1.15, *p* = 0.54, SL; *y* = 0.00 × *x* +1.27, *p* = 0.06. Bacterial measurements from virgin adult females (aged 6-12 days) under non-irradiated conditions. After measuring body weight, homogenised samples from four flies were diluted 100-fold and cultured on MRS plates for 72 hours to quantify colony-forming units (CFU). The variance represents the 80% confidence interval. See Supplementary Fig. 7 for microbiota analysis. **a^ii^,** Bacterial abundance. *n* = 16, BF10 = 11.17. Asterisks indicate significance: ** *p*MCMC < 0.01; replicate line 1, generation 48. CFU from same dataset as Fig. 5a. **b,** Impact of antibiotic mixture (Abx) treatment and gut content infection on fecundity under BL toxicity. Fecundity measured as the number of offspring (adults), *n* = 5, BF10 >100; replicate line 1, generation 47. **c**, Phenotypic effects of Abx treatment (**d**–**f**). Abx was administered continuously from the embryo to adult, resulting in a significant reduction in the adult microbiota. The efficacy of Abx treatment was confirmed by regular plating of adults on MRS plates, where CFU counts were zero in all observations. See Supplementary Fig. 9a for details. **d**, Body weight, *n* = 100, BF10 >100; replicate line 1, generation 55. **e**, Relative TAG level, *n* = 10, BF10 = 1.47; replicate line 1, generation 55. **f,** Midgut length, *n* = 16, BF10 >100; replicate line 1, generation 48. Box plots show the median (centre line), the interquartile range (box), the full range of the data (from minimum to maximum), the notch (which visualises the variability of the data around the median) and the mean (triangle). Statistical analyses for panels **a^ii^**–**f** were performed using a HBM with MCMC methods, posterior median, and Bayesian 95% CI. The magenta, bold lines and shaded areas show the posterior median and the Bayesian 95% CI. Different letters indicate significant differences at *p*MCMC < 0.05. See Supplementary Data 6 for details of all results, including statistical models.

### Gut microbiota promotes BL tolerance and lipid accumulation in SL flies

Acetic acid bacteria, including *Acetobacter*, are known to influence fecundity and TAG levels in *Drosophila*^37–45^. In addition, both bacterial abundance and body weight were drastically reduced in SL flies following BL irradiation, suggesting that the sensitivity of these flies to BL toxicity may be related to the gut microbiota (Supplementary Fig. 8). To test the hypothesis that gut microbiota play a role in shaping the phenotype of SL flies, we used an antibiotic mixture (Abx) that depletes the bacterial abundance (Supplementary Fig. 9a). Interestingly, we found that BL tolerance in the SL flies, as measured by fecundity, was totally abolished by Abx treatment, suggesting the critical role of gut microbiota (Fig. 5b and Supplementary Data 6). Feeding the Abx-treated SL flies with the gut contents of SL flies rescued these phenotypes, whereas feeding with those of UCL flies barely did (Supplementary Fig. 9b–d, Fig. 5b, Supplementary Data 6 and Supplementary Data 7). Consistently, Abx treatment significantly reduced the body weight and the TAG level selectively in the SL flies (Fig. 5c–e and Supplementary Data 6). This result highlights the dependence of SL flies’ body weight and lipid accumulation on their gut microbiota.

The midgut of Abx-treated SL flies was longer than that of UCL flies regardless of treatment, and it was even longer in non-treated SL flies (Fig. 5f and Supplementary Data 7). A positive correlation between body weight and midgut length in non-treated SL flies (Supplementary Fig. 10 and Supplementary Data 7) suggested that gut microbiota may contribute to midgut elongation. Furthermore, enhanced oxidative stress resistance was observed only in non-treated SL flies (Supplementary Fig. 11a, b and Supplementary Data 7). These observations align with previous reports that gut microbiota is linked to gut morphology and oxidative stress resistance^29,46,47^. Overall, these results suggest that the characteristic phenotype in SL flies is shaped by their gut microbiota.

### Parental exposure to BL affects gut microbiota–associated traits in progeny

We investigated why only the gut contents from SL flies could rescue BL tolerance in infection experiments, despite the dominance of the same gut bacterial species across strains (Supplementary Fig. 7a and Fig. 5b). We noticed that the flies used for gut contents had different parental experiences depending on whether they belonged to the UCL or SL. Additionally, parental exposure to BL has been reported to alter the transcriptome of the progeny^48^. Thus, we hypothesised that parental exposure to BL irradiation might influence gut microbiota–associated traits in the progeny, leading to quantitative or qualitative changes in gut microbiota. We exposed adult males and females from both strains to the same BL toxicity used in the selection experiment for 3 days, followed by a 3-day recovery period. We then randomly picked flies to be the parent, producing progeny that were influenced by the parental experience (Fig. 6a). First, we examined the body weight and BL tolerance in Abx-treated SL flies that were fed with gut contents from flies whose parents had or had not received BL irradiation. Abx-treated SL flies fed with gut contents from SL flies whose parents had experienced BL irradiation showed the greatest increase in body weight (Supplementary Fig. 12a^i^–b and Supplementary Data 8). However, flies fed with gut contents from *A. persici* in different cocktails, each standardized in concentration, exhibited similar body weight increases (Supplementary Fig. 12c and Supplementary Data 8). This suggests that quantitative rather than qualitative changes in gut microbiota due to parental BL irradiation are responsible for the increases in host body weight. Furthermore, gut contents from UCL flies whose parents had experienced BL irradiation also rescued the phenotypes of Abx-treated SL flies, restoring fecundity to the level conferred by gut contents from SL flies (Supplementary Fig. 13a^i^–b and Supplementary Data 8). These results indicate that parental BL irradiation affects the progeny’s bacterial abundance. Consistent with this idea, parental BL irradiation increased the progeny’s body weight, bacterial abundance, and midgut length in both UCL and SL flies (Supplementary Fig. 14, Fig. 6b, c and Supplementary Data 8). Notably, even without parental BL irradiation, the midgut of SL flies was longer than that of non-irradiated UCL flies, suggesting that this inherited trait had been conserved during the laboratory selection of SL flies (Fig. 6c). In both strains, parental BL irradiation induced midgut elongation even after antibiotic treatment, suggesting that in *Drosophila* this is a common response dependent on parental experience. The increased abundance of beneficial bacteria under BL toxicity may improve fitness, highlighting the dependence of BL tolerance in SL flies on gut microbiota.

**Fig 6:**
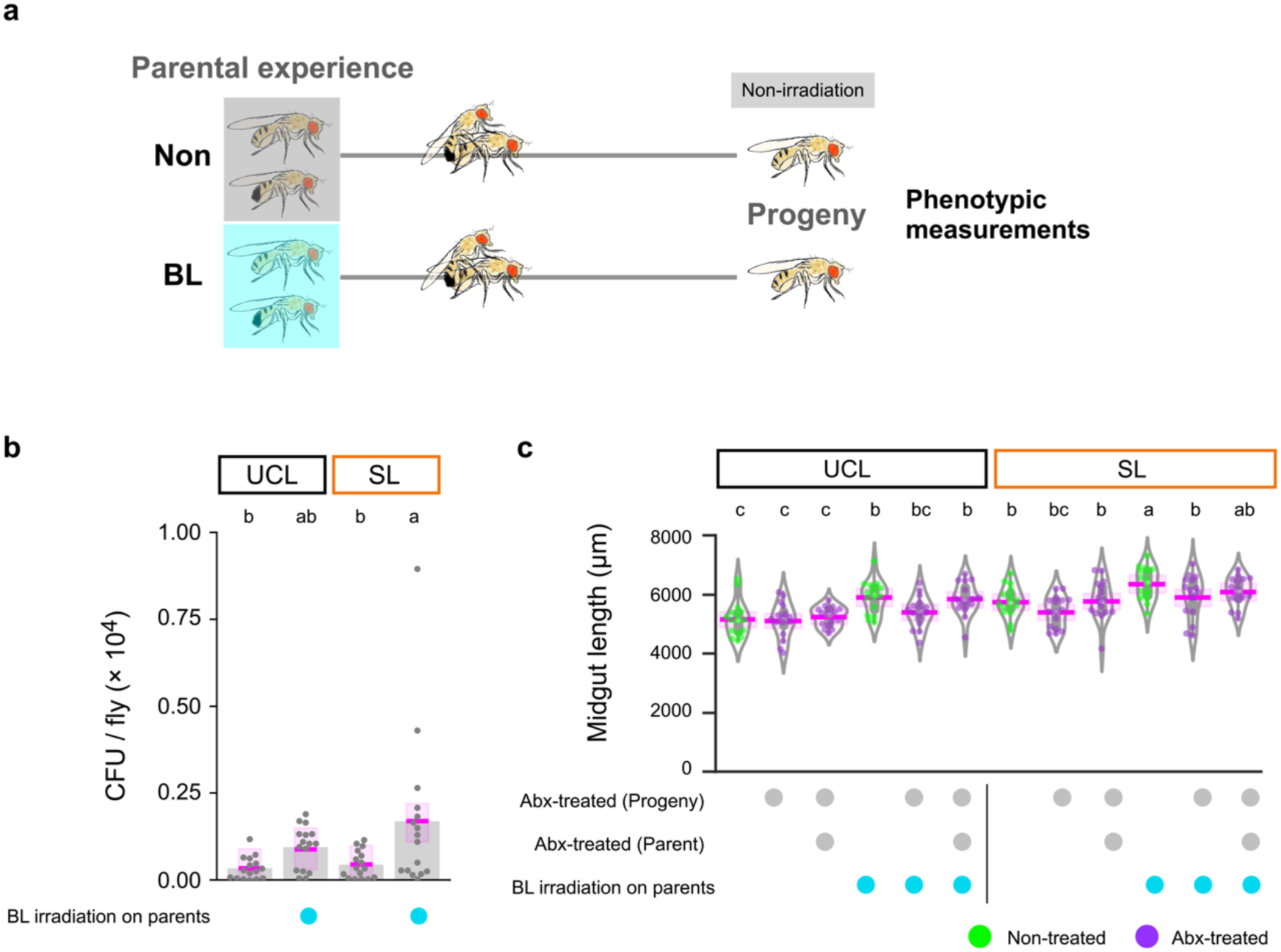
Parental experience of BL irradiation promotes midgut elongation and increased bacterial abundance in progeny. **a**, The effect of parental exposure to BL toxicity on offspring traits. Offspring with different parental experiences were generated. **b**, Bacterial abundance. *n* = 16. BF10 = 8.93; replicate line 1, generation 53. **c**, Midgut length. *n* = 16, BF10 >100; replicate line 1, generation 54. Statistical analyses for panels **b** and **c** were performed using HBM with MCMC methods, posterior median, and Bayesian 95% CI. The magenta, bold lines and shaded areas show the posterior median and the Bayesian 95% CI. Different letters indicate significant differences at *p*MCMC < 0.05. For details, see Supplementary Data 8 for all results, including statistical models.

### Host genome and transcriptomic profiles explain the characteristic phenotype of SL flies

To investigate how the SL flies develop their phenotypes through interactions with the gut microbiota, we first performed whole-genome resequencing aimed at detecting genetic characteristics unique to SL. The first approach was to explore genes that show strain-specific variation to understand the genetic variation of SL. We identified 1,917 genes with SL-specific variation above 5% for the full sequence length of each gene (Fig. 7a, Supplementary Data 9 and Supplementary Data 10). Kyoto Encyclopedia of Genes and Genomes (KEGG) enrichment analysis of these variations revealed their association with detoxification and antioxidant response such as *Glutathione S transferase* (e.g. *GstD4*, *GstD5, GstD6, GstD7, GstD10, GstE1, GstE2, GstE5, GstE6, GstE8, GstE11, GstE14, GstO2, GstT3*) (Fig. 7b and Supplementary Data 10)^49^. Given the role of oxidative stress in BL toxicity^9,11,13^, the variation in these detoxification enzymes suggests functional changes and the effect of selective pressures. The original of Canton-S strain is isogenic in the 1920- 1930s, although it has already accumulated variants due to long laboratory breeding (note that the sequencing data is from the Canton-S strain obtained from the Bloomington *Drosophila* Stock Center (Supplementary Fig. 15a))^20,50^. Even in UCL, the detection of 694,867 total sites with heterozygous and homozygous variants across the three replicate lines supports the possibility that multiple standing variants were present (Supplementary Fig 15b–d). We analysed strain-specific homozygous variants to further elucidate the genetic basis of SL. Homozygous variants shared among the three replicate lines in SL and absent in UCL and in the Canton-S strain were identified (Supplementary Fig. 16). SL-specific SNPs and indels were detected in 658 and 2,625 genes, respectively, giving a total of 2,970 genes, and the majority of alterations were located outside the coding sequences (CDS) (Fig. 7c and Supplementary Data 11). KEGG enrichment analysis identified the Notch signaling and the Hippo signaling pathway as enriched pathways (Fig. 7d), consistent with their known function in organ size regulation^51–53^. In particular, the presence of CDS variants in genes that regulate or interact with the Hippo signaling pathway (*Par-1* (missense mutation), *misshapen* (frameshift mutation)), together with the elongated midgut observed in Abx-treated SL flies, suggests that midgut elongation was potentially influenced by these variants (Supplementary Data 11, Fig. 5f and Fig. 6c)^52,54,55^. Furthermore, the Toll and Imd pathways of the immune system were identified as enriched pathways of 4,612 genes with SL genetic characteristics (Fig. 7e and Supplementary Data 12)^56,57^. The genetic features of SL suggest that the acquisition of gut microbiota-dependent phenotypes may have been facilitated by midgut elongation and immune system modifications.

**Fig 7:**
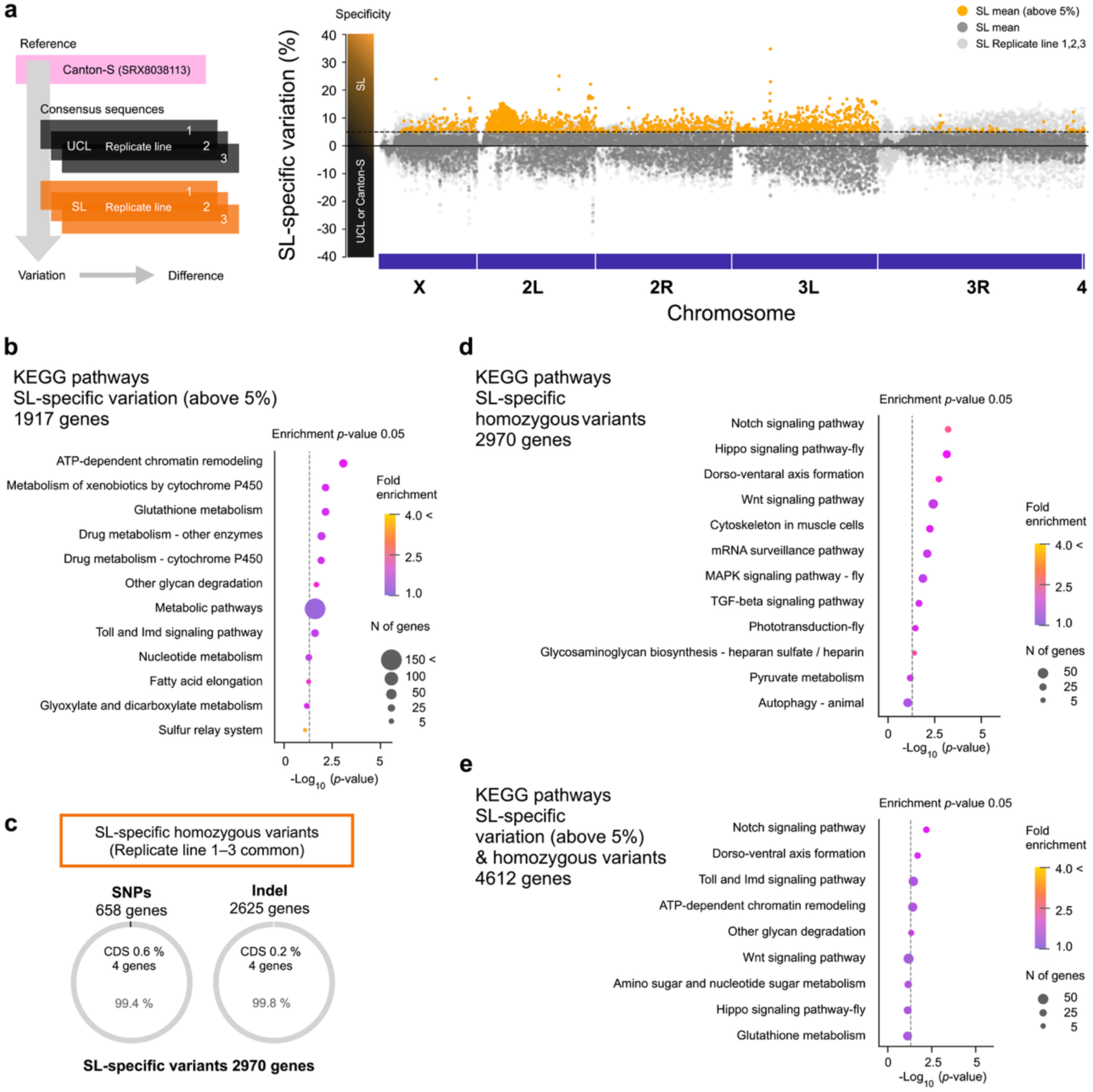
Genetic characteristics unique to SL are enriched in detoxification, antioxidant response, Hippo signaling pathway and Toll and Imd signaling pathway. **a**, Manhattan plot of variation for each gene from whole genome resequencing analysis; generation 44. Variation of genes are referenced to Canton-S strain (SRX8038113). SL- specific variation is defined as the difference between the UCL mean variation and the SL variation per replicate line and mean greater than 5%. Each SL replicate line is shown in light grey, the SL mean is shown in dark grey and SL means with more than 5% variation are highlighted in yellow. Negative values indicate higher variation in Canton-S or UCL. A value of zero indicates no difference between UCL and SL. **b**, Enrichment analysis of SL- specific variation (above 5%) using Kyoto Encyclopedia of Genes and Genomes (KEGG). **c**, Identification of SL-specific common homozygous variants. CDS, coding sequence. **d**, Enrichment analysis of SL-specific variants using KEGG. **e**, Enrichment analysis of SL- specific variation (above 5%) and variants using KEGG. For details, see Supplementary Data 9–12.

Because variations located outside the CDS are generally thought to influence gene expression and because gut microbiota may alter host phenotypes through changes in the host transcriptome^37,58,59^, we performed transcriptome analysis on the whole body of non-irradiated UCL and SL flies, with or without the Abx treatment. We identified 1,242 differentially expressed genes (DEGs), of which 1,192 (96%) were specific to non-treated SL flies (Supplementary Fig. 17a, b, Fig. 8a and Supplementary Data 13). KEGG enrichment analysis of the 1192 DEGs revealed gene enrichment in oxidative phosphorylation, metabolic pathways and the tricarboxylic acid (TCA) cycle (Fig. 8b and Supplementary Data 13).

**Fig 8:**
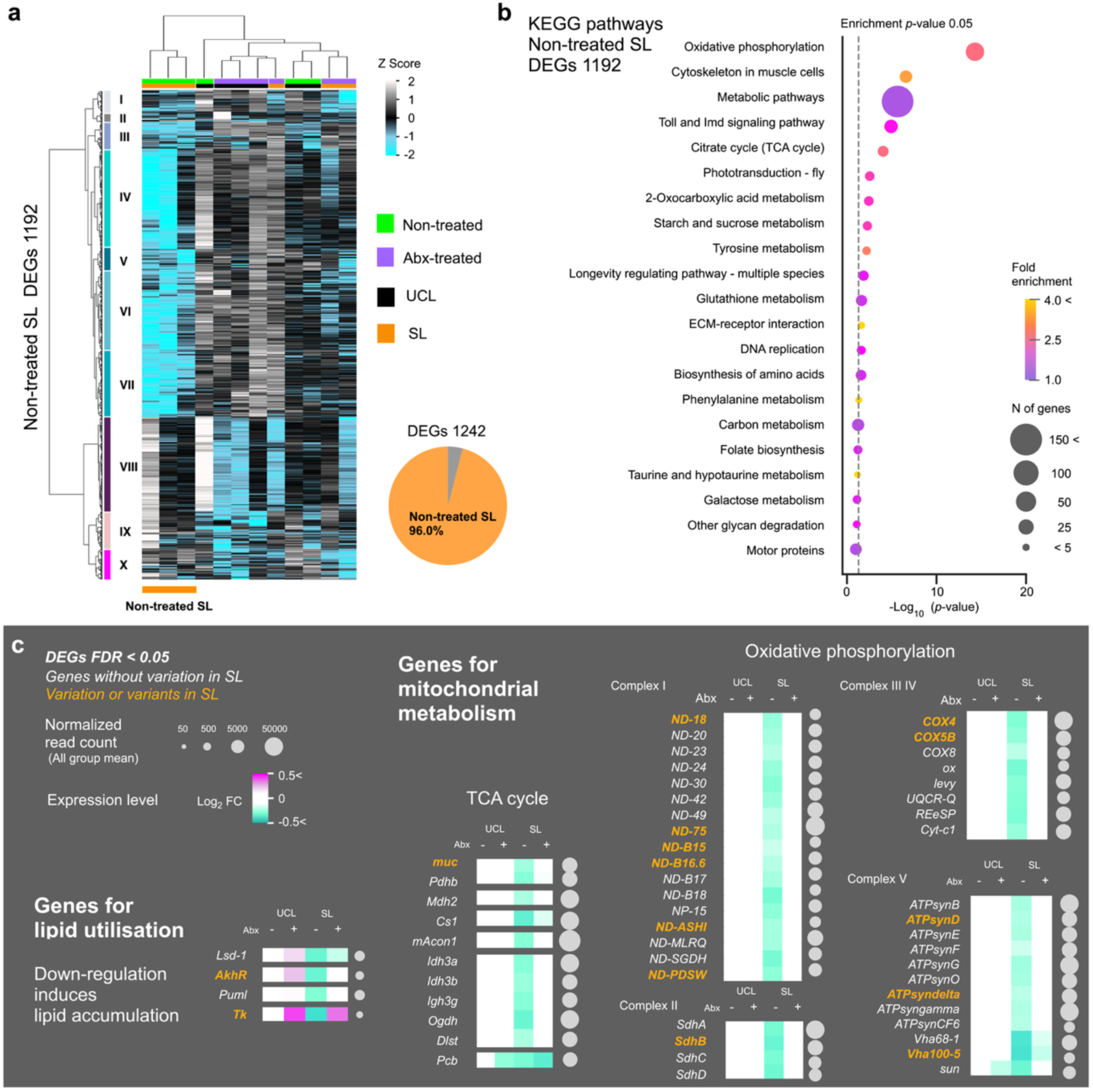
The characteristic phenotype of SL flies is explained by the host transcriptome. **a**, Transcriptome analysis of the whole body of virgin females under non-irradiated conditions. Cluster map of non-treated SL-specific differentially expressed genes (DEGs) with a false discovery rate (FDR)-adjusted *p*-value threshold of <0.05. The total number of DEGs across all comparisons was 1,242, of which 1,192 (96%) were specific to non-treated SL (pie chart); replicate line 1, generation 50. **b**, Enrichment analysis of non-treated SL DEGs using KEGG. **c**, Heatmaps illustrating functional profiles. All genes shown are DEGs (FDR-adjusted *p*-value threshold of < 0.05). Expression levels are presented as mean log2 FC for each group, calculated by first calculating log2 FC for individual samples relative to the average expression in non-treated UCL and then averaging across groups. The list of genes includes those known to induce lipid accumulation through genetic manipulation and related mitochondrial metabolism. Genes with orange symbols represent either SL-specific variations (above 5%) or common homozygous variants in SL. Circles indicate mean read counts across all groups. For details, see Supplementary Data 13.

Suppressed mitochondrial metabolism is known to lead to lipid accumulation in various animal species^60,61^. Consistent with this fact, non-treated SL flies showed downregulation of most of the genes involved in oxidative phosphorylation and the TCA cycle, which are associated with mitochondrial metabolism (Fig. 8c). In addition, several genes whose downregulation is known to induce lipid accumulation were also specifically downregulated in non-treated SL flies (Fig. 8c)^21,22,62^, including *Tachykinin* (*Tk*), which also harbored mutation in the SL flies (Fig. 8c and Supplementary Data 13). Actually, *Tk* expression is known to be modulated in response to acetate produced by gut microbiota, thereby regulating lipid utilisation in enterocytes^37,63^. As expected, approximately 30% of DEGs (387 genes) reflected the distinctive genomic profile of SL, with the Toll and Imd signaling pathways appearing most enriched (Supplementary Fig. 18a–d). Of these, 276 genes contained homozygous variants, including 6 genes associated with the Toll and Imd signaling pathways, such as *Toll* (Supplementary Fig. 18b and Supplementary Data 14). Our results suggest that the phenotypes of SL flies are influenced by the Hippo signaling pathway, *Tk*, and mitochondrial genes, and by their interactions with gut microbiota.

### Lipid accumulation via genetic manipulation confers BL tolerance

Our finding suggest that SL has acquired a gut microbiota-driven lipid accumulation (Fig.4, Fig. 5 and Fig. 8c). To prove the causal relationship between lipid accumulation and BL tolerance, we examined BL tolerance in genetically manipulated flies that are reported to have increased TAG levels^21,22^. We used a range of mutants and GAL4/UAS-mediated manipulations, including knockdowns of *Tk*, *AkhR*, and *Lipid storage droplet-1 (Lsd-1*), which were downregulated in the SL flies (Fig. 8c, Supplementary Data 13 and Supplementary Data 15). Interestingly, knocking down these genes enhanced BL tolerance to a level comparable to that in the SL flies (Fig. 9a). Furthermore, BL tolerance was also enhanced by other genetic manipulations that are known to increase lipid levels but were not detected as DEGs in our transcriptome analysis (Fig. 8c), such as knockdown of *brummer* (*bmm*), *Phosphatase and tensin homolog* (*Pten*), and *Stromal interaction molecule* (*Stim*), and overexpression of *Insulin-like receptor* (*InR*), *Lk6 kinase* (*Lk6*), and *Lipid storage droplet-2* (*Lsd-2*) (Fig. 9a). Indeed, we found a significant positive relationship between fecundity and TAG levels among flies with all of these genetic modifications (Fig. 9b), which strongly suggests that lipid accumulation is a trait that confers BL tolerance (Fig. 9c).

**Fig 9:**
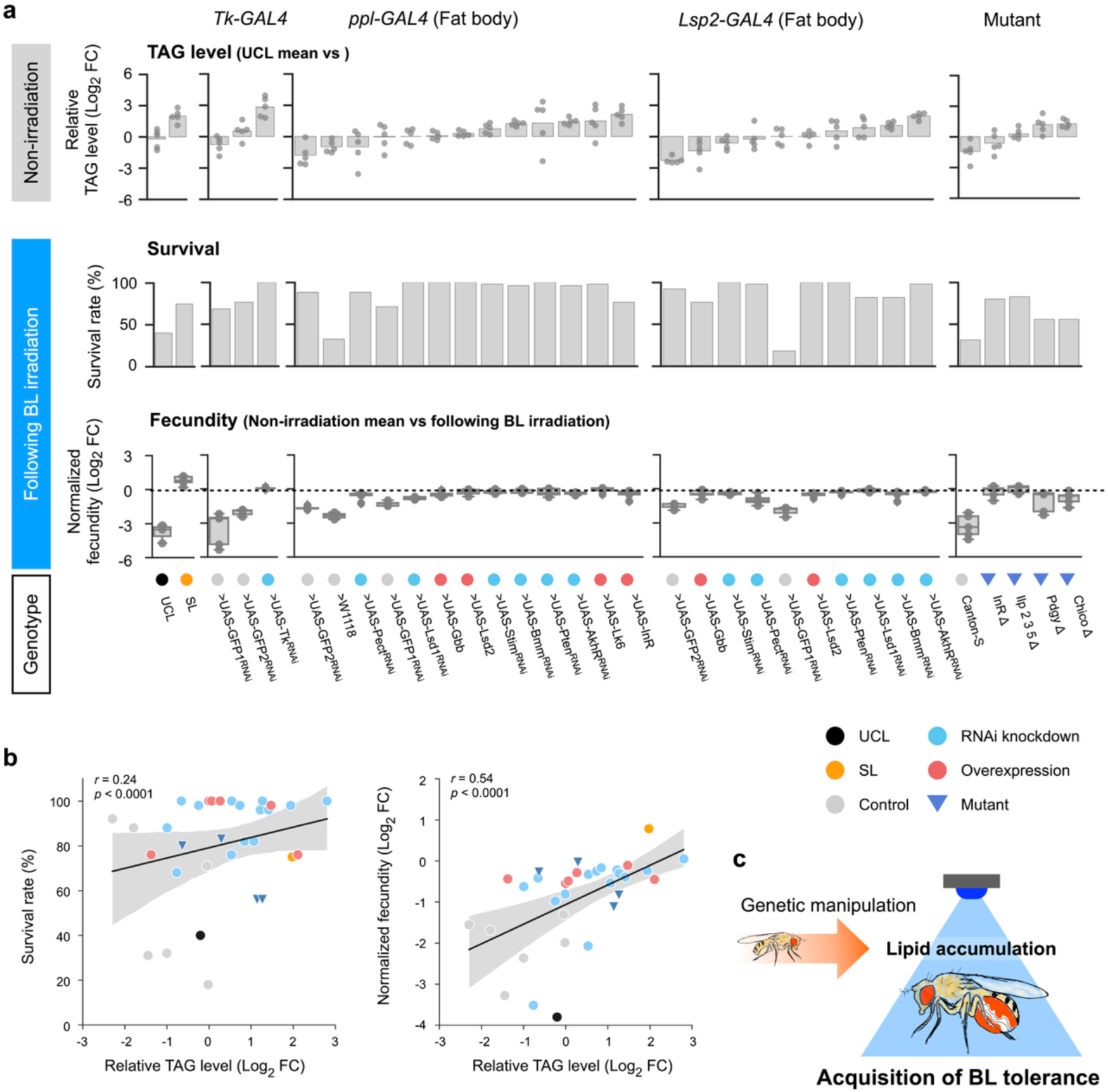
Lipid accumulation via genetic manipulation confers BL tolerance. **a**, Relative TAG levels, survival rates following BL irradiation, and normalised fecundity. TAG levels are presented as Log2 FC values relative to mean UCL levels. Data are arranged in the order of UCL, SL, the *Tachykinin* (neuropeptide) targeting *Tk-GAL4* driver, the fat body targeting *ppl-GAL4* and *Lsp2-GAL4* drivers and mutant lines. The order is sorted by TAG levels, with higher values on the right. Survival rates after 3 days of BL exposure were measured on a minimum of 40 and a maximum of 120 flies per genotype (Supplementary Data 15). Genotypes with less than 10% survival were excluded from the data. The full list is available in Data S4. Fecundity was normalised by calculating the log2 FC in the number of offspring after BL exposure relative to the unexposed control for each genotype. Negative values indicate a reduction in fecundity due to BL toxicity, while zero indicates no effect. Positive values observed in SL only indicate a beneficial effect. Raw fecundity data, including the number of offspring in both non-irradiated and BL irradiated conditions, are presented in Supplementary Data 17 (Fig. 9a). Genotype labels and colors indicate the effector, the target gene and the type of manipulation (overexpression; red, knockdown; skyblue or null mutation; royalblue). UCL and SL TAG levels were derived from the same dataset as in Fig. 4f (replicate line 1, generation 57). Box plots show the median (centre line), the interquartile range (box), the full range of the data (from minimum to maximum). **b**, Correlation analysis. *n* = 33, Data using mean values for each genotype in Fig.9a. The variance represents the 95% confidence interval. **c**, Summary of experimental results: Lipid accumulation induced by genetic manipulation confers BL tolerance. For details, see Supplementary Data 15.

### The transcriptome of *A. persici* in the host and the growth patterns vary between the host strains

Finally, we tried to investigate the possible mechanisms underlying the contribution of *Acetobacter*. To our knowledge, seventeen *Acetobacter* species have been identified as gut bacteria in *Drosophila* and are known to affect host metabolism and fitness^31,33,35,38,41–46,64,65^. Phylogenetic analysis based on reported full-length 16S rRNA gene sequences suggesting that *A. persici* is closely related to *A. malorum* and *A. cerevisiae* (Fig. 10a). In both UCL and SL, the dominant OTU ID (397 bp) representing 89-98% of the microbiota was identified as *A. persici* Dm-48 strain with 99.75% sequence identity (Fig. 10a, Supplementary Fig. 19). Analysis of the complete genome of *A. persici* using publicly available data showed that its chromosome is 3,230,507 bp, while its plasmid is 526,169 bp, containing 2,898 and 492 CDS, respectively (Fig. 10b). In order to infer the transcriptomic profile of *A. persici* within the host, we first extracted reads that were not mapped to the host genome in the RNA-seq data, and then mapped these reads to the complete *A. persici* genome. Bacterial reads were detected in non-treated flies, whereas fewer than 1,000 reads were recovered from Abx-treated flies, likely reflecting reduced bacterial abundance. As this mRNA was obtained by poly(A) selection of the host, there is a possibility of bias in the results. The sequencing depth was approximately 10X, primarily capturing extreme variation and probably missing low-expressed genes (Fig. 10c, Supplementary Fig. 20a and Supplementary Data 16). Nevertheless, a total of 225 DEGs were detected, including 124 downregulated and 101 upregulated genes (Supplementary Fig. 20b and Fig. 10d). Gene Ontology (GO biological process) enrichment analysis revealed a frequent enrichment of metabolic pathways, suggesting that the colonisation state of the *A.persici* within the SL differs from that in the UCL (Supplementary Fig. 20c and Supplementary Data 16). *A. persici* harbours 15 genes involved in acetic acid production^66^. Interestingly, key genes such as *Alcohol dehydrogenase* (*ADH*) and *Acetyl-CoA hydrolase* (EC 3.1.2.1) were found to be specifically upregulated in the SL (Fig. 10d and Supplementary Data 16). These upregulated genes may contribute to the microbiota-dependent lipid accumulation of SL. The top 5 genes with the lowest *p*-values were all upregulated and included potentially oxidative stress-related genes such as *NADPH-dependent FMN reductase* and *Paraquat-inducible protein* (Fig.10d). The ability of *A. tropicalis*, a close relative of *A. persici*, to restore host survival via microbiota-mediated detoxification was previously documented (Fig. 10a)^64^. Given that antioxidant mechanisms are thought to contribute to resilience to BL toxicity, it is noteworthy that genes associated with the oxidation-reduction process were detected as enriched DEGs (Fig. 10f and Supplementary Data 16). The upregulation of oxidoreductases and translocases indicates their potential antioxidant role by neutralising ROS and transporting key metabolites that support cellular redox balance (Fig. 10f). These results suggest that *A. persici* in SL has a different transcriptomic profile, especially in metabolism. However, it remains uncertain whether this difference is due to bacterial genomic variation, differences in bacterial colonisation state or host-mediated effects. We then investigated whether *A. persici* showed phenotypic differences by analysing the growth patterns of isolates from each host strain. Fifty colonies were cultured from each host strain. Consistent with the transcriptomic variation, *A. persici* isolates from the UCL and SL strains showed divergent growth patterns (Fig. 10g^i^– g^iv^ and Supplementary Data 16). In the SL, 26% (13 colonies) failed to grow, whereas only 8% (4 colonies) failed to grow in the UCL. Additionally, UCL showed a higher variance, whereas SL showed a lower variance (Fig. 10g^iii^). This phenotypic difference indicated a potential bias towards SL-dominant bacterial strains. Overall, these results suggest that BL toxicity may have acted as a selective pressure shaping bacterial strain dynamics or driving host-microbiota co-evolution.

**Fig 10:**
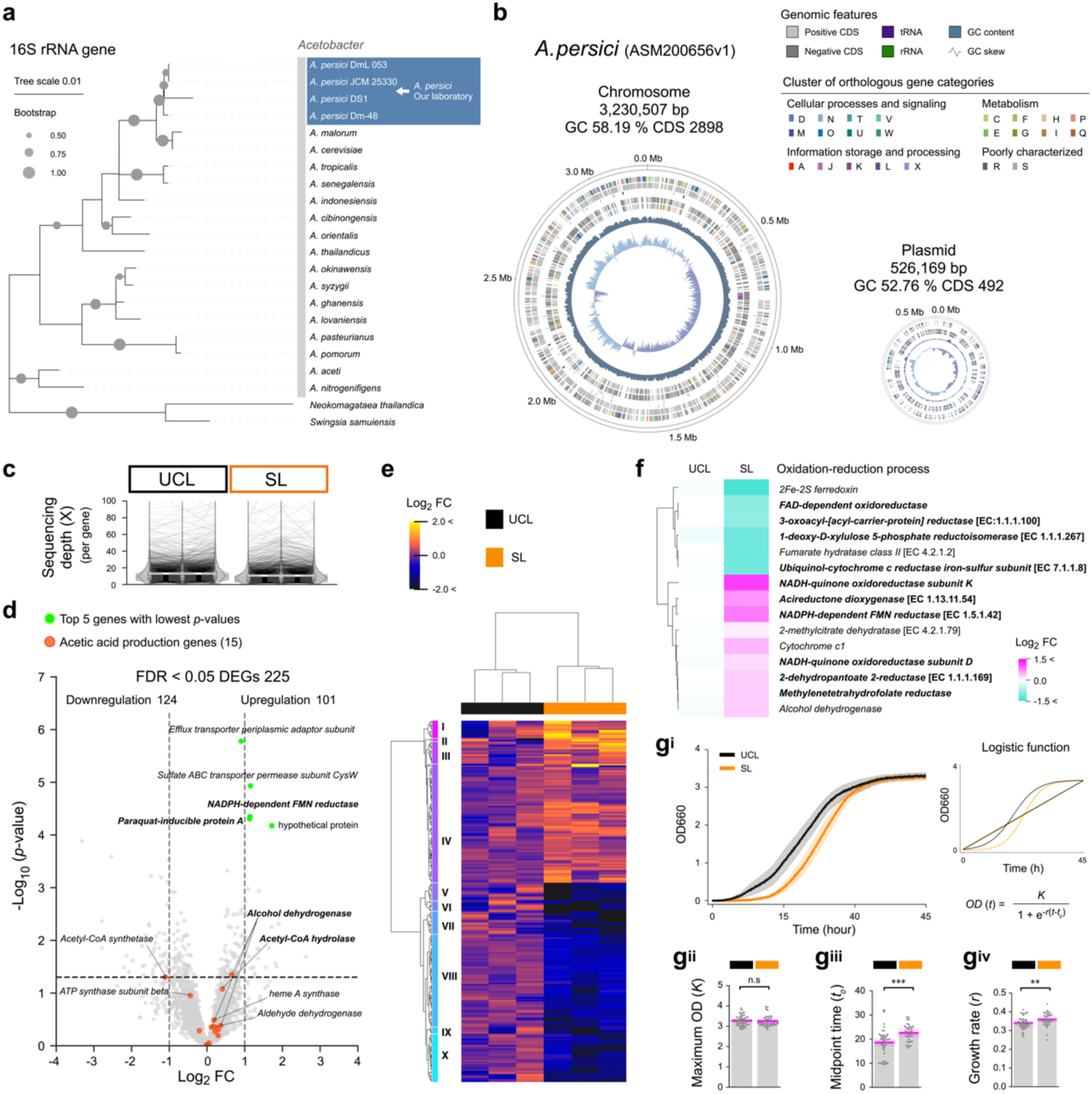
Phylogenetic relationships, distinct transcriptome and divergent growth patterns of *A. persici* in SL. **a**, Phylogenetic tree based on the 16S rRNA gene of *Acetobacter* species identified in the *Drosophila* gut microbiota. *Neokomagataea thailandica* and *Swingsia samuiensi*s were used as outgroups. Several strains of *A. persici* were analysed to improve resolution. The arrows indicate the inferred phylogenetic position of the dominant *A. persici* in our laboratory. As the full-length sequence was not available, it was excluded from the phylogenetic tree. See Supplementary Fig. 19 for the phylogenetic relationships of the *A. persici* strains. The tree was constructed using maximum likelihood with 1,000 bootstrap replicates. **b**, Complete genome of *A. persici* based on ASM200656v1. The alphabetical annotations represent functional categories based on the Database of Cluster of Orthologous Genes. **c**, Sequencing depth calculated from unmapped *Drosophila* reads mapped to *A. persici.* Sequencing depth estimated from the length per gene length and total bases mapped to the gene (mean values for UCL: 13.10, SL: 10.63). Genes corresponding to the same locus are connected by lines. The central white line represents the mean value. See Supplementary Fig. 19a for an overview of the depth distribution. See Fig. 8 for host transcriptome data. **d**, Number of *A. persici* DEGs across host strains and corresponding volcano plot. Genes with the lowest *p*-values in the top five are shown in lime, while those involved in acetic acid production are highlighted in red. Genes highlighted in the main text are shown in bold. **e**, Cluster map of *A. persici* DEGs (FDR-adjusted *p*-value <0.05) in the SL. **f**, List of genes associated with the oxidation-reduction process (Gene Ontology: biological process). Only DEGs (FDR-adjusted *p*-value <0.05) are shown. Bold indicates oxidoreductases or translocases. **g^i^**, Bacterial growth curve between isolates from host strains. *n* = 44 (UCL), 37 (SL). The bacterial growth curve shows the mean OD values with a 95% confidence interval, estimated by bootstrap resampling (*n* = 10,000, seed = 4). **g^ii^**, Maximum OD (*K*); BF10 = 0.52, **g^iii^**, Midpoint time (*t0*); BF10 = 0.37, **g^iv^**, growth rate (*r*); BF10 = 8.94, Asterisks indicate significance: ** *p*MCMC < 0.01; *** *p*MCMC < 0.001, n.s = not significant; replicate line 1, generation 52. **g^i^**–**g^iv^** represent the same dataset. Statistical analyses for panels **g^ii^**–**g^iv^** were performed using HBM with MCMC methods, posterior median, and Bayesian 95% CI. The magenta, bold lines and shaded areas show the posterior median and the Bayesian 95% CI. For details, see Supplementary Data 16 for all results, including statistical models.

## Discussion

In this study, we established BL-tolerant *Drosophila* strains through laboratory selection, demonstrating that excessive BL exposure drives the ‘adaptive obesity’, particularly lipid accumulation mediated by gut microbiota. This supports the increasingly recognized role of microbiota in host adaptation to environmental stress, in which specific bacterial populations contribute to adaptive host phenotypes^33,43,67–69^. Our findings further reveal that elongation of the host midgut in response to parental BL toxicity facilitates an increase in beneficial bacteria, ultimately enhancing host fitness via lipid accumulation under excessive BL exposure (Supplementary Fig. 21).

### Lipid storage is one of the factors that determines BL tolerance

SL flies exhibited increased lipid accumulation (Fig. 4), and genetic manipulation that increased lipid contents conferred BL tolerance (Fig. 9). This acquisition of BL tolerance can be explained by the role of lipid accumulation in protecting against ROS, which are produced upon BL irradiation^12,13^. Intracellular lipid droplets play a critical role in shielding cells from oxidative stress, as reduced TAG levels are associated with increased sensitivity to such stress^23–25,70^. Indeed, the accumulation of lipid droplets in response to intracellular ROS has been observed in many species^25,70^. Additionally, lipid droplets are closely linked to fecundity, as higher TAG levels are associated with improved fecundity, whereas reduced TAG levels can lead to oocyte degeneration^71,72^. Thus, lipid accumulation not only helps defend against BL toxicity but also supports the maintenance of fecundity. Furthermore, the transcriptome of SL flies revealed a broad downregulation of genes for mitochondrial metabolism (Fig. 8c). Mitochondria, while crucial for energy production, also contribute to ROS generation^73^. It has been reported that BL toxicity causes a decrease in ATP production^12,13^. Therefore, inhibition of both mitochondrial metabolism and lipid utilisation may be an adaptive strategy to accumulate lipids while minimising oxidative stress, and obesity may serve to compensate for insufficient ATP levels.

### The potential of the gut microbiota for benefit enhancement in *Drosophila*

*Acetobacter* is generally the dominant bacteria in *Drosophila*^34–36,41,42^. The bacterial composition and relative abundance within the microbiota varied between laboratories, with *Acetobacter* sometimes accounting for over 90% of the community^30,32–34^. Generally, the microbiota of laboratory flies is influenced by factors such as the host genome and environmental conditions, including diet^34,36,42^. We identified *A. persici* as the predominant bacterial species and showed the critical role of its increased abundance in the acquisition of BL tolerance. *Acetobacter* species, which are well-known symbionts of *Drosophila* and other insects, are recognized for their acetate production^74^. Acetyl-CoA, derived from acetate, is essential for lipid utilisation^28,75^ and also plays a crucial role in the acetylation of both histone and non-histone proteins, facilitating adaptive responses through tissue plasticity^37,76,77^. The elevated TAG levels observed in SL flies may be explained by the increased acetyl-CoA pool resulting from gut microbiota–derived acetate. The increased expression of acetic acid production genes in *A. persici* from the SL supports this hypothesis (Fig. 10d)^66^. Additionally, gut microbiota might act as a barrier against physiological stress induced by BL toxicity. The significant reduction in gut microbiota following BL irradiation in SL flies suggests the evolution of mechanisms that utilise bacteria as a direct nutritional resource or as enzymes (Supplementary Fig. 8 and Fig. 10f). These findings align with previous research showing that *Acetobacter* enhances the fitness of the host fly^29,35,43–45^. Notably, *Acetobacter* infection enhances oxidative stress tolerance in *Drosophila*, whereas oxidative stress reduces *Acetobacter* abundance, consistent with the findings of Brown *et al.* (Supplementary Fig. 8 and Supplementary Fig. 11)^64^. Several studies in *Pseudomonadota*, including *Acetobacter*, have shown that gut bacterial enzymes in nematodes and insects adaptively contribute to host stress tolerance^33,64,78,79^. Consistently, our transcriptome analysis revealed that antioxidant enzymes are part of the upregulated cluster in *A. persici* from SL (Fig. 10d, f and Supplementary Data 16), suggesting their potential role as bacterial metabolism benefiting the host. In addition, the infection experiments primarily suggested that bacterial abundance strongly influenced the outcome; yet infection with gut contents did not fully rescue the fecundity of SL (Fig. 5b and Supplementary Fig. 13b). This implies that in addition to bacterial abundance, qualitative differences in *A. persici* (such as colonisation status, within the host response, metabolism and bacterial strains) may also play a critical role in host BL tolerance. Given that bacterial evolution is likely to be faster than host genetic change, it is plausible that some genetic differences have arisen in SL-derived *A. persici*. This is suggested by the slightly higher weight gain rate observed for SL-derived *A. persici* in infection experiments with standardised bacterial concentrations, and by differences in the growth patterns of isolated bacteria, with SL strains showing less variance (Supplementary Fig. 12c and Fig.10g^i^, g^iii^). However, the rate of microbiome evolution during the experimental evolution process is unknown. The difference in bacterial phenotypes highlights the importance of investigating bacterial strain dynamics in response to BL toxicity and host-microbiota co-evolution in future studies.

### Acquisition of BL tolerance in SL via adaptive obesity

When specific bacteria are implicated in host fitness-related phenotypes, their relative enrichment often leads to reduced microbiota diversity^33,68^. Given that the microbiota composition did not differ between strains (Supplementary Fig. 7a), the initial diversity of the gut microbiota in our study was probably low. Despite low microbiota diversity, increased abundance of *A. persici*, possibly together with reduced strain diversity (dominance of specific strains), appears to confer an adaptive phenotype in SL. Characteristic features such as shortened wings and an elongated midgut were also observed in the SL phenotype. We propose that gut elongation plays a key role in evolutionary adaptation driven by the gut microbiota. Both scenarios could contribute to the formation of the SL phenotype: changes in organ size may be influenced by independent pathways or result from a trade-off in resource allocation. The increase in bacterial abundance appears to be associated with midgut elongation and genetic changes, particularly those in the Hippo signaling pathway and the Toll and Imd signaling pathways (Fig. 4c, Fig. 5f, Fig. 6c, Fig. 7d, e and Supplementary Fig. 18b). Because our bacterial quantification was based on CFU measurements from whole-body homogenates, the direct relationship between midgut elongation and bacterial abundance remains uncertain. Nevertheless, the observation that SL flies retain an elongated midgut after Abx treatment, but experience a significant reduction in body weight, suggests that gut microbiota are necessary for the acquisition of obesity (Supplementary Fig. 10). The resulting phenotype from laboratory selection may have been influenced by a combination of epigenetics, de novo mutations, genetic drift, and variant selection. We are not able to separate these contributions, and further analysis will be needed to clarify the underlying mechanisms. The *Drosophila* total sequence length is approximately 140 Mbp^80^, with a de novo single nucleotide mutation (SNM) rate of 3.3 × 10^-9^ per site per generation^81^. This corresponds to an estimated 0.462 de novo SNMs per generation across the genome. Over 44 generations (as in the WGS performed), this amounts to approximately 26 de novo SNMs. However, we identified SL-specific homozygous variants at 1,985 sites (corresponding to 658 genes) shared by the three replicate lines, a level that cannot be explained by the accumulation of naturally occurring variants alone. Although de novo mutations may have occurred, we predict that most of the SL-specific variants, including both SNPs and indels, are derived from standing variations already present in the ancestral population. The possibility that BL toxicity increases the SNM rate cannot be ruled out and requires further investigation of the molecular evolutionary process. We also found that parental irradiation with BL, when combined with accumulated mutations in later generations of SL flies, is critical for the midgut elongation and bacterial abundance (Fig. 6b, c). This is consistent with previous studies showing that parental exposure to environmental stressors, such as starvation and BL toxicity, induces epigenetic inheritance or alternations in the transcriptome^48,82^. In *Drosophila*, the midgut undergoes plastic elongation or shortening in response to environmental conditions^83,84^. This physical expansion of the gut, which increases spatial capacity, offers several advantages, including more efficient nutrient absorption, increased bacterial abundance, and maintenance of gut homeostasis^71,85–89^. However, midgut elongation alone, even after one generation with parental exposure to BL toxicity, is insufficient for BL tolerance (Fig. 2a and Fig. 6c). Full BL tolerance requires both midgut enlargement and specific genetic changes. Among these genetic factors, the genes for the neuropeptide *Tk* and its receptor *TkR99D* are particularly important (Fig. 8c, Fig. 9 and Supplementary Data 11), as they may regulate lipid utilisation in response to acetate produced by the gut microbiota^37,62^. Laboratory evolution studies typically use genetically diverse populations, such as wild populations or multiple lines from the *Drosophila* Genetics Reference Panel^90^. As we used Canton-S (low genetic diversity), it is possible that microbiota-mediated lipid accumulation was more optimal than the development of dedicated lipid synthesis systems driven by host genetic changes. Microbiota-mediated evolutionary adaptation and co-evolution under low host genetic diversity remain interesting topics for future research.

### Implications of our findings for pest control

BL toxicity indicates wavelength-specific effects across insect species and developmental stages^9^. Reports also show no changes in sensitivity between mutant or overexpression models of photoreceptors^10^, suggesting a complex underlying mechanism. Our study identifies lipid storage and gut microbiota as critical factors in BL tolerance. In addition, we have shown that BL toxicity strongly inhibits ovarian development and exerts persistent suppressive effects even in the absence of lethality (Supplementary Fig. 2). In the context of pest control, the combination of BL irradiation with treatments that disrupt insect lipid storage and gut microbiota could potentially improve efficacy. In parallel with the development of these methods, research into whether similar tolerance mechanisms exist in other insect species could lead to more effective chemical-free BL pest control.

Our findings highlight the importance of gut microbiota–mediated lipid accumulation as an adaptive trait for BL tolerance. The elongation of the midgut, associated with an increased abundance of beneficial bacteria, turns out to be a key factor in this evolutionary adaptation. Such phenotypes may be driven by genetic and epigenetic changes from selective pressures and parental BL exposure. These evolutionary responses to excessive BL exposure demonstrate how host traits evolve under a toxic BL environment by maximising the benefits provided by gut microbiota. Our study reveals an insect evolutionary process via the microbiome and presents a concept of evolutionary adaptation initiated by midgut elongation.

## Methods

### Fly cultures and diet

Flies were maintained at 23–26°C on a 16:8 hour light:dark cycle using cold cathode fluorescent lamps as the light source. Detailed light conditions are provided in Supplementary Fig. 1. The fly diet consisted of 5 g glucose (Fujifilm Wako Pure Chemicals Co., Osaka, Japan), 6 g dry brewer’s yeast (Asahi Group Holdings Ltd., Tokyo, Japan), 1 g agar (Fujifilm Wako Pure Chemicals Co.), 0.4 mL propionic acid (Fujifilm Wako Pure Chemicals Co.), 0.4 mL 70% ethanol (Fujifilm Wako Pure Chemicals Co.), 0.01 g butyl *p*-hydroxybenzoate (Fujifilm Wako Pure Chemicals Co.), and 100 mL distilled water.

### BL irradiation

Virgin flies aged 1–5 days were used. Unless otherwise specified, flies were exposed to 462 nm BL for 3 days. BL irradiation conditions varied across experiments:

Fig. 2b–c, Fig. 5b, Fig. 6b, c, Supplementary Fig. 2a, b, Supplementary Fig. 3a, Supplementary Fig. 8, Supplementary Fig. 9c, d, Supplementary Fig. 12b, c and Supplementary Fig. 13b: Photon flux density 10.3 × 10¹⁸ photons·m⁻²·s⁻¹ for females. Fig. 2a, c, Supplementary Fig. 3a: 6.5 × 10¹⁸ photons·m⁻²·s⁻¹ for males.

Fig. 2d: Continuous BL irradiation for 5 days, 8 × 10¹⁸ photons·m⁻²·s⁻¹ for males and 10 × 10¹⁸ photons·m⁻²·s⁻¹ for females.

Supplementary Fig. 4: Continuous BL irradiation for 10 and 4 days, 5 × 10¹⁸ photons·m⁻²·s⁻¹ and 15 × 10¹⁸ photons·m⁻²·s⁻¹ for females.

Fig. 9: 465 nm BL, 9.5 × 10¹⁸ photons·m⁻²·s⁻¹ for females.

Irradiation was performed in a multi-room incubator (LH-30CCFL-8CT; Nippon Medical & Chemical Instruments Co., Ltd, Osaka, Japan) at 24–26°C, with LED panels (IS-mini^®^, ISL- 150 × 150 Series; CCS Inc., Kyoto, Japan) installed on the ceiling. Photon flux density was measured using a high-resolution spectrometer HSU-100S (Asahi Spectra Co., Ltd., Tokyo, Japan) with ND 0050 filter (Asahi Spectra Co., Ltd) and spectroradiometer MS-730 (EKO Instruments Co., Ltd., Tokyo, Japan). For details see Supplementary Fig. 1. All light condition measurements used in the experiment are included in the Supplementary Data 17 (Light condition).

### Laboratory selection for BL tolerance

We used the Canton-S strain of *Drosophila* as the ancestral population for laboratory selection. The Canton S strain used in this study was obtained from Tokyo Metropolitan University in 2016. It has been maintained under laboratory conditions for at least 30 years and has been bred in our laboratory in over 1,000 populations over 6 years. Two strains were established, SL and UCL; the former was subjected to BL toxicity every generation and the latter was used for controls. Each strain included three replicate lines. For each strain, 60–100 virgin flies aged 1–5 days were placed in two 60 mm × 90 mm glass Petri dishes for BL irradiation. SL and UCL flies (positive control group) were exposed to 462 nm BL for 3 days. Instead of BL irradiation, the UCL, used as a negative control group, was kept in darkness for 3 days. We followed this standard approach as dark conditions are commonly used as a control in BL toxicity studies^7,10,11,48^.

For climbing assays, Irradiated flies were placed in a breeding environment (for details, see “Fly cultures and diet” in Methods) for a 2-day recovery period. For the data shown in Supplementary Fig. 3a and Supplementary Fig. 9c, the impact of BL toxicity was assessed in a 4-minute climbing assay and the flies were categorized into four states: “Low impact”: *Intact*, capable of climbing.

“High impact” (unable to climb): *Phase 1*, alive but unable to climb; *Phase 2*, alive but non-ambulatory; *Phase 3*, dead.

For the artificial selective breeding process, the "Low impact" SL flies were allowed 5 days for mating and egg-laying. In the early stages of selection, there was no clear difference in climbing ability, and fecundity was low; thus, we used as many intact individuals as possible as parents. From the 17th generation, when tolerance began to emerge, we prioritised intact individuals that climbed above the 2 cm threshold and selected preferentially for higher tolerance. Each vial containing 3 mL of diet medium 3-mL contained 9–13 pairs, with a total of 4–8 vials (for details, see Supplementary Data 17 (Fig. 2a)). After 5 days, the diet medium containing eggs and larvae was transferred to a larger plastic box (40 mL diet medium, 72 × 72 × 100 mm; Insect Breeding Square Dish, SPL Life Sciences, Gyeonggi-do, Korea) to ensure sufficient food supply. One box was used for every 4 or 5 vials. UCL parents were randomly chosen from the negative control group and reared under the same conditions as the SL flies. To evaluate fecundity, the total number of pupae was counted and divided by the number of parents to determine the fecundity per female for each generation.

### Negative geotaxis climbing assay in non-irradiated conditions

Young adult flies, which typically climb 4 cm in 4 seconds, were tested for negative geotaxis^91^. In Supplementary Fig. 3b, ten flies aged 5–10 days were placed in an empty vial, gently tapped three times to ensure they reached the bottom and given 10 seconds to climb. Climbing rate was defined as the flies reaching a 6 cm reference line within this time. Each vial was tested three times with a 1-minute interval between trials and the mean climbing rate was calculated.

### Survival assays

Virgin flies aged 5–10 days were used. In the continuous BL irradiation assay, 50 or 42 flies were placed in each glass Petri dish containing standard diet medium and exposed to 462 nm BL (Fig. 2d and Supplementary Fig. 4). In the starvation assay (Fig. 4g), 10 flies were placed in each vial with 1% (w/v) agar (Fujifilm Wako Pure Chemicals Co.) in distilled water. In the oxidative stress assay, 10 flies were placed in each vial with a medium prepared with either 10 mM paraquat dichloride standard (Fujifilm Wako Pure Chemicals Co.) (Fig. 4h) or 10 mM 1,1’-dimethyl-4,4’-bipyridinium dichloride (Tokyo Chemical Industry Co., Ltd., Tokyo, Japan) (Supplementary Fig. 11a, b), 5% (w/v) glucose (Fujifilm Wako Pure Chemicals Co.), and 1% (w/v) agar (Fujifilm Wako Pure Chemicals Co.) in distilled water. The surviving flies were manually counted every 2 h. Survival curves were visualised using the KaplanMeierfitter from the lifelines (Python version 3.8.3 version 0.25.9).

### Measurement of fecundity

To measure fecundity (Fig. 2a, Fig. 5b, Fig. 9, and Supplementary Fig. 13b), after 3 days of darkness (non-irradiation) or BL irradiation followed by a 3-day recovery period, 5 surviving females were randomly chosen and mated with 5 non-irradiated virgin males (aged 1–10 days) in individual pairs within vials containing 5 mL of diet medium for 4 days of egg-laying. The total numbers of pupae and emerged adults were counted to evaluate fecundity and emergence rate (in Fig. 2a and Fig. 2c, only the total number of pupae was measured).

Fig. 2c used non-irradiated males from either the UCL or SL. Fig. 5b and Supplementary Fig. 13b used non-irradiated, Abx-treated males from the UCL or SL. Fig. 9 used non-irradiated Canton-S males.

### Measurement of body weight, organ size and oocyte number

Phenotypic measurements were performed on non-irradiated flies in Fig. 3a, b, Fig. 4c, Fig. 5a, d, f, Fig. 6c, Supplementary Fig. 8 and Supplementary Fig. 14 and on BL-irradiated flies in Fig. 2b, Supplementary Fig. 2a, b and Supplementary Fig. 8. As ovarian development can progress with age, we ensured that phenotypic comparisons between strains were made at the same chronological age^19^. Virgin flies aged 4–12 days were anesthetized by chilling on ice. Body weight was measured using an AUW220D balance (Shimadzu, Kyoto, Japan). Wings or ovaries were dissected and photographed under a SteREO Discovery.V12 stereomicroscope and an Axiocam 305 color camera (Carl Zeiss, Oberkochen, Germany). Wing length was measured as the distance between the alula opening and the distal edge of the 3rd longitudinal vein, using Fiji ImageJ2 (version 2.3/1.54f), following a previous method^92^. Stage 14 oocytes were counted under the stereomicroscope. Midgut lengths of regions R1 to R5 were measured manually from the midgut images using Fiji ImageJ2^89^.

### Lipid droplet staining and imaging

In Fig. 4b^i^ and b^ii^, we performed dissection and fluorescence microscopy to visualise lipid droplets in the fat body of the abdominal epithelium of virgin female flies aged 6–12 days kept in non-irradiated conditions. After removing other tissues from the abdomen, we fixed the abdominal epithelium in PBS containing 4% paraformaldehyde (Fujifilm Wako Pure Chemicals Co.) for 20 minutes. After fixation, we washed the samples twice with PBS (Gibco, Carlsbad, CA, USA) for 15 minutes each time. The fixed abdominal epithelium was stained with Nile Red (Fujifilm Wako Pure Chemicals Co.) at a concentration of 10 μg/mL overnight according to a previous method^93^. Following this, we washed the samples twice with PBS for 10 minutes each time and then stained them with NucBlue (Thermo Fisher Scientific, Waltham, MA, USA) for 3 minutes, followed by a 5-minute wash with PBS. All staining procedures were conducted in a light-shielded environment at room temperature. Imaging was performed under an Eclipse Ti2 microscope equipped with a DS-Ri2 camera (Nikon, Tokyo, Japan). The images were analysed using the Python package OpenCV (Python version 3.8.3 version 4.5.3.56) to obtain the mean fluorescent intensity of Nile Red.

### Lipid extraction and derivatization of fatty acids

Lipids were extracted using the Bligh and Dyer method^94,95^. Two virgin female flies aged 6– 12 days and kept in non-irradiated conditions were homogenised in 100 μL of PBS (Gibco), added 600 μL of methanol (Fujifilm Wako Pure Chemicals Co.) and chloroform (Fujifilm Wako Pure Chemicals Co.) in a 2:1 (v/v) ratio, vortexed for 1 minute, and shaken at 1000 rpm for 2 h at 4°C. Next, we added 200 μL of chloroform and 250 μL of distilled water, vortexed for 1 min, centrifuged at 9000 rpm for 2 min, and collected the organic lower phase. We then added 400 μL of chloroform to the remaining mixture, vortexed for 1 minute, and centrifuged as above. The two organic extracts were pooled and dried under vacuum. The dried lipids were derivatized using the Fatty Acid Methylation Kit (Nacalai Tesque, Inc., Tokyo, Japan) and the Fatty Acid Methyl Ester Purification Kit (Nacalai Tesque, Inc.). FAMEs were collected.

### Gas chromatography mass spectrometry (GC-MS) analysis

The FAMEs were analysed using a GC-MS instrument (GCMS-QP2010 Ultra (Shimadzu)) equipped with a DB-5 ms column (30 m × 0.25 mm ID, 0.25 µm film thickness) (J&W Scientific, Folsom, CA, USA). Helium was used as the carrier gas at a pressure of 100 kPa. The column oven was set at 40°C for 5 minutes. The temperature was increased at a rate of 10°C/min to 280°C and maintained at 280°C for 10 minutes. The injection port temperature was 220°C, and the analysis was performed in split mode. Fatty acids were identified using a similarity search against the standards Supelco F.A.M.E. Mix C4-C24 (Supelco, Bellafonte, PA, USA) and relevant references^96,97^. The total ion chromatogram area for each compound was measured. On the basis of the obtained data, we compared the total amount of FAMEs. The log-transformed average values of each compound in each replicate line were visualized using a cluster map. Clusters were classified by hierarchical clustering using the Ward method based on Euclidean distances between compound amounts^98^.

### Measurement of TAG

Relative TAG levels were measured by homogenising 10 virgin female flies aged 6–12 days and kept in non-irradiated conditions, in 1 mL of PBS (Fujifilm Wako Pure Chemicals Co.) containing 0.1% Triton X-100 (Nacalai Tesque Inc.). The homogenate was heated at 70°C for 10 minutes and then centrifuged at 20,000 × *g* for 15 minutes at 4°C. A 10-μL aliquot of the supernatant was used for TAG measurement using the LabAssay Triglyceride kit (Fujifilm Wako Pure Chemicals Co.). For protein normalization, protein concentration was measured using the Bradford method with the Protein Assay CBB Clean Up Kit and Protein Assay CBB Solution (both from Nacalai Tesque Inc.), with the standard calibration curve prepared using bovine serum albumin solution (Nacalai Tesque Inc.). Absorbance was measured using the BioTek Synergy H1 Multimode Reader (Agilent Technologies Inc., Santa Clara, CA, USA). Relative TAG levels were normalised to protein content, adjusted on the basis of the average values from UCL (Fig. 4f and Fig. 9) or non-treated UCL (Fig. 5e), and the Log2 fold change (FC) was computed accordingly.

### Measurement of reflectance and transmittance

Virgin female flies, aged 6–12 days and kept in non-irradiated conditions, were anesthetized on ice. The abdominal reflectance and transmittance were measured by capturing hyperspectral images under a BX51WI microscope (Olympus, Tokyo, Japan) fitted with an SC-108 hyperspectral imaging system (EBA Japan Co., Ltd., Tokyo, Japan).

### DNA extraction

Virgin female flies, aged 5–8 days and kept in non-irradiated conditions, were used. For Fig. 7 and Supplementary Fig. 7a, the same sample of 30 flies was used, with 10 flies used specifically for Supplementary Fig. 7bii. The flies were rinsed in a 1.5-mL tube with 99% ethanol (Fujifilm Wako Pure Chemicals Co.) for 30 seconds. Then, 180 µL of PBS and beads were added, and the flies were homogenised at 3500 rpm for 30 seconds using the Micro Smash MS100R (Tomy Seiko Co., Ltd., Tokyo, Japan) while keeping the mixture cold. DNA was extracted using the DNeasy Blood & Tissue Kit (Qiagen, Hilden, Germany) according to the kit protocols.

### 16S rRNA amplicon sequencing (V3–V4 hypervariable region)

Virgin female adult flies were used after surface sterilisation under non-irradiated conditions. For details of DNA samples, see “DNA extraction” in Methods. Quality checks were performed using the 2100 BioAnalyzer (Agilent Techologies Santa Clara, CA, USA), followed by library preparation with the NEBNext^®^ Ultra™ II DNA Library Prep Kit (Illumina, San Diego, CA, USA). DNA amplification was performed by PCR using the V3-V4 region primers 341F (5’-CCTAYGGGRBGCASCAG-3’) and 806R (5’-GGACTACNNGGGTATCTAAT-3’). PCR cycling conditions were as follows: 95°C for 3 minutes, followed by 25 cycles of 95°C for 30 seconds, 55°C for 30 seconds, 72°C for 30 seconds, with a final extension at 72°C for 5 minutes and holding at 4°C. Amplicon sequencing was conducted on an Illumina NovaSeq 6000 with the following parameters: paired-end reads of 250 bp × 2 (PE250) and an output of 50,000 reads per sample. In silico analyses were performed on a Mac OS (Catalina version 10.15) terminal using Jupyter Notebook (Python version 3.9 version 6.4.11). Quality was assessed with FastQC (version 0.11.8). Data were analysed using QIIME 2 (version 2023.7) with the Silva 138 99% OTUs full-length database (MD5: b8609f23e9b17bd4a1321a8971303310). The QIIME dada2 denoise-paired parameters were set as follows: --p-trim-left-f 30, --p-trim-left-r 30, --p-trunc- len-f 250, and --p-trunc-len-r 250. To improve classification accuracy, taxonomic filtering was performed by removing all unassigned taxa using the qiime taxa filter-table command, resulting in a refined feature table. Sequences of operational taxonomic units were classified to determine gut microbiota composition. Relative abundance calculations were performed using the qiime feature-table relative-frequency function.

### Bacterial abundance and identification

Body weights of 4 virgin adult females were measured after 3 days of non-irradiation or BL irradiation followed by a 3-day recovery period (Fig. 5a^i^–a^ii^, Fig. 6b, Supplementary Fig. 8). 4 surviving females aged 6–12 days were randomly chosen, immersed in 70% ethanol for 1 minute, washed three times with PBS homogenised in 500 μL of PBS in a 2-mL tube (stock solution), and diluted 1:100 with PBS. A 10-μL aliquot was plated onto an MRS agar plate (5.5 g MRS broth (Becton, Dickinson and Company, Franklin Lakes, NJ, USA), 1 g agar (Fujifilm Wako Pure Chemicals Co.), and 100 mL distilled water). The plates were incubated for 72 hours at 30–31°C, and colony-forming units (CFU) were counted. Single bacterial colonies were isolated, and their 16S rRNA genes were amplified by PCR using universal bacterial primers 27F (5’ AGAGTTTGATCCTGGCTCAG 3’) and 1492R (5’ GGTTACCTTGTTACGACTT 3’). PCR cycling conditions were as follows: 95°C for 3 minutes, followed by 30 cycles of 94°C for 30 seconds, 55°C for 45 seconds, 72°C for 1 minute, with a final extension at 72°C for 7 minutes and holding at 4°C. Sequencing of the amplified 16S RNA gene fragments was performed on an ABI3500 Sanger sequencing platform (Thermo Fisher Scientific) using the BigDye™ Terminator v3.1 Cycle Sequencing Kit (Thermo Fisher Scientific)^99,100^. Bacterial species were identified using Microbial Nucleotide BLAST.

### Abx treatment

The Abx was prepared according to established protocols^101^; the diet was supplemented with 100 μg/mL streptomycin sulfate (Fujifilm Wako Pure Chemicals Co.), 50 μg/mL tetracycline hydrochloride (Fujifilm Wako Pure Chemicals Co.), and 200 μg/mL rifampicin (Fujifilm Wako Pure Chemicals Co.). Flies were reared on a diet containing Abx from the embryo stage to adult.

### Capillary feeding (CAFE) assay

The assay was adapted from a previous study with minor modifications^102^. Briefly, a group of four male flies, aged 5-8 days, was kept in a plastic vial with two glass capillaries inserted (BF100-50-15, Sutter Instrument, CA, USA). The gender was selected because female flies often lay eggs inside the capillaries, which prevents precise measurement.

Flies were fed for 24h with liquid food (5% sucrose, 2% yeast extract, 0.25% nipagin in ethanol, and 0.005% Sulforhodamine B sodium salt (red dye) in water), and images of the capillaries were taken before and after the experiments. Reduction in liquid volume was quantified to estimate consumption. To account for evaporation, control capillaries without flies were included and their loss of fluid was used as a correction.

### Parental experience of BL irradiation

Virgin flies aged 1–5 days were BL-irradiated for 3 days (462 nm BL, 6.5 × 10¹⁸ photons·m⁻²·s⁻¹ for males and 10.3 × 10¹⁸ photons·m⁻²·s⁻¹ for females) and allowed to recover for 3 days. Non-irradiated parents were used as controls to ensure that the offspring had no parental experience of irradiation. Flies exposed to BL irradiation were assumed to experience BL toxicity. Parental choice was made without assessing differences in survival status, such as climbing ability, to produce offspring with different parental experiences. Randomly chosen males and females were allowed to mate and lay eggs for 4 days.

### Infection with gut contents or A. persici cocktails

In Fig. 5b, Supplementary Fig. 12a and Supplementary Fig. 13, 30 virgin female flies aged 5–8 days were immersed in 70% ethanol for 1 minute, washes three times with PBS and homogenised in 500 μL of PBS in a 2-mL tube (stock solution). The latter was diluted 1:10 with PBS. To infect with the gut microbiota, 20 μL of the diluted stock solution was added to the diet, and virgin females aged 1–4 days were maintained on this diet for 4 days. PBS was used as a control. Body weight was measured before and after infection. Climbing ability and fecundity were evaluated after 3 days of non-irradiation or BL irradiation followed by a 3-day recovery period. In Supplementary Fig. 12b, A*. persici* was cultured and isolated on MRS plates for 48 hours. These isolates, along with the standard strain of *A. persici* (JCM 25330), were cultured in 200 μL of MRS broth (5.5 g MRS broth (Becton, Dickinson and Company) and 100 mL distilled water) in 96-well plates for 72 hours. Bacterial suspensions were centrifuged to obtain pellets, and the 10-μL bacterial culture pellet was diluted 1:30 with PBS and inoculated into 20 μL of diet. Autoclaved MRS was also diluted 1:30 with PBS, and 20 μL of this mixture was used as the control.

### Whole-genome resequencing, variation estimation, variant identification, and enrichment analysis

Thirty virgin adult females were used after surface sterilisation under non-irradiated conditions. For details of DNA samples, see “DNA extraction” in Methods. Quality checks were performed using a BioAnalyzer, followed by library preparation with the NEBNext^®^ Ultra DNA Library Prep Kit (Illumina). Sequencing was conducted on an Illumina NovaSeq 6000 with the following parameters: paired-end reads of 150 bp (PE150), a data output of 10 G bases per sample, and approximately 66.7 million reads per sample. In silico analyses were performed using a macOS (Catalina version 10.15.7) terminal and Jupyter Notebook (Python version 3.9 version 6.4.11). The quality of the fastq files was assessed using seqkit stat (version 0.16.1) and then fastqc (version 0.11.8). Reads were trimmed using Trimmomatic (version 0.39) with the following parameters: LEADING:20, TRAILING:20, SLIDINGWINDOW:4:15, and MINLEN:20, based on Illumina adapter sequences (CTGTCTCTTATACACATCT, AGATCGGAAGAGCACACGTCTGAACTCCAGTCA, AGATCGGAAGAGCGTCGTGTAGGGAAAGAGTGT). Trimmed reads were re-evaluated with seqkit stat and FastQC to confirm their quality. Mapping was performed using the reference genome BDGP6.46, INSDC Assembly GCA_000001215.4 (details provided in Supplementary Data 9). Indexing and mapping were conducted using bwa (version 0.7.17), followed by conversion from SAM to BAM format using samtools (version 1.10). PCR duplicates were marked with GATK (version 4.2.6.1), and alignment statistics were calculated from the BAM file. Paired-end reads were merged using samtools, and the merged BAM file was sorted. Consensus sequences for UCL, SL, and Canton-S strains were generated using the reference genome and VCF files. Gene sequences were extracted based on BDGP 6.32.109.chr.gtf3 annotations and processed using BioPython’s SeqIO module. Gene sequence alignments between strains were performed using BioPython’s AlignIO module, MAFFT (version 7.0), and variation for each gene were calculated using Canton-S (SRX8038113)^50^ as a reference. Variation was calculated as (the number of mismatched bases to the reference gene / the total sequence length of the gene) × 100. Gaps were counted as 0.5 to account for insertions, deletions and sequencing errors. SL-specific variation was determined by subtracting the variation of each SL replicate line from the mean variation of UCL replicate lines 1-3. Genes with a mean variation of more than 5% across replicate lines 1-3 were classified as SL-specific variation. Variants were called using sambamba (version 1.0.1) and varScan (version 2.4.6) with the following parameters: sambamba mpileup, Heterozygous: SNP; pileup2snp --min-coverage 10 --min-var-freq 0.20 --*p*-value 0.05 (filter; default), Indel; pileup2indel --min-coverage 10 --min-var-freq 0.20 --*p*-value 0.10 (filter; --min-reads2 4 --min-var-freq 0.15 --*p*-value 0.05), Homozygous: SNP; pileup2snp --min-coverage 10 --min-var-freq 0.90 --*p*-value 0.05 (filter; default), Indel; pileup2indel --min-coverage 10 --min-var-freq 0.90 --*p*-value 0.10 (filter; -- min-reads2 4 --min-var-freq 0.15 --*p*-value 0.05). Homozygous variants were identified by extracting rows with matching Cons and VarAllele values. Common chromosomes and positions across replicates (lines 1, 2, and 3) within each strain were defined to identify strain-specific shared homozygous variants. Shared and unique homozygous variants among UCL, SL, and Canton-S^102^ were analysed. homozygous variants were annotated using SnpEff (version 5.0) and visualized using IGV (version 2.8.12). Variants classified as having a low impact (“LOW”) by SnpEff were excluded from the analysis list (full list available in Supplementary Data 11). Enrichment analysis was conducted with DAVID (version 2023q4) and KEGG (version 108.1).

### RNA extraction

Ten virgin female flies, aged 5–8 days and kept in non-irradiated conditions, were frozen in liquid nitrogen and homogenised in Trizol (Molecular Research Center, Inc., Cincinnati, OH, USA). Chloroform (Fujifilm Wako Pure Chemicals Co.) was then added, and the mixture was vortexed twice and centrifuged at 20,000 × *g* for 10 minutes at 4°C. The upper aqueous phase was transferred to a new tube, and 200 µL of isopropyl alcohol was added. The mixture was vortexed, incubated at room temperature for 10 minutes and centrifuged again at 20,000 × *g* for 10 minutes at 4°C; the supernatant was discarded. To wash the RNA pellet, 400 µL of 75% ethanol (prepared with DEPC-treated water) was added, followed by vortexing. The sample was then centrifuged at 20,000 × *g* for 5 minutes at 4°C. The supernatant was discarded, and the pellet was centrifuged once more for 1 minute.

The pellet was air-dried and resuspended in DEPC-treated water.

### RNA-sequence and enrichment analysis

Quality assessment was initially performed using a BioAnalyzer. Library preparation was conducted with the NEBNext Ultra II Directional RNA Library Prep Kit (Illumina) and the NEBNext Poly(A) mRNA Magnetic Isolation Module (New England Biolabs, Ipswich, MA, USA). Sequencing was performed on an Illumina NovaSeq 6000, with a read length of PE150 (150 bp × 2 paired-end), generating a data output of 6 G bases per sample and approximately 40 million reads per sample. In silico analyses and read trimming followed the protocols described in the "Whole-genome resequencing, variant identification, and enrichment analysis" section in Methods. Poly(A) tails were removed using cutadapt, and mapping was conducted with hisat2 (version 2.2.1) using the BDGP6.46 reference genome (INSDC Assembly GCA_000001215.4) (for details, see Supplementary Data 13). Gene-wise read counting was accomplished using HTSeq (version 2.0.4) with the dmel-all- r6.54.gtf annotation file. Read counts were normalised with edgeR (version 3.40.2) using the trimmed mean of M values method. Subsequent principal component analysis and differential expression analysis were carried out with edgeR and scikit-learn (version 1.0.2). DEGs were identified by comparing non-treated SL with non-treated UCL, abx-treated SL, or abx-treated UCL groups, applying a false discovery rate–adjusted *p*-value threshold of <0.05. Python libraries such as pandas (version 2.0.3), numpy (version 1.25.2), scipy (version 1.11.2), and stats were used to filter DEGs with *p-*values below 0.05. For the DEGs, log2 FC values and Z scores normalised within the range of −2 to 2 were calculated. These Z scores were then used to create a cluster map using the Ward method, which classified genes and treatments on the basis of their similarity patterns. Enrichment analysis was performed using DAVID and KEGG. Gene lists related to specific biological functions were retrieved from KEGG (for TCA cycle, oxidative phosphorylation and Toll and Imd signaling pathways) and FlyBase (for lipid metabolic processes). In Fig. 8c and Supplementary Fig. 18d, heatmaps illustrating functional profiles were generated using log2 FC values. These values were compared for each gene and each group and normalised against the mean value of the non-treated UCL group.

### Genetics and evaluation of TAG levels and BL tolerance

We collected genetically engineered flies with known elevated TAG levels and reared them at 23–25°C under a 12-hour light/dark cycle. Using the GAL4/UAS system and null mutants, we generated flies with RNAi knockdown or overexpression of target genes. The following strains from the Bloomington *Drosophila* Stock Center were used (see Supplementary Data 15): Null mutants, *w;Akh* (84448), w;*Bmm* (15828), *yw,Pdgy* (19990), *yw;Chico* (14337), *w;InR* (8267), *w;Ilp 2 3 5* (30889), *w;Lk6* (87087); GAL4 drivers, *yw;Lsp2*(6357), *w;ppl* (58768), *w;Tk* (51973); UAS strains, *yv;UAS-AkhR^RNAi^* (51710), *yv;UAS- Pect^RNAi^*(63710), *yv;UAS-Pten^RNAi^* (33643), *yv;UAS-Bmm^RNAi^*(25926), *y sc v sev;UAS- Stim^RNAi^* (52911), *y sc v sev;UAS-Lsd-1^RNAi^* (65020), *yv;UAS-Tk^RNAi^* (25800), *yw;UAS-Gbb* (63059), *yw;UAS-InR* (8263), *w;UAS-Lsd-2* (98116), *w;UAS-Lk6* (8709), *w,UAS-Rpr1* (5823). We used F1 progenies for the experiments, obtained by crossing GAL4 driver males with UAS effector females. As controls, we used wild-type Canton-S and w1118 genetically washed by crossing with the Canton-S flies, F1 progenies from crosses of w1118 with UAS effectors, and F1 progenies from crosses of GAL4 drivers with *y sc v;eGFP^RNAi^* (41555) and *y sc v;eGFP^RNAi^* (41556) UAS effectors. Virgin female flies aged 1– 5 days were irradiated with either darkness or BL for 3 days and recovered for 3 days.

Their survival rates were assessed, and 5 flies were randomly chosen to mate with 5 Canton-S males for 4 days (for details, see “Measurement of fecundity” in Methods). TAG storage was measured in virgin female flies aged 7–12 days. Relative TAG levels are presented as log2 FC relative to the mean values across all treatments. Fecundity was calculated as log2 FC relative to the non-irradiation condition for each treatment. Data are from SL and UCL (replicate lines 1, generation 57).

### Phylogenetic tree based on the 16S rRNA gene

16S rRNA gene sequences were obtained from the Integrated Microbial Genomes (IMG) database, with the addition of *A. persici* DS-1 strain^66^ for *A. persici*. Based on previous studies, *Neokomagataea thailandica* and *Swingsia samuiensis* were used as outgroups^103^. Sequence lengths ranged from 1,391 to 1,498 bp. The phylogenetic tree in Supplementary Fig. 19 was constructed by including the *A. persici* sequence (397 bp) from the dominant OTU ID, which accounted for 89-98% of the microbiome analysis in Supplementary Fig. 7a. Based on these results, the phylogenetic position of *A. persici* in our laboratory was inferred in Fig. 10a. Phylogenetic analyses were performed using MEGA (version 11.013), with alignments performed using ClustalW. A maximum likelihood (ML) method with 1,000 bootstrap replicates was used (substitution type: general time reversible model; ML heuristic method: Nearest-Neighbour Interchange; initial tree for ML: NJ/BioNJ). The resulting phylogenetic tree was visualised using iTOL (version 7.0).

***A.persici* circular genome visualisation and transcriptome analysis**

The complete genome of *A. persici* (GCA_002006565.1_ASM200656v1) was used for circular genome visualisation and transcriptome analysis for reference genome. The circular genome representation was generated using GenoVi (version 0.2.16)^104^. To isolate the microbial reads, we extracted the sequences that remained unmapped in the host transcriptome analysis (for the initial mapping, see “RNA-sequence and enrichment analysis” in Methods). These reads were filtered from the SAM files using Samtools (samtools view -b -f 4) and converted to FASTQ format. Quality was assessed using FastQC, and overrepresented sequences not corresponding to *A. persici* were removed.

The resulting reads were trimmed using Trimmomatic (LEADING:20, TRAILING:20, SLIDINGWINDOW:4:15, MINLEN:20). A second alignment to the host genome was then performed using HISAT2 to further eliminate host-derived sequences. The final unmapped reads were aligned to the complete *A. persici* genome. Genome indices for both host and *A. persici* were constructed using HISAT2, and all alignments were performed using the same parameters: hisat2 -x reference genome index -1 -2 -k 3 --mp 2.1 --np 1 --rdg 10.5 -- rfg 10.5 --seed 42 --score-min L,0,-0.9) (for details, see Supplementary Data 16). Gene expression was quantified using featureCounts (subread-1.5.2.-MacOSX-x86_64, version 2.0.8) and parameters: featureCounts -t CDS -g gene_id -a genomic.gft (GCA_002006565.1) -M -p. Gene expression levels were quantified using transcripts per million (TPM). Reads per kilobase (RPK) were calculated as expression / (length / 1000). TPM values were then derived by normalising the RPK of each gene to the total RPK within its respective GeneID. Sequence depth was estimated using the formula (total bases mapped to the gene) / (target gene length). DEGs were identified by comparing the UCL and SL groups using an FDR adjusted *p*-value threshold of <0.05. To quantify changes in gene expression, we calculated the log2 FC of the transcripts. The mean TPM was calculated for each gene in the UCL group, and the fold change was determined as the ratio of individual sample TPM values to this mean. The dataset was then log2 transformed. Enrichment analysis was performed using ShinyGO (version 0.82) and Gene ontology (GO) biological process (version Ensembl 92). Gene lists related to oxidation-reduction process was retrieved from GO biological process. The genomic.gft (GCA_002006565.1), UniProt (version 2025_01) and KEGG databases were used to determine protein identities, enzyme classifications and functional annotations for each gene. For details of the host transcriptome analysis and visualisation, see “RNA-sequence and enrichment analysis” in Methods.

### Growth experiments of isolated *A. persici*

*A. persici* was isolated from 10 adult females and cultured on MRS plates for 48 hours. 50 isolates were pre-cultured in 200 μL MRS broth (5.5 g MRS broth and 100 mL distilled water) in 96-well plates for 72 hours. Bacterial growth was then assessed by culturing in 50 mL of MRS broth with continuous shaking at 60 rpm. Cultures were maintained at 30-31°C throughout the incubation period. Optical density (OD) was measured using a compact rocking incubator equipped with a spectrophotometer (TVS062CA ADVANTEC Tokyo, Japan).

### Statistical analysis

#### State space model and Bayesian estimation

The state space model is a statistical framework designed to analyse time series data with autocorrelation, based on state and observation equations^105–107^. It was applied to estimate fecundity across generations, incorporating replicate lines as random effects. In Fig. 2a, intergenerational fecundity data were analysed using a Bayesian state space model implemented with pystan (Python 3.9, version 2.17.0.0). This model included the following components: Data block; Defined the number of generations (length of the time series data), a vector of explanatory variables, and three outcome variables corresponding to each replicate line, each following a normal distribution. Parameters block; Estimated a vector of levels with drift components (*mu*), vectors of random effects (*r1*, *r2*, *r3*), coefficients of the explanatory variables (*b1*, *b2*, *b3*), the standard deviation representing the magnitude of the drift component (*s_z*), and the standard deviations of the random effects (*s_r1*, *s_r2*, *s_r3*). Transformed parameters block; Calculated the expected values for each outcome variable (*lambda1*, *lambda2*, *lambda3*) as the sum of the level with drift component (mu), the product of the explanatory variables’ coefficients (*b*) and values (*ex*), and the random effects (*r*). Model block; Assumed that the random effects followed a normal distribution with a mean of 0 and a given standard deviation. The state equation, *mu[i] ∼ normal(2 * mu[i-1] - mu[i-2], s_z*), expressed changes in the level with drift components using a second-order local linear regression. The observation equations, *y1[i] ∼ normal(lambda1[i], sigma1), y2[i] ∼ normal(lambda2[i], sigma2)*, and *y3[i] ∼ normal(lambda3[i], sigma3)*, defined the observed values as following a normal distribution. Generated quantities block; Sampled the predicted values for each outcome variable using the normal distribution. Random numbers were generated according to the posterior distribution using the Markov Chain Monte Carlo (MCMC) method with parameters set to iter=10000, chains=4, and control={’adapt_delta’: 0.8, ’max_treedepth’: 20}. Convergence was confirmed by ensuring that the *R*hat of the predicted values was <1.10 and by examining the trace plots (for details, see Supplementary Data 2).

#### Hierarchical Bayesian model

Unless otherwise specified, hierarchical uncertainty estimation, accounting for random effects, was performed using Hierarchical Bayesian Model (HBM)^108^. The model specification and execution used the following tools and versions: Software; R (version 4.0.1), rstan (version 2.32.2, Stan version 2.26.1), brms (version 2.20.4), and bayesplot (version 1.10.0). Bayesian model comparison; The Bayes factor (BF10) quantifies the strength of evidence for the alternative hypothesis (H1) over the null hypothesis (H0)^109^. A BF10 > 1 indicates support for H1, whereas a BF10 < 1 favours H0. Guidelines for interpretation are as follows: BF10 > 3 indicates moderate evidence, BF10 > 10 indicates strong evidence, and BF10 > 100 provides conclusive support for H1. Conducted using BF10 with the loo function, computing *exp(0.5 * (elpd1 - elpd2))*, where elpd1 denotes the model’s expected log pointwise predictive density (elpd_loo) and elpd2 denotes that of the null model. MCMC sampling; Parameters were set to seed=1, chains=4, iter=20000, warmup=5000, and thin=4. If convergence was not attained, adjustments were made to iter=25000 and warmup=10000, ensuring that *R*hat values remained below 1.10.

Prior distributions; Uniformly set to Gaussian. Model details; Comprehensive details including the response variable, predictors, random effects, and estimation results are available in Supplementary Data 1, Supplementary Data 3, 4, Supplementary Data 6–8, Supplementary Data 16. Results included posterior median estimates, Bayesian 95% credible intervals (CI), and approximate *p*-values (*p*MCMC) computed from posterior sampling. Posterior sampling and *p*MCMC calculation; To test whether the posterior distribution of the difference in each treatment was above or below 0, we followed references^110–113^. Posterior samplings of each parameter were stored in diff_samples, and pairwise comparisons of treatments were made using the formula: *p*MCMC = 2 * min(mean(diff_samples > 0), mean(diff_samples < 0)).

#### Logistic function

*Lethal dose*, the survival rate at each dose was calculated by assigning a value of 1 if the survival time exceeded the median survival time of UCL at 10 × 10¹⁸ photons·m⁻²·s⁻¹ (88 h) and 0 otherwise. The LD50 was estimated using a logistic function: The logistic function, defined as *f*(*x*) = 100 / (1 + exp(*slope* × (*x* – LD50))), describes the survival probability as a sigmoidal function of dose. *Bacterial growth curves*, modelled using a logistic function defined as *f*(*t*) = *K* / (1 + exp(-*r* × (*t* - *t₀*))), where *K* is the maximum optical density (OD), *r* is the growth rate, and *t₀* is the midpoint of the growth phase. Parameter estimation was performed by fitting this function to the OD measurements over time.

#### Correlation and Regression analysis

The Pearson correlation coefficient or linear regression and 95% confidence intervals were calculated using the pearsonr function or linregress function from the scipy.stats library.

## Data availability

16S rRNA amplicon sequencing, whole-genome resequencing, and RNA-sequence data have been deposited in the DNA Data Bank of Japan (DDBJ). All raw read data have been uploaded to DDBJ under BioProject PRJDB19030, PRJDB19027, and PRJDB19028. The source data for all figures are available in the Supplementary Data 17.

## Code availability

The code for all analyses and visualisations is available on reasonable request from the corresponding author Y.T.

## Acknowledgements

We are grateful to Dr. Takema Fukatu, Dr. Yuto Yoshinari, Dr. Takashi Nishimura, and Dr. Koichiro Tamura for their valuable discussions regarding the study. We also thank Dr. Masaki Iwata, Dr. Shuhei Miyashita, Dr. Chisato Kobayashi, Dr. Katsuya Taniyama, Dr. Takashi Makino, Dr. Yuma Takahashi, Dr. Takahisa Ueno, Dr. Masashi Murakami, Dr. Misako Okumura, Dr. Takahiro Chihara Dr. Toshinori Hayashi, Dr. Takeshi Igawa, Dr. Ritsuko Suyama and Dr. Sota Tanaka for their important suggestions. We would like to thank Dr. Atsuhiko Nagasawa and Dr. Yuki Chiba for teaching us the GC-MS analysis, and Dr. Fu Namai for teaching us microbiome analysis. We also thank Mr. Yusaku Hayashi for instructing us on how to dissect the fat body. Ms. Ayako Abe and Mr. Kento Fukuoka provided technical assistance. We extend our gratitude to Dr. Haruki Kitazawa and Dr. Shoichiro Kurata for lending us their laboratories. We thank the Rhelixa, Inc. for conducting the next-generation sequencing and assisting with the data acquisition, ELSS, Inc for proofreading and editing the English language of this manuscript, Bloomington Drosophila Stock Center for providing the Drosophila stocks used in this study, and Japan Collection of Microorganisms RIKEN BioResource Research Center for providing the bacterial strains used in this research. This work was supported by JST SPRING, grant numbers JPMJSP2102 and JPMJSP2114 to Y.T.

## Author information

Authors and Affiliations

Graduate School of Agricultural Sciences, Tohoku University, Sendai, Japan

Yuta Takada, Wakako Ikeda-Ohtsubo, and Masatoshi Hori

Frontier Research Institute for Interdisciplinary Sciences, Tohoku University, Sendai, Japan

Toshiharu Ichinose

Graduate School of Pharmaceutical Sciences, Tohoku University, Sendai, Japan

Naoyuki Fuse

Graduate School of Life Sciences, Tohoku University, Sendai, Japan

Toshiharu Ichinose, Kokoro Saito, and Hiromu Tanimoto

## Author contributions

Y.T. conceived and designed the study, conducted all the experiments, analysed all the data and wrote the original, second, third, fourth and final versions of the manuscript. Y.T. and M.H. designed laboratory selection experiment. Y.T. and W.I. designed and conducted the infection experiment and bacterial assays. Y.T., T.I., and N.F. performed the whole-genome and RNA sequencing. Y.T., K.S., T.I., and H.T. designed the genetic and CAFE assays. Y.T. and K.S. performed the CAFE assay. Y.T., W.I., N.F., and T.I. edited the first draft of the manuscript. Y.T. and T.I. edited the second, third, and fourth versions. H.T. and M.H. supervised the study. All authors participated in discussions and reviewed the manuscript.

## Supplementary figures

**Supplementary Fig. 1:**
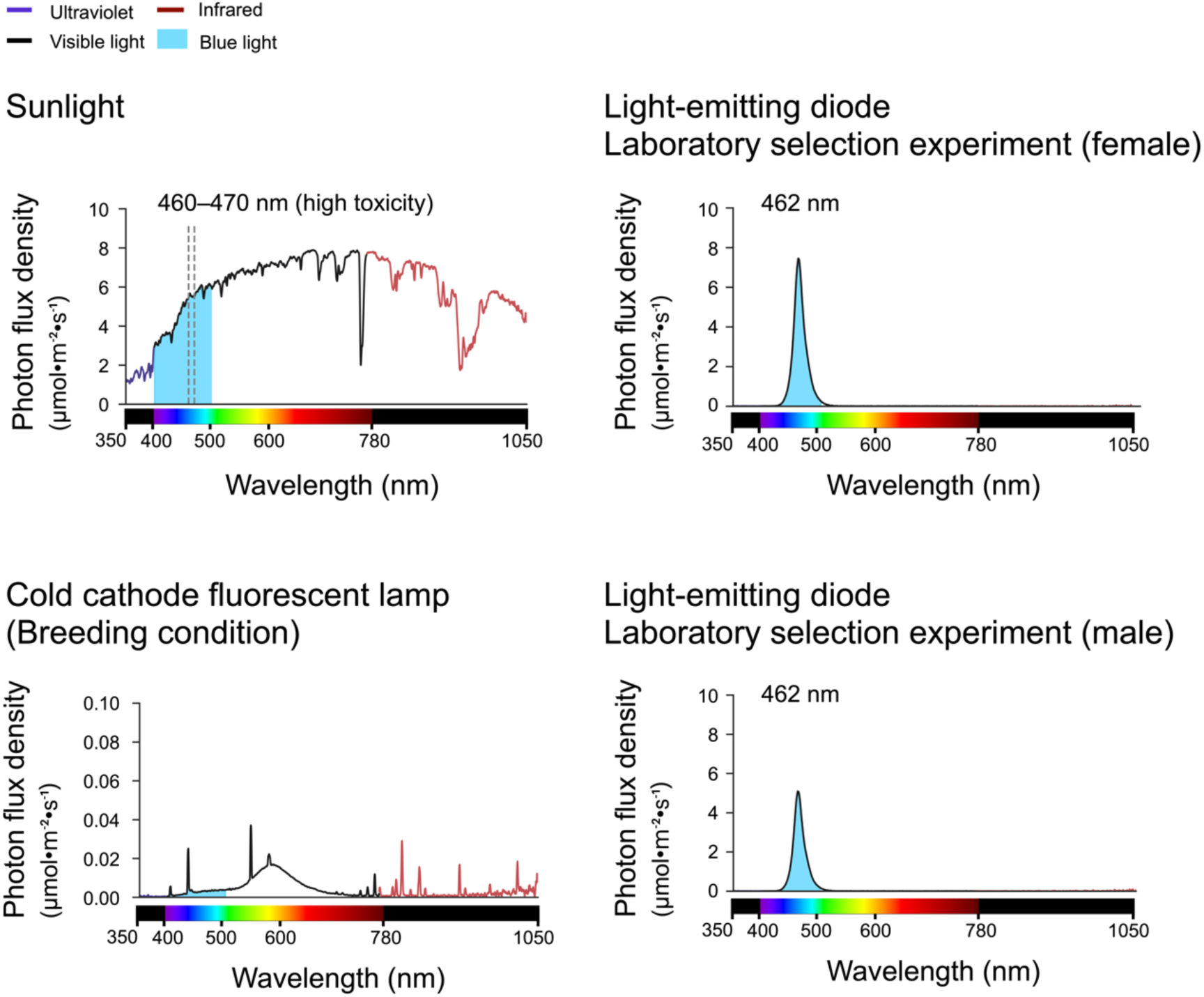
Sunlight contains a significant amount of blue light. Sunlight photon flux density: Sendai, Japan; 38°N, 140°E, clear sky; November 21, 2021; 11:30 AM (approximately 4000 μmol·m⁻²·s⁻¹, blue light (BL): approximately 470 μmol·m⁻²·s⁻¹). 460–470 nm is a highly toxic wavelength range for *Drosophila* adults. Laboratory selection experiment (female) light condition photon flux density: approximately 175 μmol·m⁻²·s⁻¹, BL: approximately 175 μmol·m⁻²·s⁻¹ (10.3 × 10¹⁸ photons·m⁻²·s⁻¹)). Laboratory selection experiment (male) light condition photon flux density: approximately 120 μmol·m⁻²·s⁻¹, BL: approximately 120 μmol·m⁻²·s⁻¹ (6.5 × 10¹⁸ photons·m⁻²·s⁻¹)). Breeding light condition photon flux density: approximately 3 μmol·m⁻²·s⁻¹, BL: approximately 0.3 μmol·m⁻²·s⁻¹. For details, see Supplementary Data 17 (Light condition and Supplementary Fig. 1).

**Supplementary Fig. 2:**
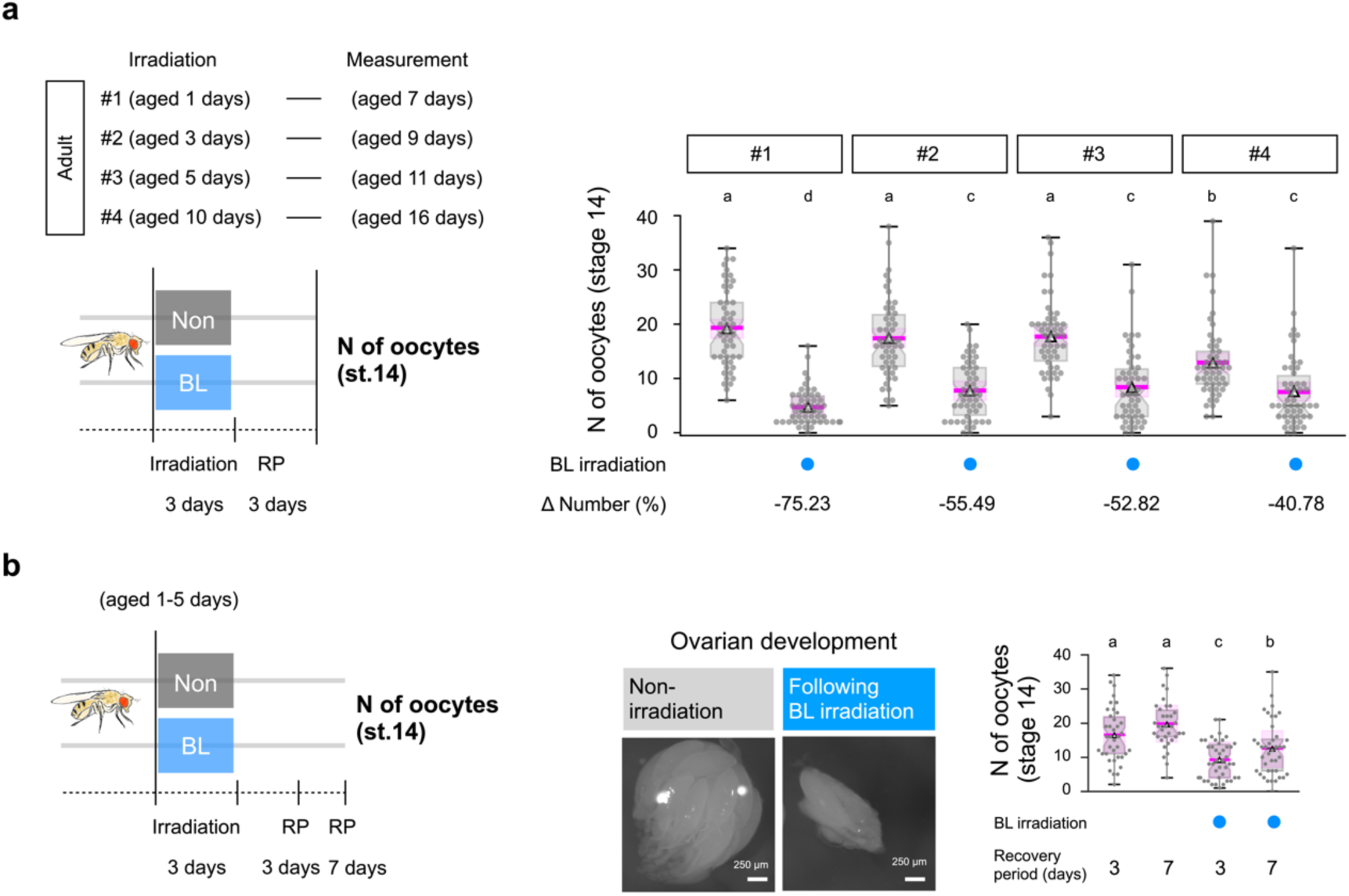
Dramatic impact on fitness due to BL toxicity a, Impact of BL toxicity on flies (Canton-S strain) of different ages subjected to a single irradiation for 3 days and a recovery period (RP) of 3 days. *n* = 50, Bayes factor 10 (BF10) > 100. Δ Number is percent change in same age non-irradiation vs following BL irradiation. **b**, Impact of BL toxicity on UCL flies (generation 34) subjected to a single irradiation for 3 days: ovarian development, the number of stage 14 oocytes after recovery. *n* = 34–42. BF10 > 100. Statistical analyses for panels **a** and **b** were performed using Hierarchical Bayesian Model (HBM) with Markov Chain Monte Carlo (MCMC) methods, posterior median, and Bayesian 95% credible intervals (CI). The magenta, bold lines and shaded areas show the posterior median and the Bayesian 95% CI. In **a** and **b**, different letters indicate significant differences at *p*MCMC < 0.05. For details, see Supplementary Data 1 for all results, including statistical models.

**Supplementary Fig. 3:**
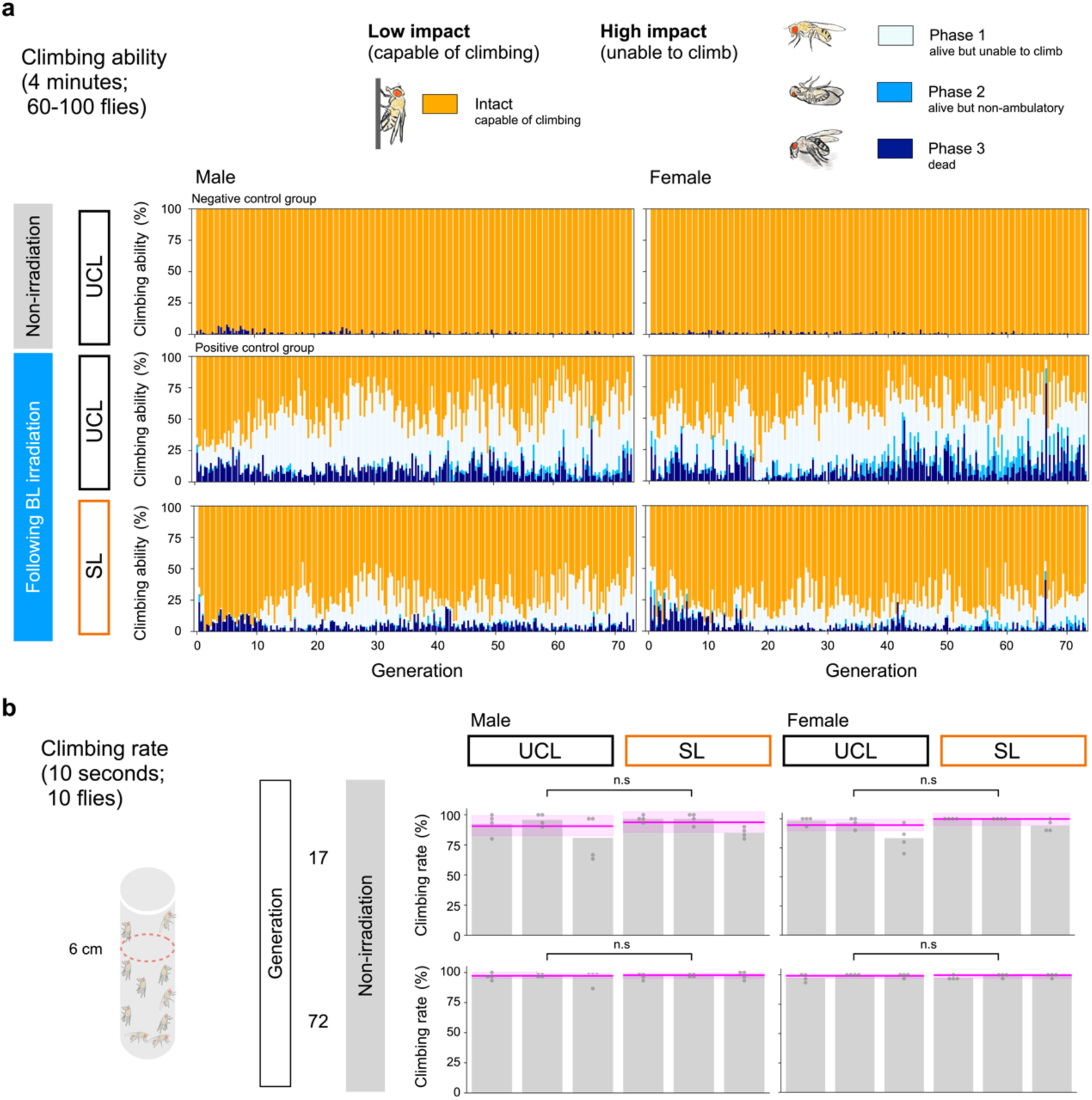
SL flies are BL tolerant. **a**, Climbing ability and generational changes in male and female flies after exposure to BL irradiation. Climbing ability is represented by a total of 60–100 flies per bar, with bars representing replicate lines 1, 2, and 3. All categories were exposed to BL except the negative control group (top). Negative control group: The UCL flies were kept in darkness for 3 days instead of BL irradiation. Positive control group: A subset of UCL flies in each generation was exposed to BL under the same conditions as SL. **b**, Climbing rate under non-irradiated conditions. The percentage of 10 flies that crossed the 6 cm reference line within 10 seconds of tapping. Each measurement was repeated three times, and the average was taken as the climbing rate. *n* = 12 (corresponding to 120 flies). Statistical analyses for panels **b** were performed using a HBM with MCMC methods, posterior median, and Bayesian 95% CI. The magenta, bold lines and shaded areas show the posterior median and the Bayesian 95% CI. n.s = not significant. For details, see Supplementary Data 1 for all results, including statistical models. The Supplementary Data 17 (Light condition) contains all the light condition measurements used in the experiment.

**Supplementary Fig. 4:**
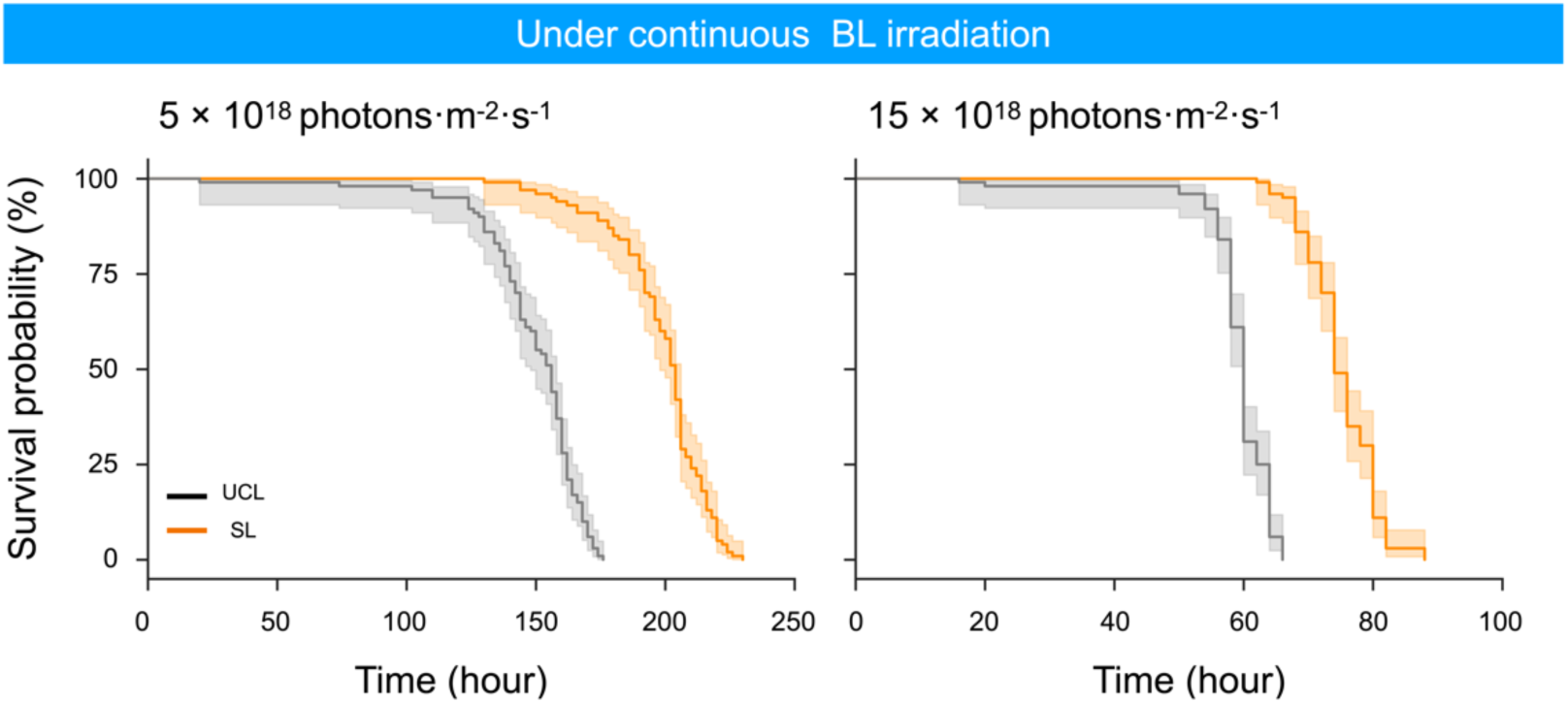
The 73rd generation SL flies show high survival under BL irradiation. Lifespan under continuous BL irradiation. Kaplan–Meier survival curve, 5 × 10¹⁸ photons·m⁻²·s⁻¹, Female *n* = 100, BF10 >100. Estimates: UCL 149.25 h (145.09–153.11), SL 198.65 h (194.71–202.77). 15 × 10¹⁸ photons·m⁻²·s⁻¹, Female *n* = 100, BF10 >100. Estimates: UCL 59.12 h (58.00–60.31), SL 75.28 h (74.01–76.57), *p*MCMC < 0.001. Replicate line 1, generation 73. For details, see Supplementary Data 3 for all results, including statistical models.

**Supplementary Fig. 5:**
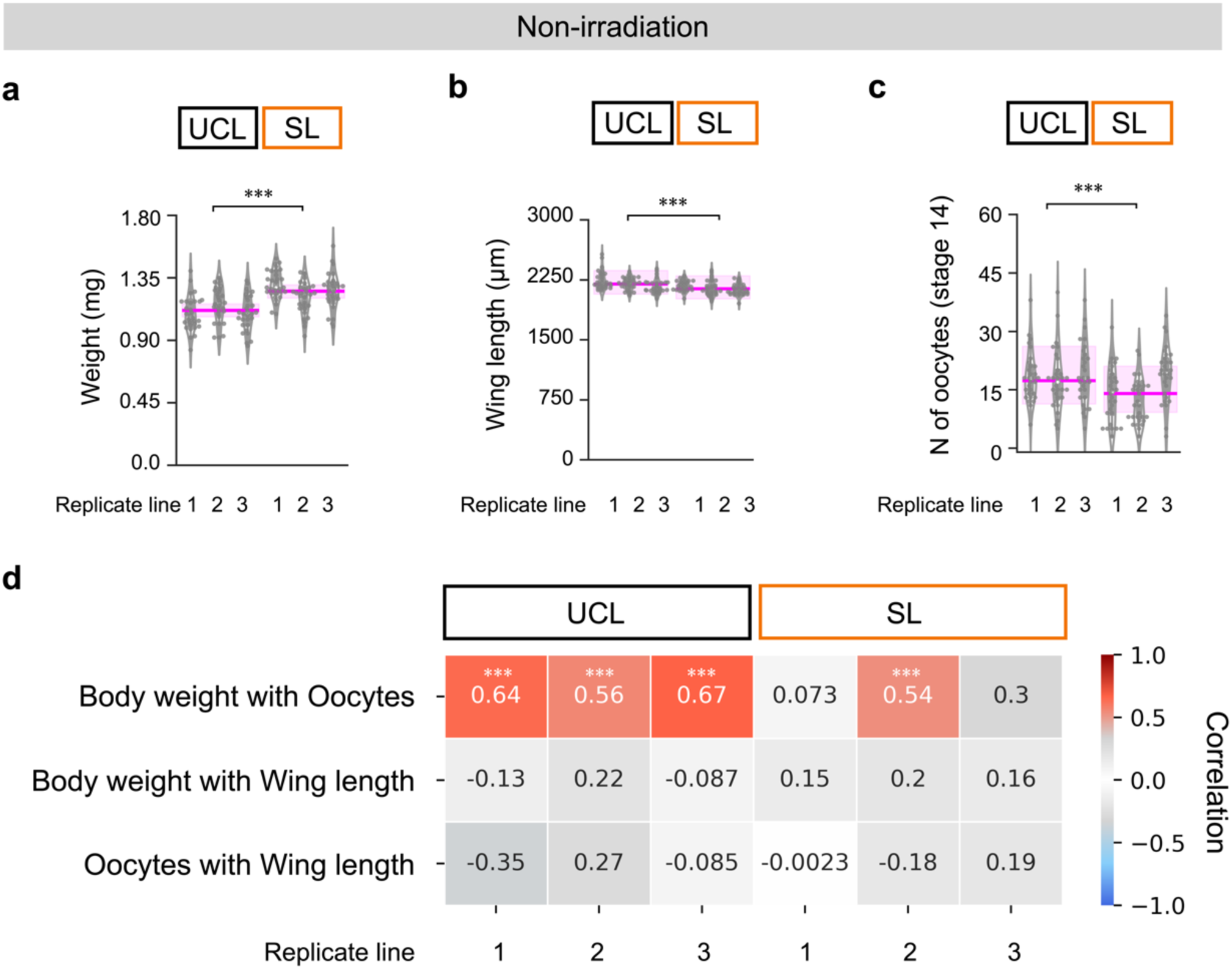
SL flies have heavier bodies, shorter wings and fewer oocytes. **a**, Body weight. *n* = 90, BF10 > 100; generation 28. **b**, Wing length. *n* = 90, BF10 = 91.1; generation 28. **c**, Number of stage 14 oocytes. *n* = 90, BF10 = 2.46; generation 28. **d,** Correlation between traits, based on data from panels **a**–**c**. *n* = 30. Correlation coefficients and *p*-values were calculated using the Pearson correlation method. Asterisks indicate significance: *** *p* < 0.001. Statistical analyses for panels **a**–**c** were performed using a HBM with MCMC methods, posterior median, and Bayesian 95% CI. The magenta, bold lines and shaded areas show the posterior median and the Bayesian 95% CI. Asterisks indicate significance levels: *** *p*MCMC < 0.001. See Supplementary Data 4 for details of all results, including statistical models.

**Supplementary Fig. 6:**
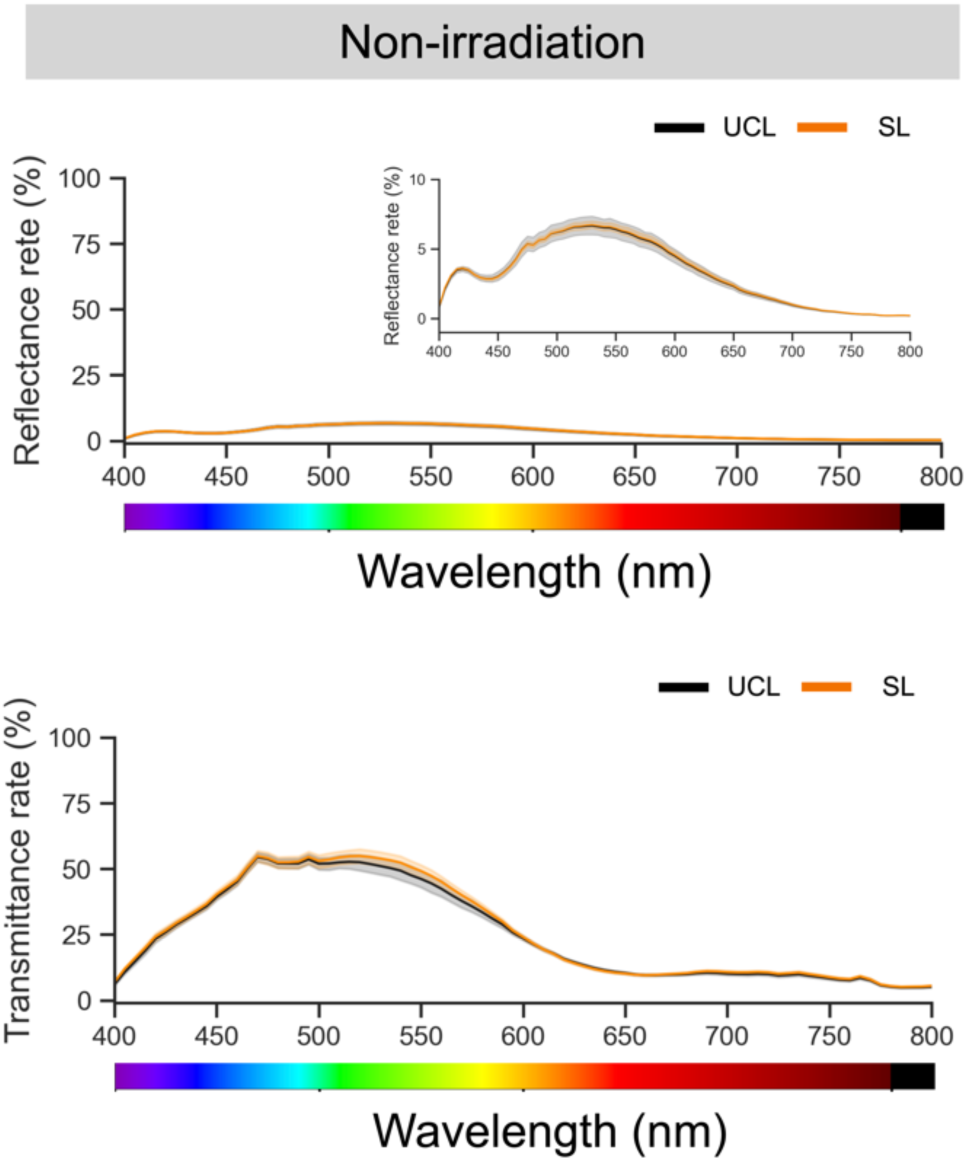
No differences in light reflection or transmission due to physical characteristics. Reflectance rate and transmittance rate in female abdomens, *n* = 16; replicate line 1, generation 49. The shaded areas show the 95% confidence interval for variation between samples.

**Supplementary Fig. 7:**
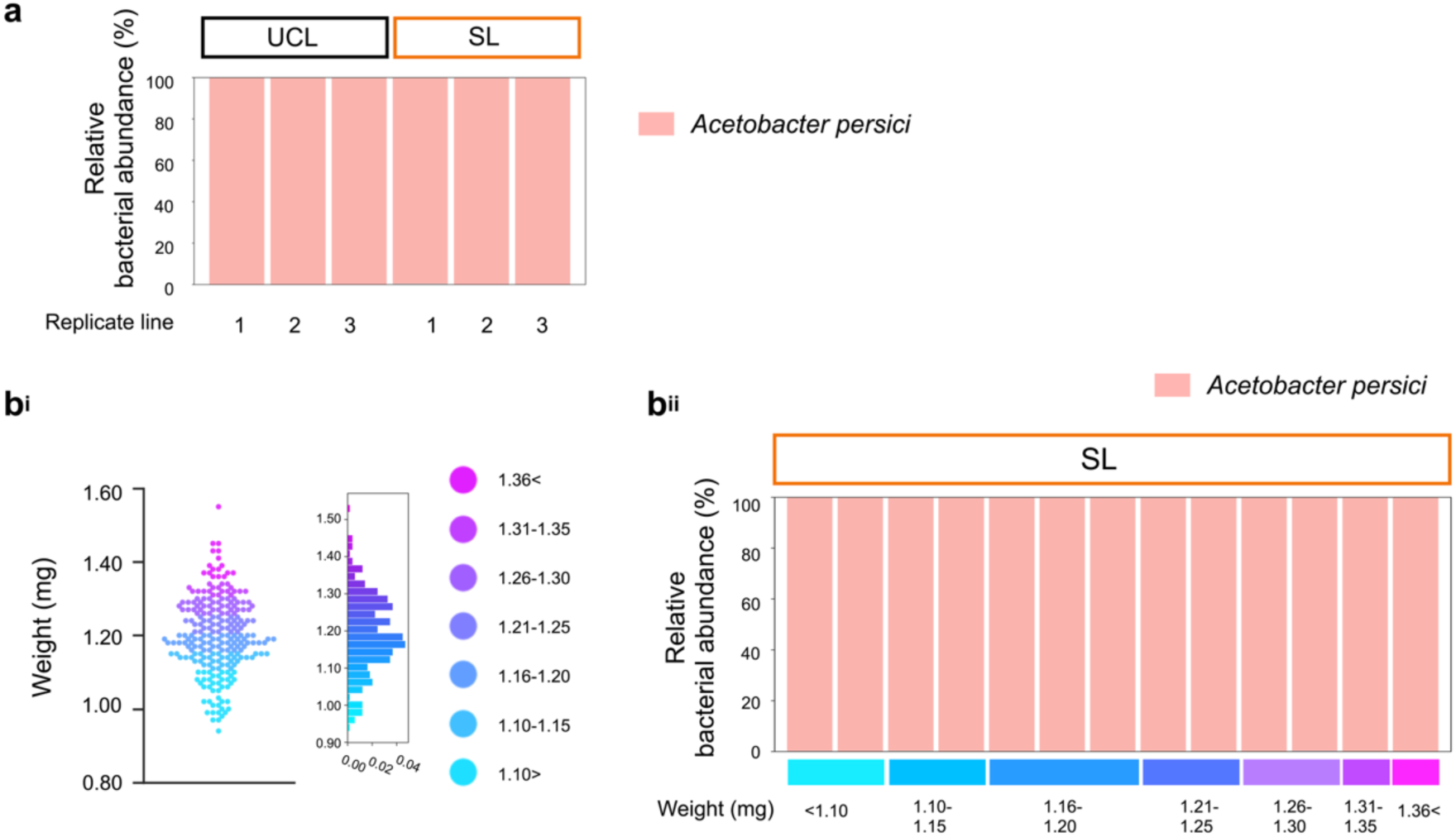
The gut microbiota of SL flies consists of a single *Acetobacter* species. **a**, Relative bacterial abundance based on 16S rRNA amplicon sequencing in SL and UCL. The DNA samples used were the same as those used for whole-genome resequencing in Fig 6 (generation 44). **b^i^**, Overview of body weight distribution in the SL flies. replicate line 1, *n* = 240; generation 45. **b^ii^**, Relative bacterial abundance based on 16S rRNA amplicon sequencing in SL replicate line 1, generation 45. For details, see Supplementary Data 5.

**Supplementary Fig. 8:**
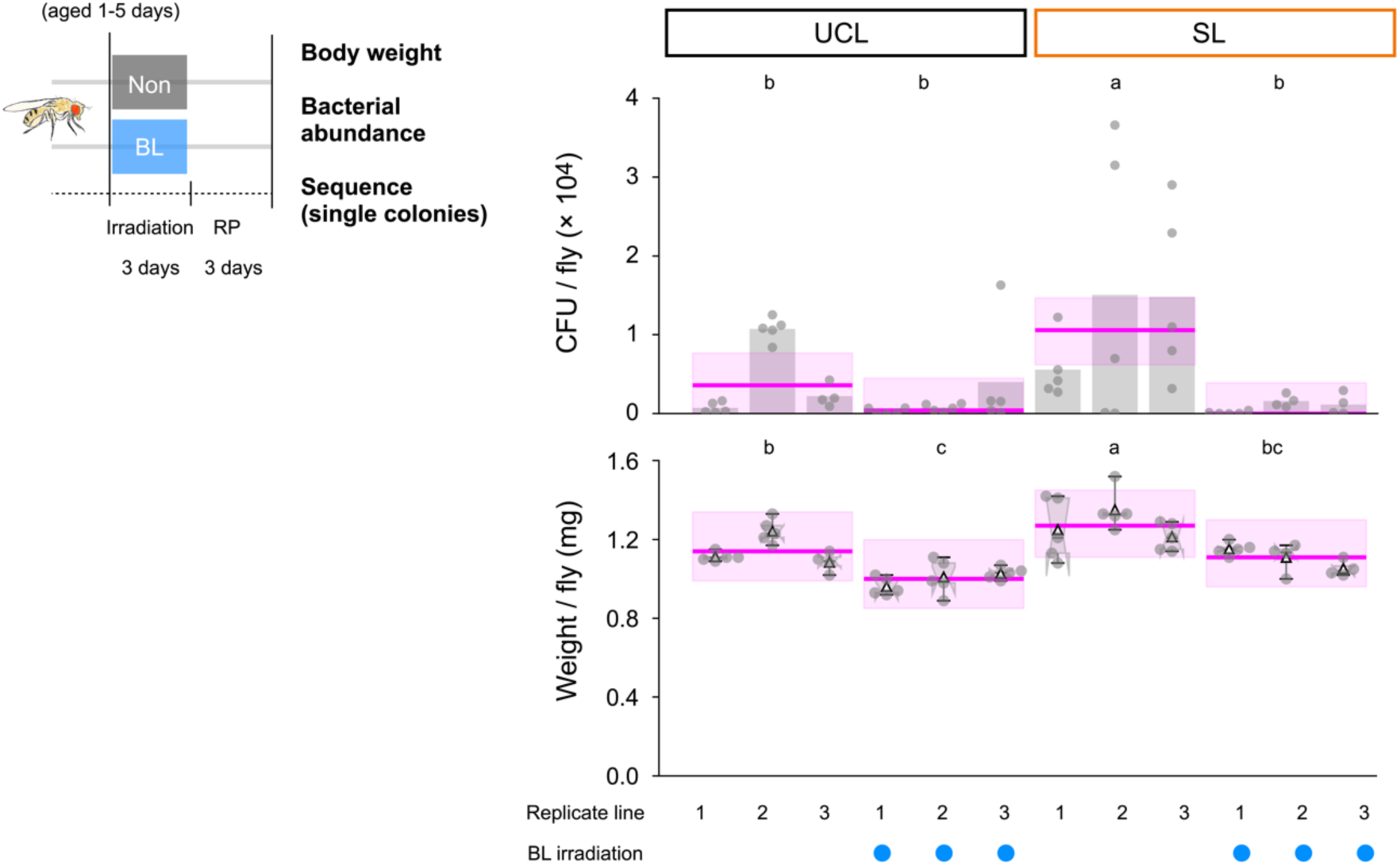
Dramatic reduction in gut microbiota and body weight of *Drosophila* after BL irradiation. **a**, Bacterial abundance and Body weight, *n* = 13–15, BF10 >100; generation 38. Randomly picked colonies were sequenced for 16S rRNA genes and all bacteria were identified as *A. persici.* Statistical analyses were performed using HBM with MCMC methods, posterior median, and Bayesian 95% CI. The magenta, bold lines and shaded areas show the posterior median and the Bayesian 95% CI. Different letters indicate significant differences at *p*MCMC < 0.05. See Supplementary Data 6 for details of all results, including statistical models.

**Supplementary Fig. 9:**
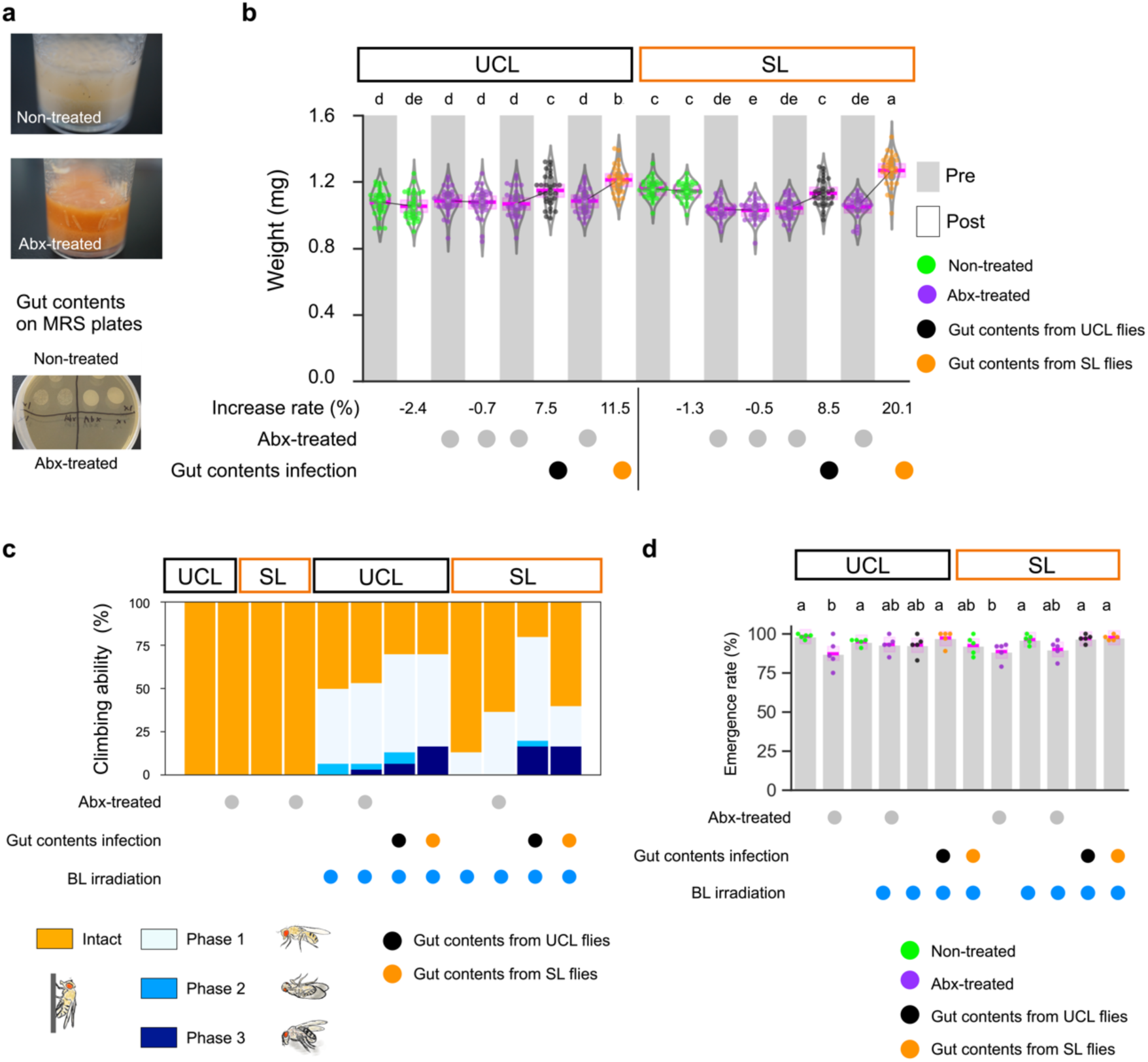
The unique phenotypes of SL flies depend on gut microbiota. **a**, Effect of antibiotic mixture (Abx) treatment on gut microbiota. **b**, Impact of Abx treatment and infection with gut contents on body weight. Grey bars represent pre-infection, while white bars indicate post-infection after 4 days. *n* = 30, BF10 > 100; replicate line 1, generation 47. **c**, Climbing ability following BL irradiation. Each bar represents a total of 30 flies; replicate line 1, generation 47. **d**, Emergence rate, based on data shown in Fig 5b. BF10 = 1.30; replicate line 1, generation 47. Statistical analyses for panels **b** and **d** were performed using HBM with MCMC methods. The magenta, bold lines and shaded areas show the posterior median and the Bayesian 95% CI. Different letters indicate significant differences at *p*MCMC < 0.05. See Supplementary Data 7 for details of all results, including statistical models.

**Supplementary Fig. 10:**
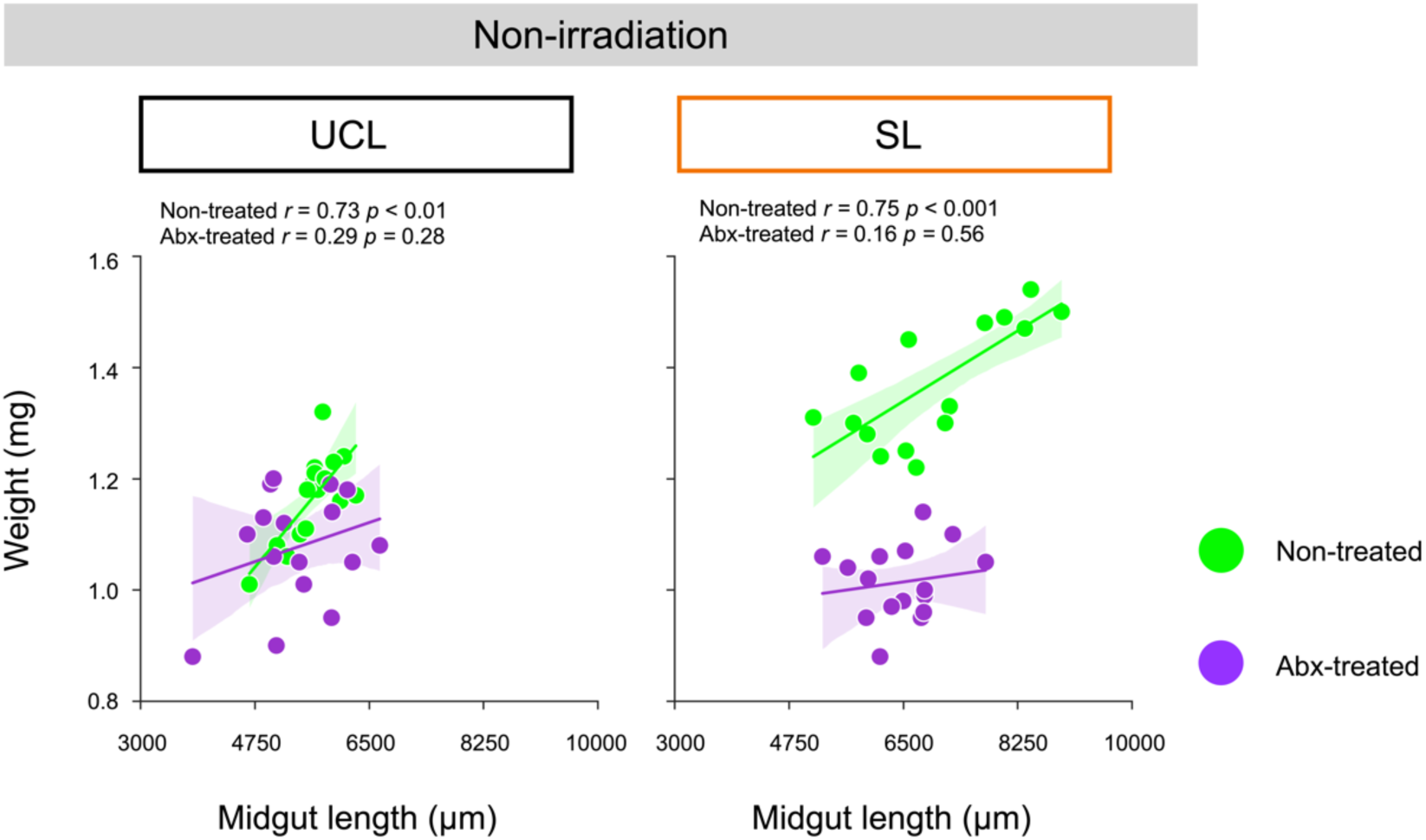
*Drosophila* body weight and midgut length positively correlate in the presence of gut microbiota.Correlation between body weight and midgut length, based on data from Fig. 5f. *n* = 16, replicate line 1, generation 48. The shaded areas show the 95% confidence interval for variation between samples. Correlation coefficients and *p*-values were calculated using the Pearson correlation method. For details, see Supplementary Data 7.

**Supplementary Fig. 11:**
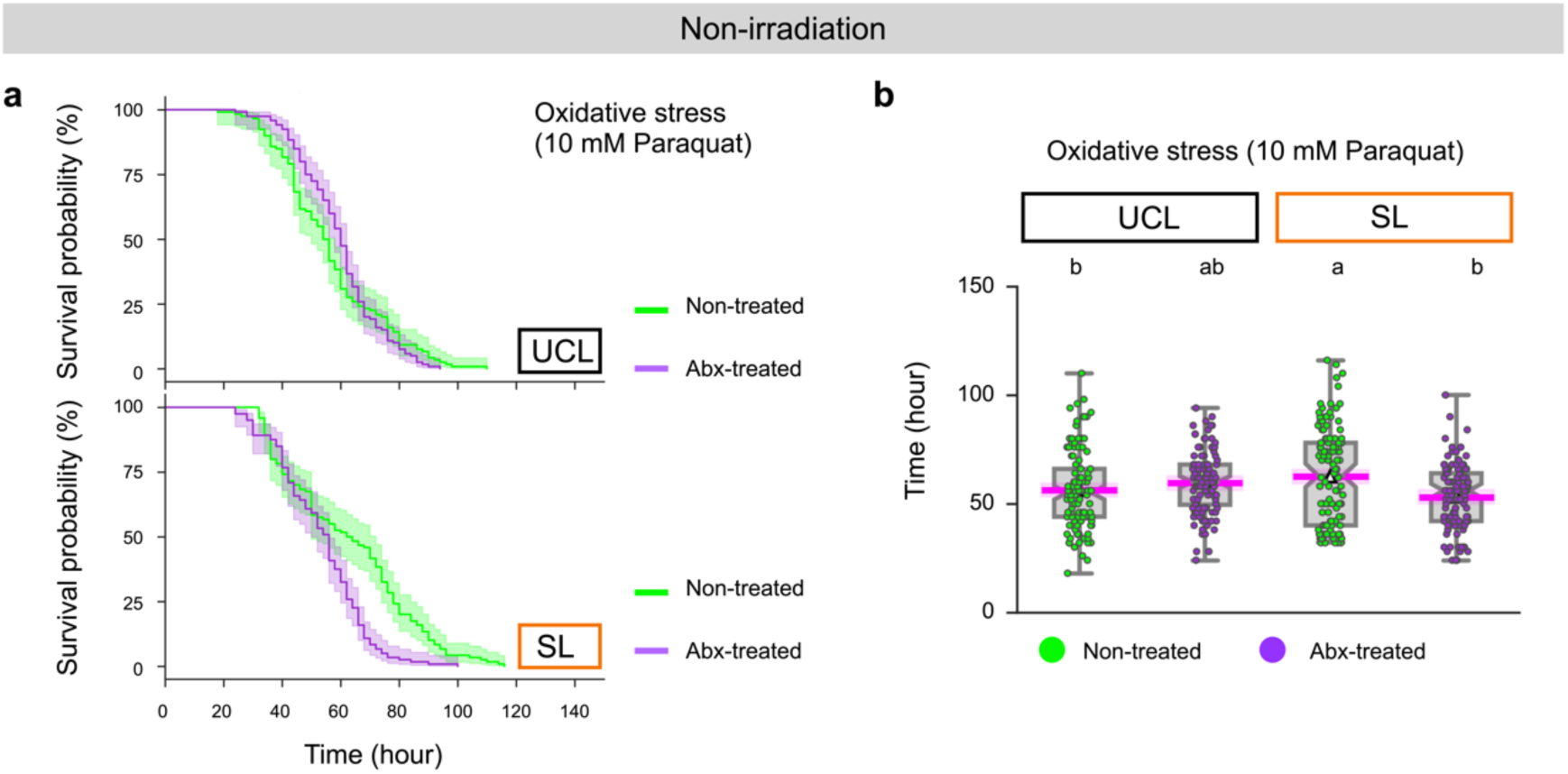
Resistance to oxidative stress in SL flies depends on gut microbiota. **a**, Oxidative stress resistance in SL flies treated with 10 mM paraquat. Kaplan–Meier survival curve, *n* = 120; replicate line 1, generation 52. **b**, Oxidative stress resistance in SL and UCL flies treated with 10 mM paraquat. *n* = 120, BF10 = 0.01. Survival: non-treated UCL 56.36 h (53.29–59.43), Abx-treated UCL 59.57 h (56.33–62.61), non-treated SL 62.48 h (59.42–65.64), Abx-treated SL 53.10 h (50.02–56.22); replicate line 1, generation 52. Statistical analysis was performed using HBM with MCMC methods. The magenta, bold lines and shaded areas show the posterior median and the Bayesian 95% CI. Different letters indicate significant differences at *p*MCMC < 0.05. See Supplementary Data 7 for details of all results, including statistical models.

**Supplementary Fig. 12:**
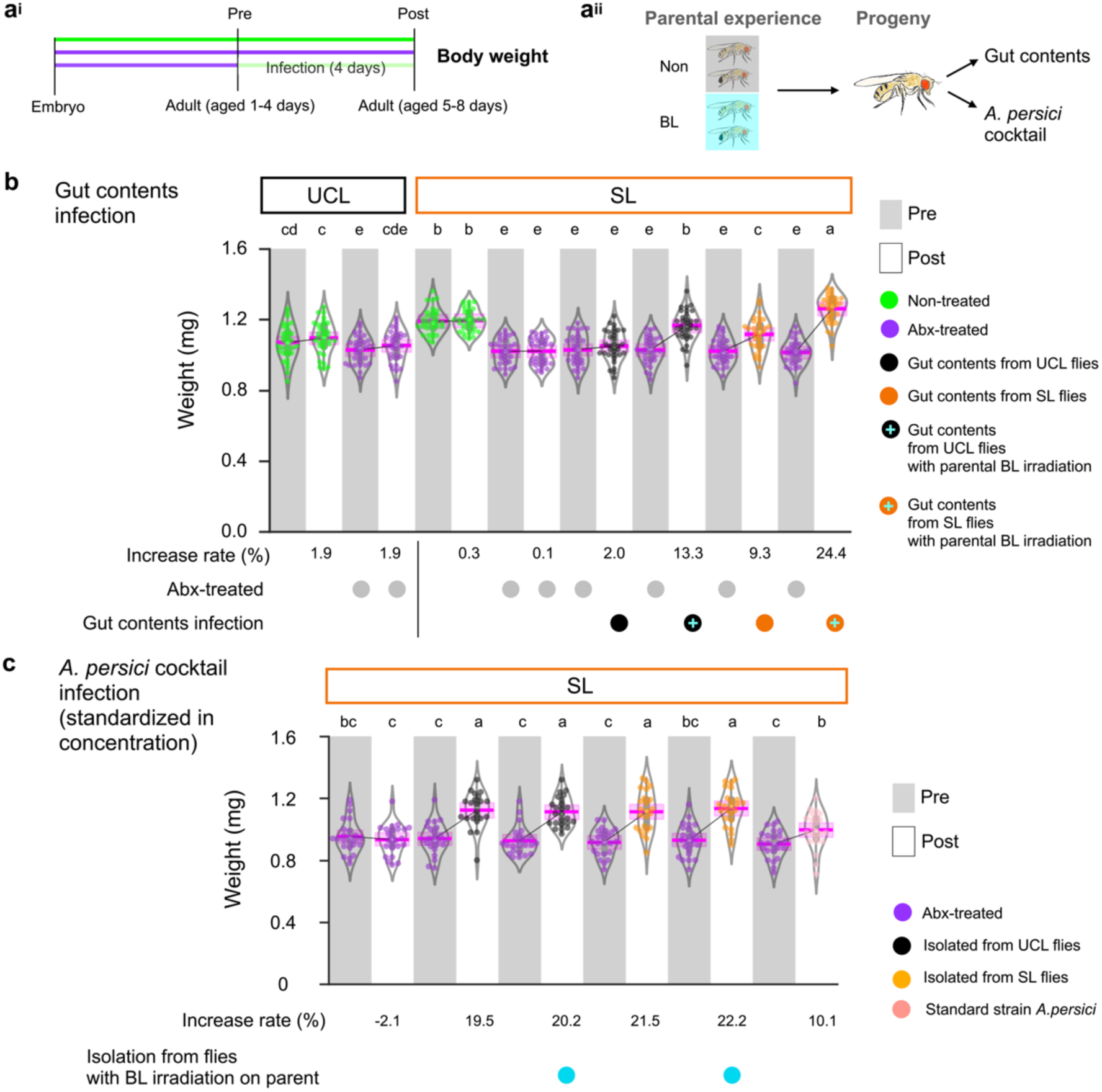
**Greater bacterial abundance leads to higher body weight. a^i^**, Schematic of the infection experiment. **a^ii^**, Overview of gut contents or isolated *Acetobacter persici* cocktail from offspring of parents exposed to BL. **b**, Impact of infection with gut contents on body weight. *n* = 30. **c**, Impact of infection with *A. persici* cocktail on body weight. *n* = 23, 24. In both panels, grey bars represent pre-infection; white bars indicate post-infection after 4 days. BF10 >100; replicate line 1, generation 53. Statistical analyses were performed using HBM with MCMC methods. The magenta, bold lines and shaded areas show the posterior median and the Bayesian 95% CI. Different letters indicate significant differences at *p*MCMC < 0.05. See Supplementary Data 8 for details of all results, including statistical models.

**Supplementary Fig. 13:**
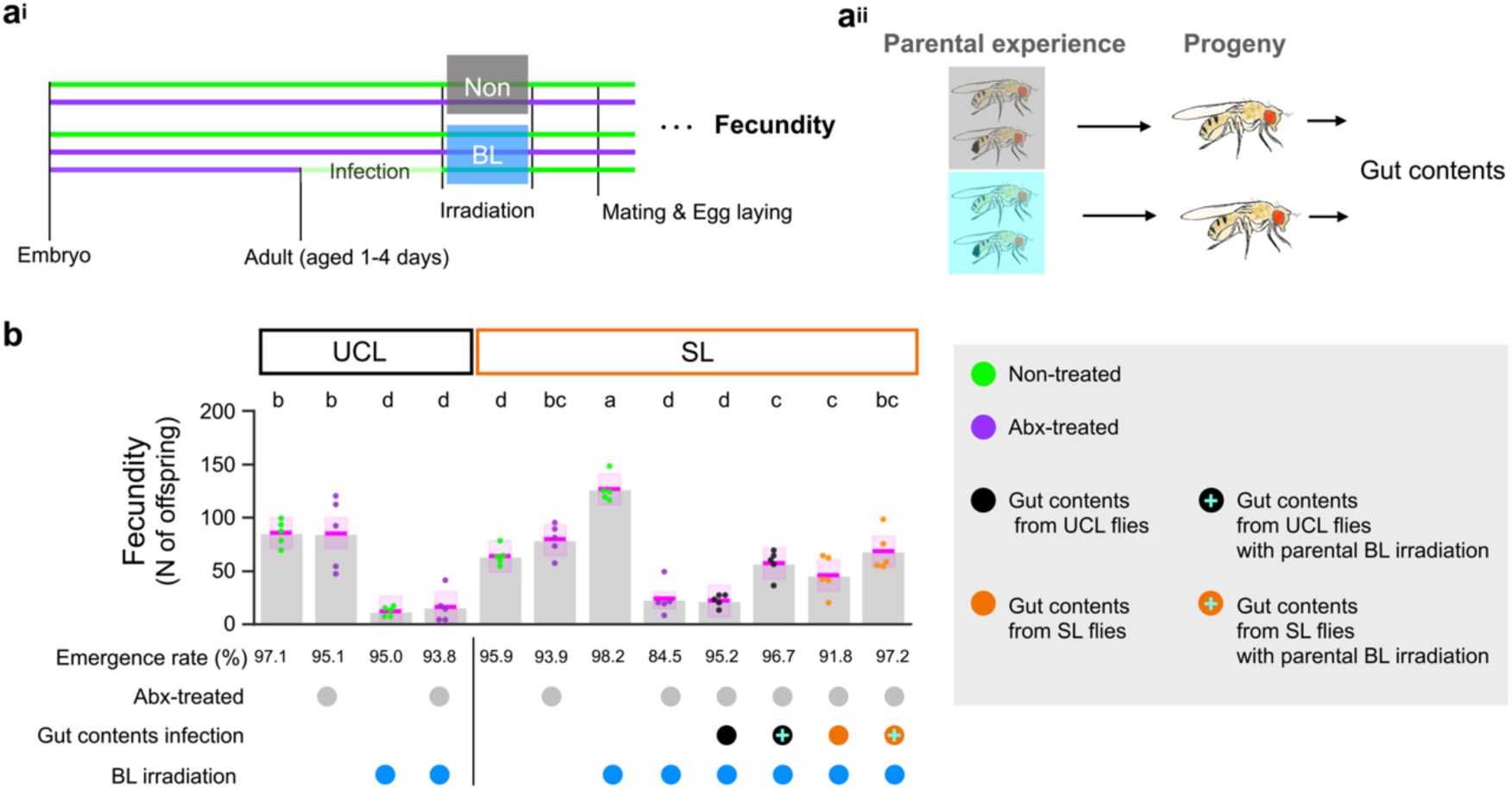
Gut contents from BL-exposed UCL parental flies restore fecundity in Abx-treated SL flies. **a^i^**, Schematic of the infection experiment. **a^ii^**, Overview of gut contents from offspring of parents exposed to BL. **b**, Fecundity: number of offspring (adults). *n* = 5. BF10 >100; replicate line 1, generation 53. Statistical analysis was performed using HBM with MCMC methods. The magenta, bold lines and shaded areas show the posterior median and the Bayesian 95% CI. Different letters indicate significant differences at *p*MCMC < 0.05. See Supplementary Data 8 for details of all results, including statistical models.

**Supplementary Fig. 14:**
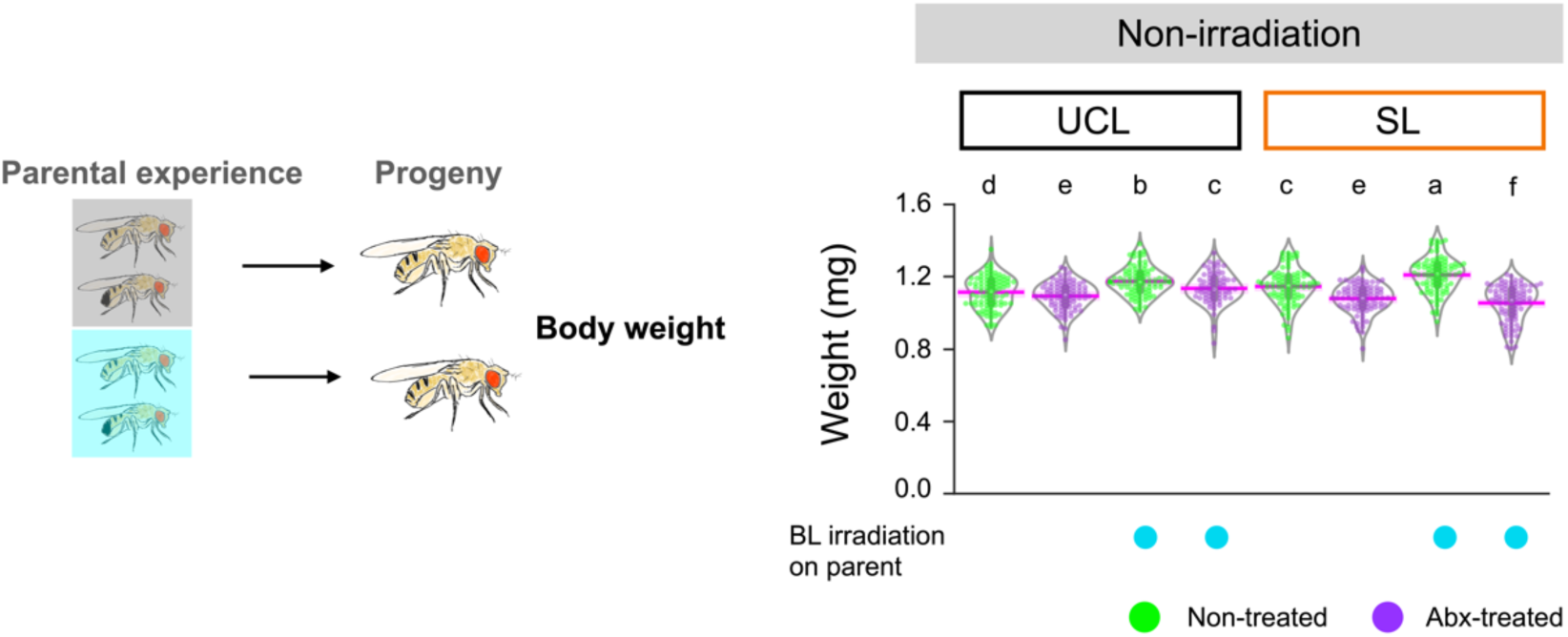
Parental BL experience increases body weight in progeny. Overview of the offspring of parents exposed to BL. Data is based on female body weight. *N* = 72–90, BF10 > 100; replicate line 1, generation 46. Statistical analysis was performed using HBM with MCMC methods. The magenta, bold lines and shaded areas show the posterior median and the Bayesian 95% CI. Different letters indicate significant differences at *p*MCMC < 0.05. See Supplementary Data 8 for details of all results, including statistical models.

**Supplementary Fig. 15:**
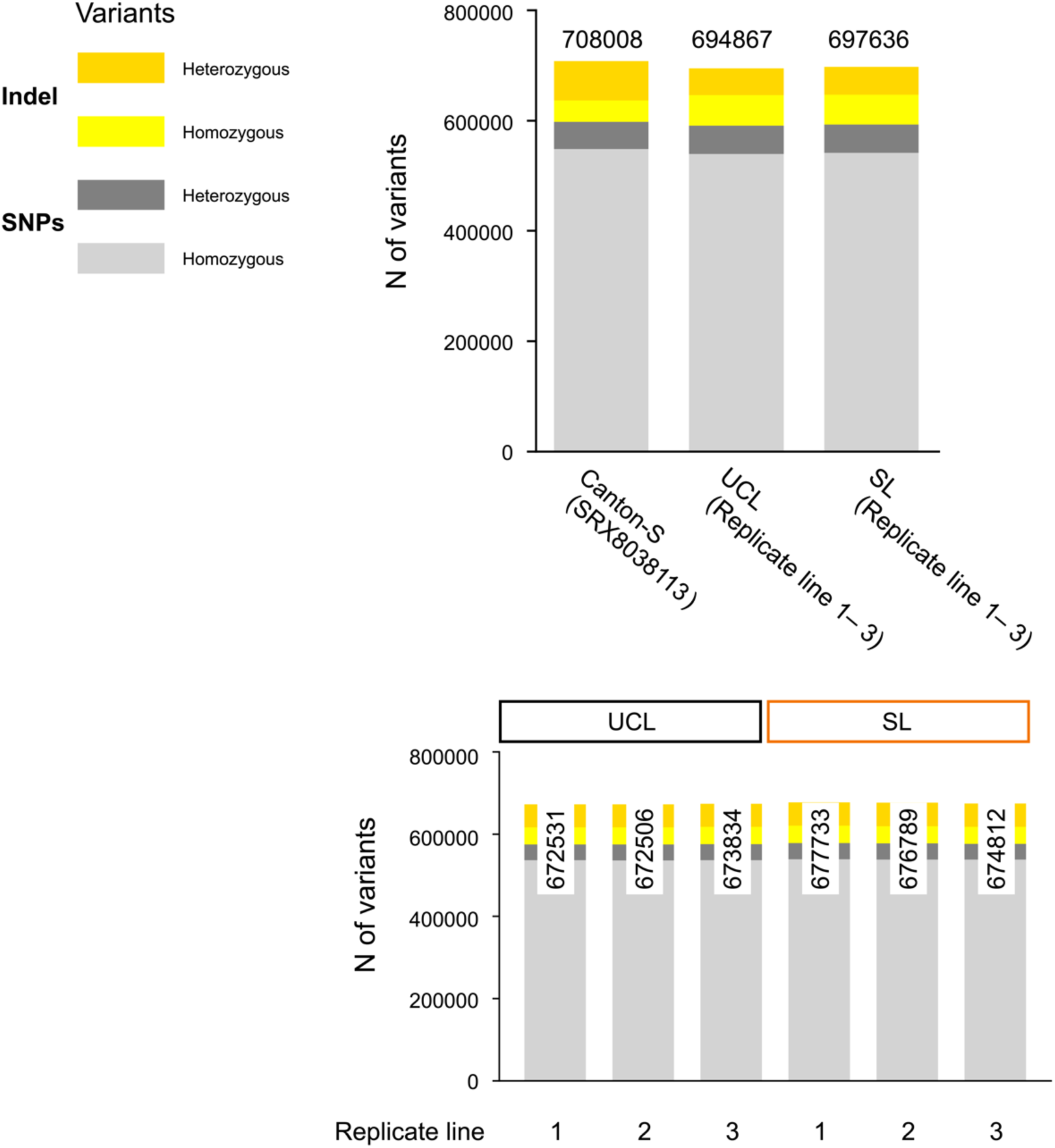
Number of variants potentially present in the strains used for the selection experiment. Number of variants in the Canton-S strain (SRX8038113), UCL and SL. Variants in UCL and SL are the total number of variants from replicate lines 1 to 3. The bottom panel shows the number of variants for each replicate line. Colour differences indicate SNPs and indels as well as homozygous and heterozygous variants. The numbers on or within the bars represent the total number of variants.

**Supplementary Fig. 16:**
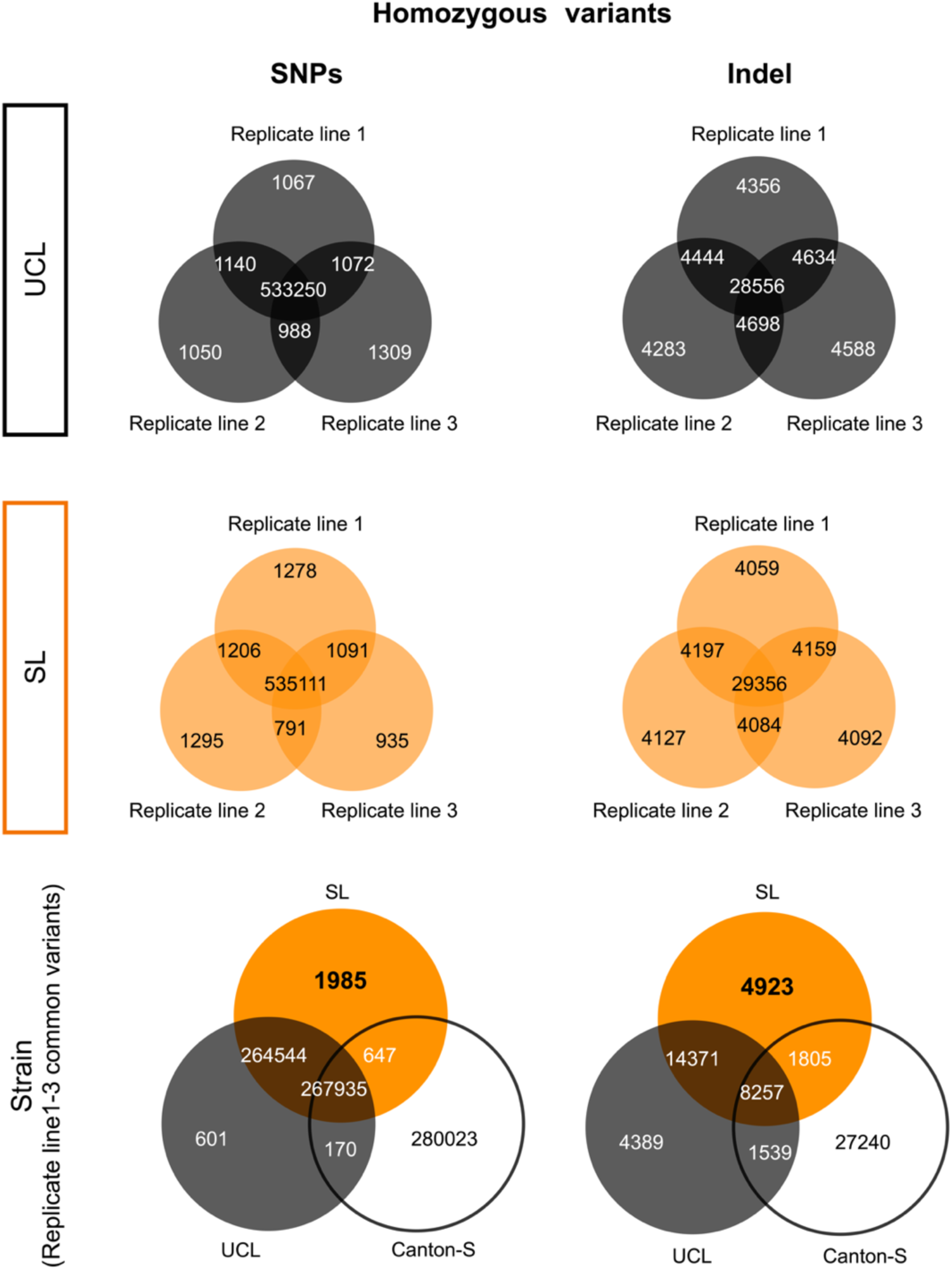
Number of homozygous variants specific to SL and shared by all three replicate lines, potentially selected under BL toxicity. Common and unique variants identified in each replicate line across different strains. Variants shared within a strain were considered strain-specific, whereas homozygous variants absent in both UCL and Canton-S were defined as SL-specific.

**Supplementary Fig. 17:**
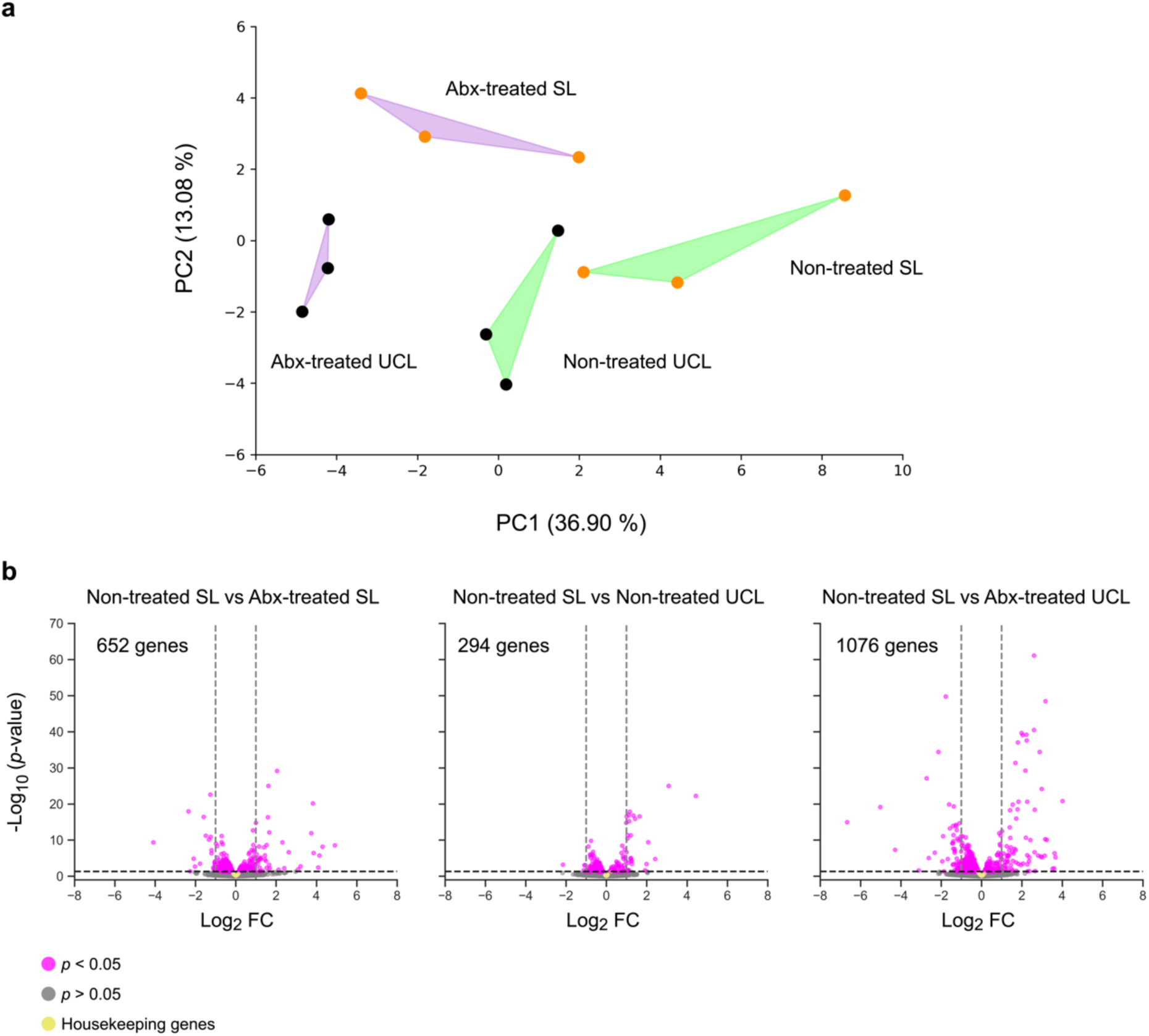
Overview of the host transcriptome profiles. **a**, Principal component analysis (PCA) of transcriptome profiles using normalised read counts. **b**, Volcano plots comparing non-treated SL with each condition. Differentially expressed genes (DEGs) with a false discovery rate (FDR)-adjusted *p*-value threshold of <0.05 are shown in magenta. Housekeeping genes were defined based on De Ferrari and Aitken, 2006, *BMC Genomics*^1^. For details, see Supplementary Data 13.

**Supplementary Fig. 18:**
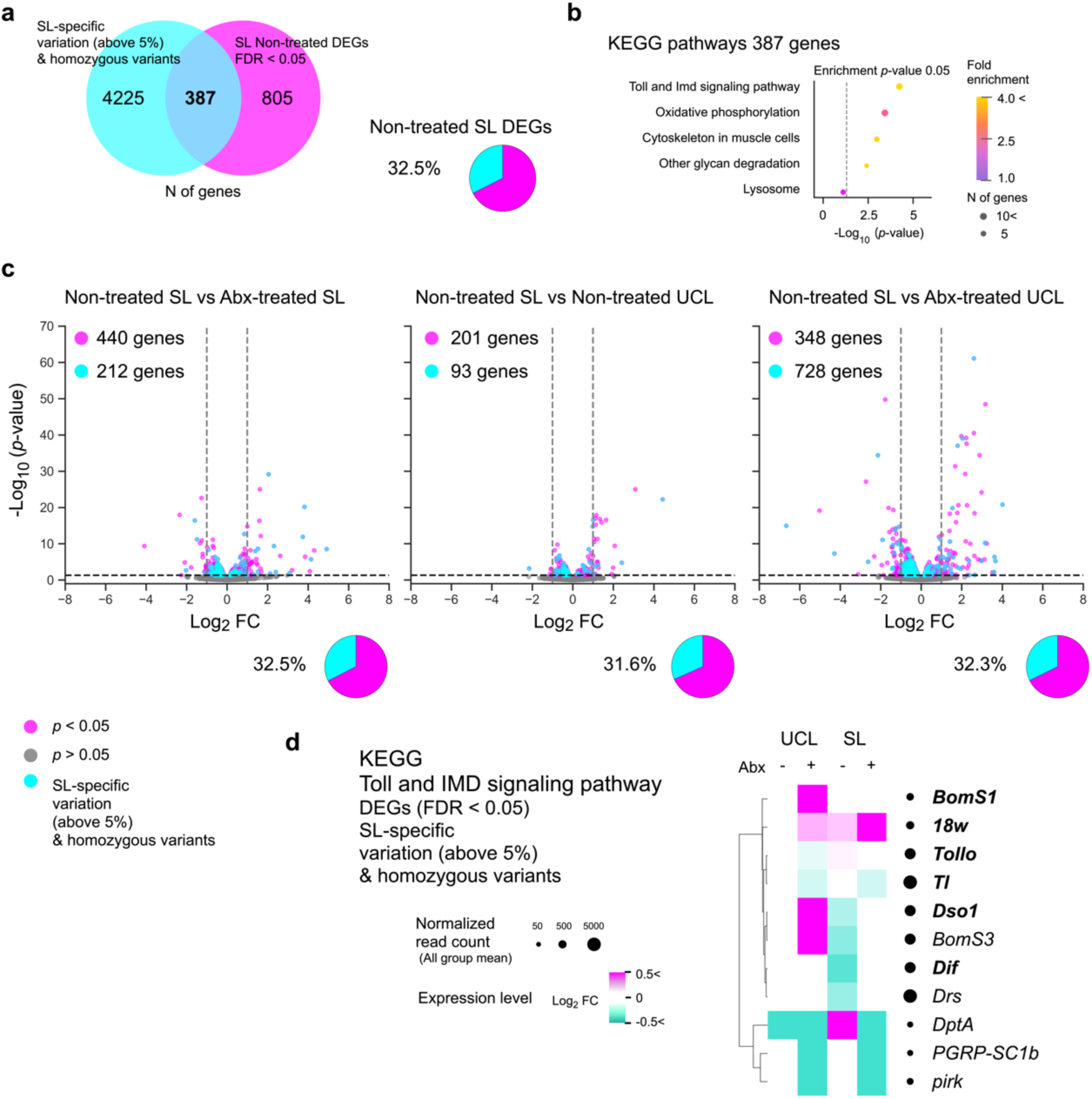
Genes shared between genetic characteristics unique to SL and non-treated SL DEGs are enriched in immune-related pathways. **a**, Venn diagram of genetic characteristics unique to SL and differentially expressed genes (DEGs) in non-treated SL, showing a 32.5% overlap. **b**, Enrichment analysis of 387 overlapping genes using the Kyoto Encyclopedia of Genes and Genomes (KEGG), with the top hit identified as the Toll and Imd signaling pathway. **c**, Volcano plots comparing non-treated SL with each condition, highlighting genetic characteristics unique to SL. Approximately 32% overlap was observed across all treatments. **d**, Expression levels of Toll and IMD signaling pathway genes shared between genetic characteristics unique to SL and non-treated SL DEGs. Expression levels are presented as mean log2 FC for each group, calculated by first calculating log2 FC for individual samples relative to the average expression in non-treated UCL and then averaging across groups. Circles indicate mean read counts across all groups. Homozygous variants are indicated in bold. For details, see Supplementary Data 14.

**Supplementary Fig. 19:**
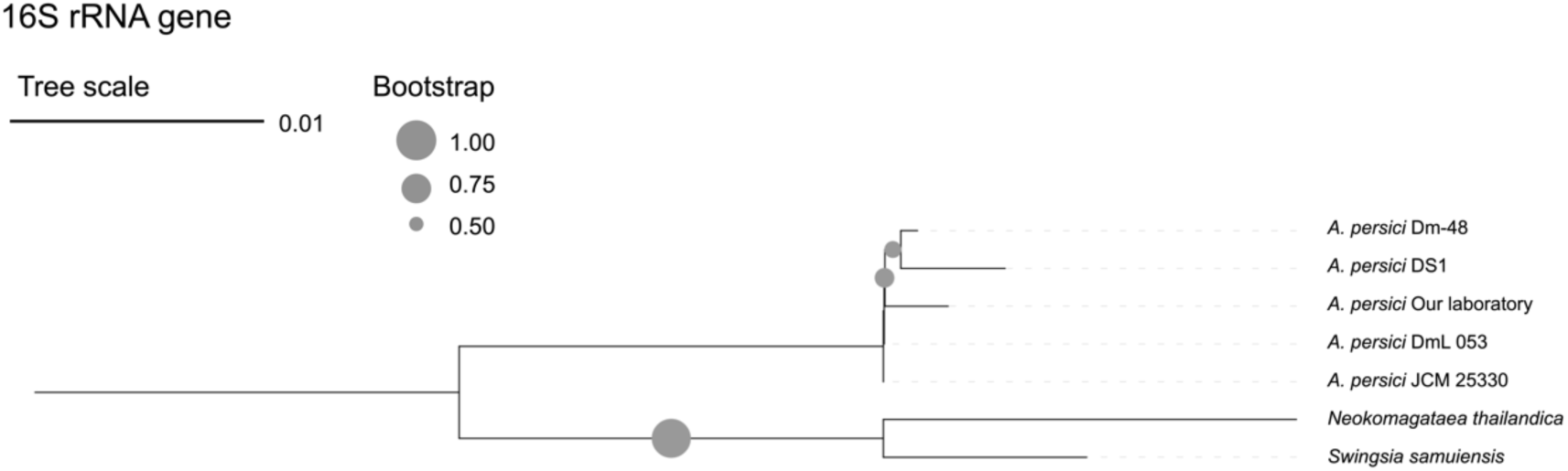
Phylogenetic tree based on the 16S rRNA gene of *A. persici* Phylogenetic relationship between the dominant OTU ID (397 bp), which comprised 89-98% of *A. persici* in our laboratory, and other *A. persici* strains. Other bacterial species were analysed using full-length sequences (1,391-1,498 bp). *Neokomagataea thailandica* and *Swingsia samuiensi*s were used as outgroups. The tree was constructed using maximum likelihood with 1,000 bootstrap replicates.

**Supplementary Fig. 20:**
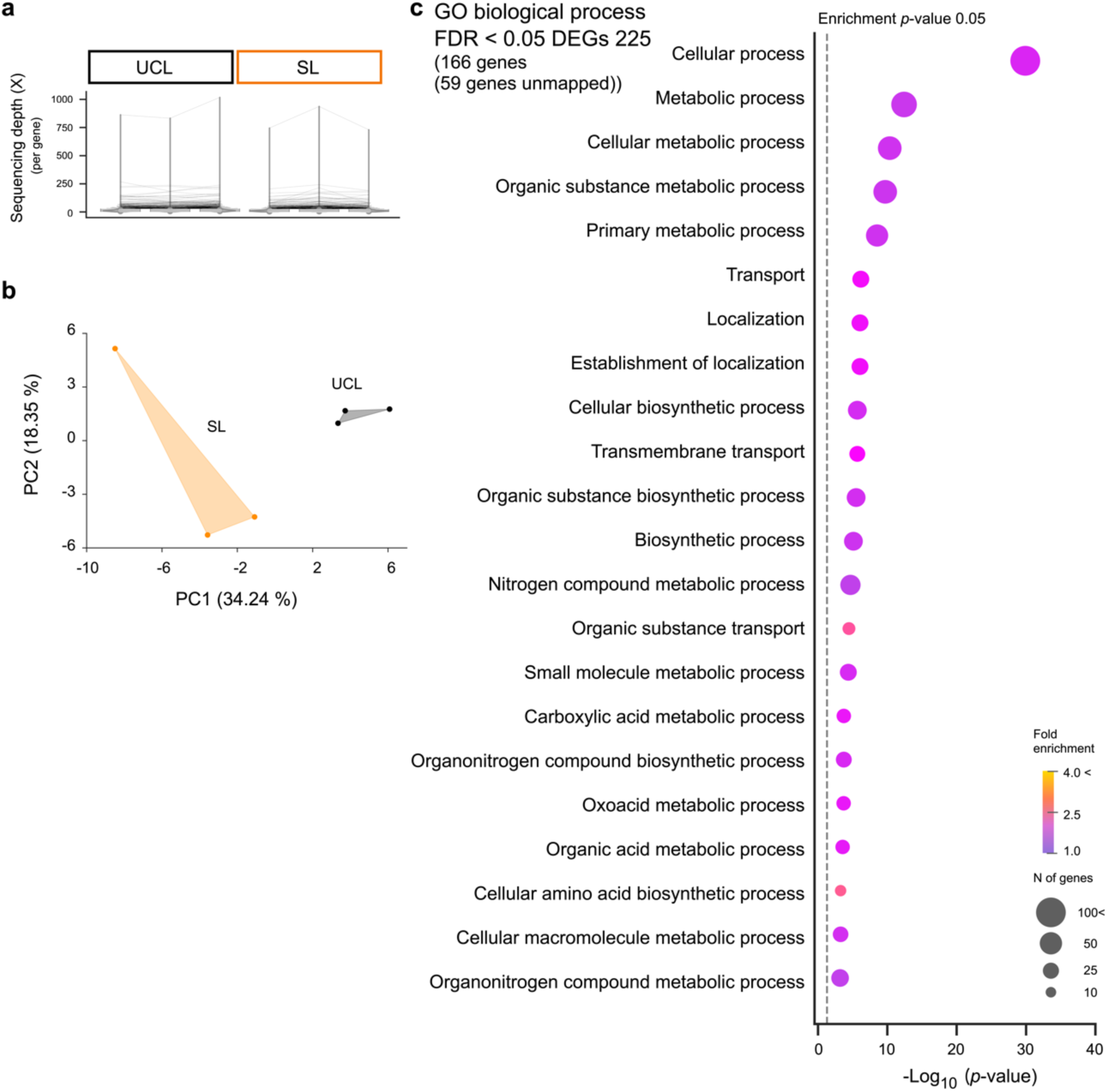
Transcriptome profile of the gut microbiota *A. persici* **a**, Sequence depth based on mapping results. After extracting unmapped reads from the *Drosophila melanogaster* host (GCA_000001215.4), the dominant gut bacteria *A. persici* (ASM200656v1) was used as a reference for mapping. The same genes are connected by lines. The gene with the highest sequencing depth across all samples was *Large ribosomal subunit protein bL12*. See Fig 10c for magnification and mean values. **b**, PCA based on transcripts per million (TPM). **c**, Enrichment analysis of 225 DEGs (Gene Ontology (GO); biological process), predominantly indicative of metabolic pathways. GO enrichment analysis was then performed on 166 genes mapped by DEGs to the STRING database. For details, see Supplementary Data 16.

**Supplementary Fig. 21:**
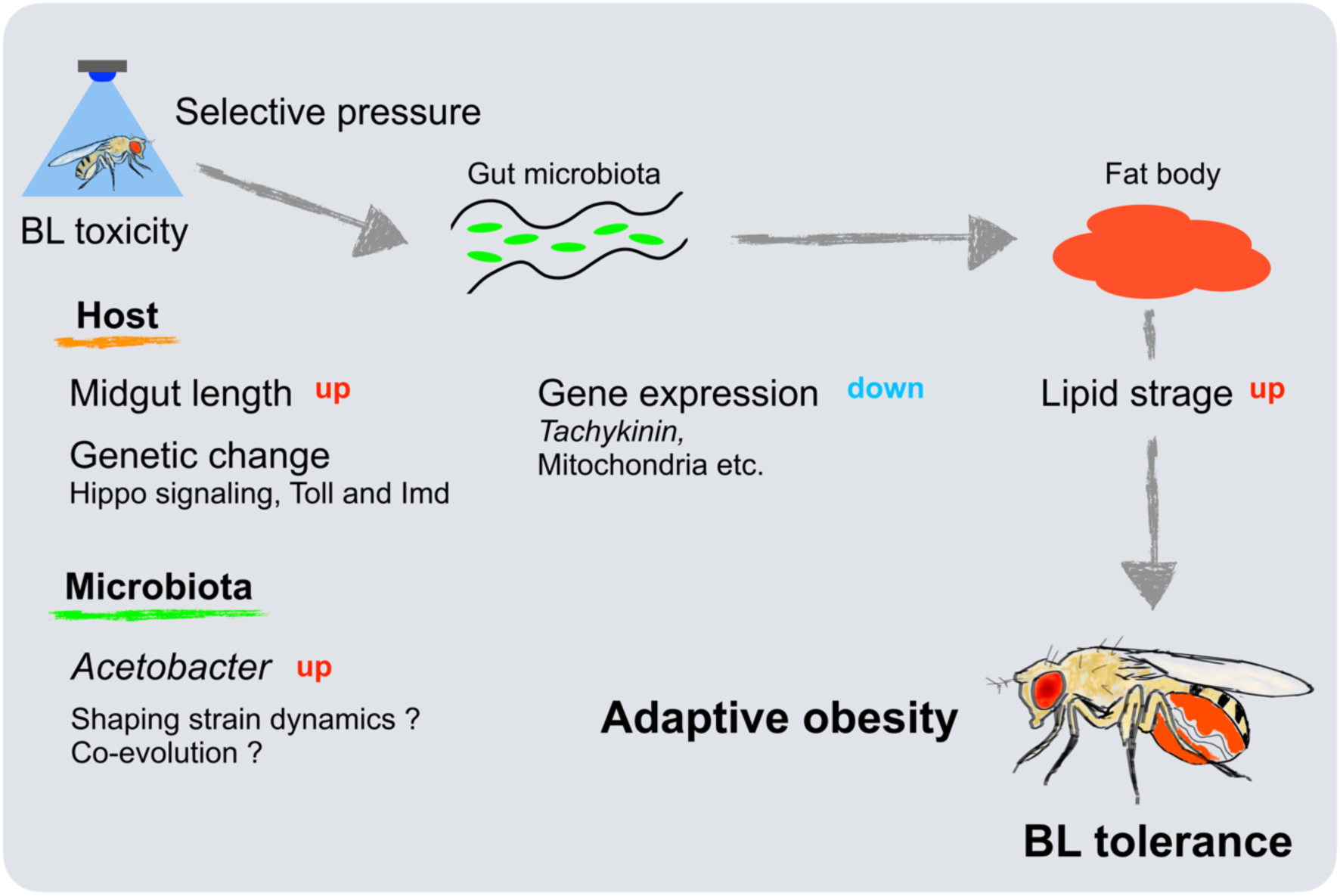
Gut microbiota–mediated obesity drives evolutionary adaptation to BL toxicity. Evolutionary adaptation to BL toxicity involves selective pressure and the intergenerational transmission of parental experience, which drives midgut elongation, increased bacterial abundance, and optimisation of both host genetic information and microbiota interaction. The shaping of bacterial strain dynamics or bacterial evolution may occur in parallel with host evolution. This process may be associated with increases in bacterial abundance and phenotypic changes, suggesting that both quantitative and qualitative changes in bacterial traits may alter host interactions. These changes can alter gene expression related to lipid metabolism, facilitating the formation of adaptive obesity characterised by lipid accumulation via the gut microbiota.

